# Genome-wide association study on 13,167 individuals identifies regulators of hematopoietic stem and progenitor cell levels in human blood

**DOI:** 10.1101/2021.03.31.437808

**Authors:** Aitzkoa Lopez de Lapuente Portilla, Ludvig Ekdahl, Caterina Cafaro, Zain Ali, Natsumi Miharada, Gudmar Thorleifsson, Kristijonas Žemaitis, Antton Lamarca Arrizabalaga, Malte Thodberg, Maroulio Pertesi, Parashar Dhapola, Erik Bao, Abhishek Niroula, Divya Bali, Gudmundur Norddahl, Nerea Ugidos Damboriena, Vijay G. Sankaran, Göran Karlsson, Unnur Thorsteinsdottir, Jonas Larsson, Kari Stefansson, Björn Nilsson

## Abstract

Understanding how hematopoietic stem and progenitor cells (HSPCs) are regulated is of central importance for the development of new therapies for blood disorders and stem cell transplantation. To date, HSPC regulation has been extensively studied *in vitro* and in animal models, but less is known about the mechanisms *in vivo* in humans. Here, in a genome-wide association study on 13,167 individuals, we identify 9 significant and 2 suggestive DNA sequence variants that influence HSPC (CD34^+^) levels in human blood. The identified loci associate with blood disorders, harbor known and novel HSPC genes, and affect gene expression in HSPCs. Interestingly, our strongest association maps to the *PPM1H* gene, encoding an evolutionarily conserved serine/threonine phosphatase never previously implicated in stem cell biology. *PPM1H* is expressed in HSPCs, and the allele that confers higher blood CD34^+^ cell levels downregulates *PPM1H*. By functional fine-mapping, we find that this downregulation is caused by the variant rs772557-A, which abrogates a MYB transcription factor binding site in *PPM1H* intron 1 that is active in specific HSPC subpopulations, including hematopoietic stem cells, and interacts with the promoter by chromatin looping. Furthermore, rs772557-A selectively increases HSPC subpopulations in which the MYB site is active, and *PPM1H* shRNA-knockdown increased CD34^+^ and CD34^+^90^+^ cell proportions in umbilical cord blood cultures. Our findings represent the first large-scale association study on a stem cell trait, illuminating HSPC regulation *in vivo* in humans, and identifying *PPM1H* as a novel inhibition target that can potentially be utilized clinically to facilitate stem cell harvesting for transplantation.

## Main text

Humans blood cells originate from HSPCs^1^. Although HSPCs primarily reside in the bone marrow, they continuously egress into the blood, where they constitute a tiny (∼0.1%) subset of mononuclear white blood cells. Circulating HSPCs express the surface protein CD34, allowing quantification by flow cytometry (**Supplementary Fig. 1 and 2**)^2^. Epidemiological studies^3, 4^ show that the level of circulating CD34^+^ cells varies between individuals, but remains relatively stable within individuals, pointing at a genetic component. However, the underlying genes and DNA sequence variants remain unknown^3^.

From a clinical viewpoint, novel HSPC regulators are highly sought-after to improve the treatment of blood disorders. Firstly, stem cell transplantation is a cornerstone in the treatment of blood cell malignancies. Today, the preferred way of harvesting stem cells for transplantation is by leukapheresis of peripheral blood. This requires that the CD34^+^ cell level in the donor’s blood is sufficiently high, and existing ways to mobilize CD34^+^ cells from the bone marrow into the blood are not always effective^5, 6^. Finding genes that regulate CD34^+^ levels can lead to new ways to mobilize HPSCs. Secondly, several hematologic malignancies, including acute myeloid leukemia (AML)^7, 8^, myelodysplastic syndrome (MDS)^9, 10^, and myeloproliferative neoplasms (MPN)^11^ are caused by abnormal HSPC activity. Uncovering the regulatory circuits that control HSPCs can lead to novel anti-leukemic therapies.

### Genome-wide association study

To search for regulators of blood CD34^+^ cell levels *in vivo* in humans, we carried out a genome-wide association study (GWAS) in 13,167 individuals from southern Sweden (ages 18 to 71 years; **Supplementary Table 1** and **Supplementary Fig. 3**). To quantify blood CD34^+^ cells at a large scale, we developed a high-throughput flow-cytometry workflow, and pattern recognition software for computer-assisted analysis of the large volumes of flow-cytometry data (**Online Methods**). In each sample, we analyzed up to 1 million white blood cells (**Supplementary Table 2**), and defined the CD34^+^ cell level as the number of CD34^+^ cells divided by the number of CD45^+^ mononuclear cells (**Supplementary Fig. 1**). To assess reproducibility, we sampled 660 individuals twice, with 3 to 36 months between samplings, and found a strongly significant correlation between replicates (Spearman *P* = 1.3×10^-84^, *r*^2^ = 0.44; **Supplementary Fig. 4**). We observed higher blood CD34^+^ cell levels in males than in females, but no differences between age groups (**Supplementary Fig. 5**).

Participants were genotyped on single-nucleotide polymorphism (SNP) microarrays, and imputed with reference whole-genome sequencing data to a final resolution of 18 million SNPs and small insertions-deletions (INDELs). For association analysis, we split our data into a discovery set of 10,949 individuals of Swedish ancestry and a follow-up set of 2,218 individuals of non-Swedish European ancestry using principal component analysis of the genotype data. To correct for multiple testing, we partitioned variants into five classes based on genomic annotations and applied weighted Bonferroni adjustment, taking into account the predicted functional impact of variants within each class (**Online Methods**)^12^.

In combined analysis of the two data sets, 6 loci reached significance and 2 were suggestive (within one order of magnitude from Bonferroni thresholds). Conditional analysis uncovered four independent signals at 2q22, and no underlying signals at the other loci (**Fig. 1)**. No significant heterogeneity was detected in the effect estimates between the discovery and follow-up set (**Supplementary Table 3**). The variance explained by the 9 significant variants was 4.6%. Using linkage disequilbrium (LD) score regression, we estimated the total heritability at 12.7%, which is comparable to other blood cell traits (*e.g.*, mature white blood cell counts)^13–16^. Moreover, we detected enrichments of heritability in genomic regions with accessible chromatin in HSPC subpopulations^17, 18^ (**Fig. 2a**), including hematopoietic stem cells (HSC), multi-potent progenitors (MPP), and common myeloid progenitors (CMP), indicating that our analysis preferentially identifies variants that act by altering the regulation of gene expression intrinsically in HSPCs.

**Figure 1:**
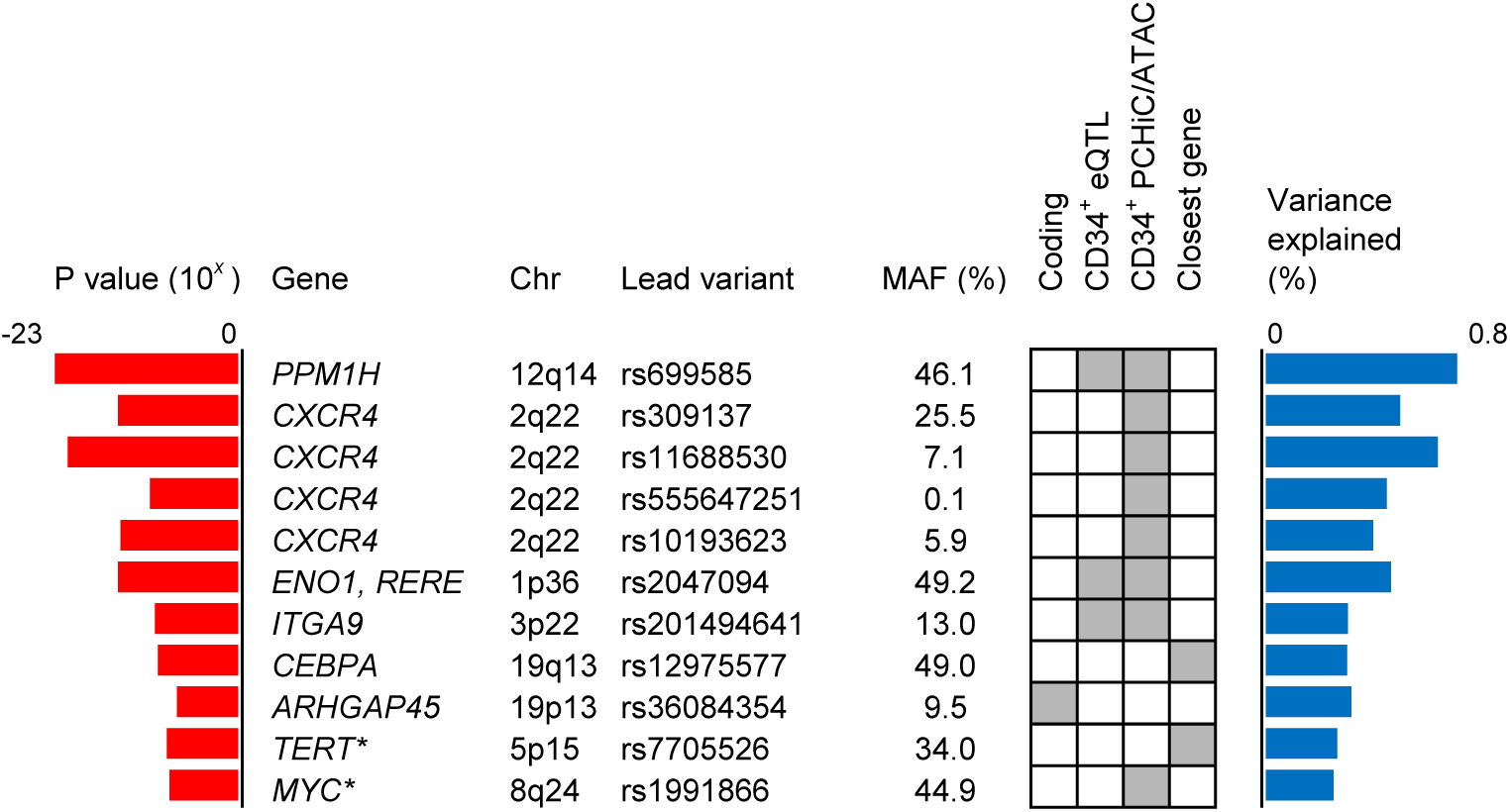
Genome-wide association study. Sequence variants influencing blood CD34^+^ cell levels identified in combined analysis of association data for 10,949 individuals of Swedish ancestry and 2,218 individuals of non-Swedish European ancestry. We identified 9 significant and 2 suggestive (*) associations (**Supplementary Table 3**). The listed variants are the most significant (lead) variants for each association. We prioritized genes as candidate genes if they: (i) had a coding variant within the 99% credible set of probable causal variants (**Supplementary Table 4**); (ii) had a *cis*-eQTL in CD34^+^ cells from 155 blood donors (**Supplementary Table 5**); or (iii) the credible set contained a regulatory variant that maps either to the promoter, or to a region with a chromatin looping interaction with the promoter in CD34^+^ cells, as determined by PCHi-C. As regulatory variants, we considered variants in genomic regions whose chromatin is accessible in HSPCs, as determined by ATAC- sequencing. If none of these criteria were fulfilled, we prioritized the closest gene. The criterion used to call each gene a candidate gene is indicated in the matrix.

**Figure 2:**
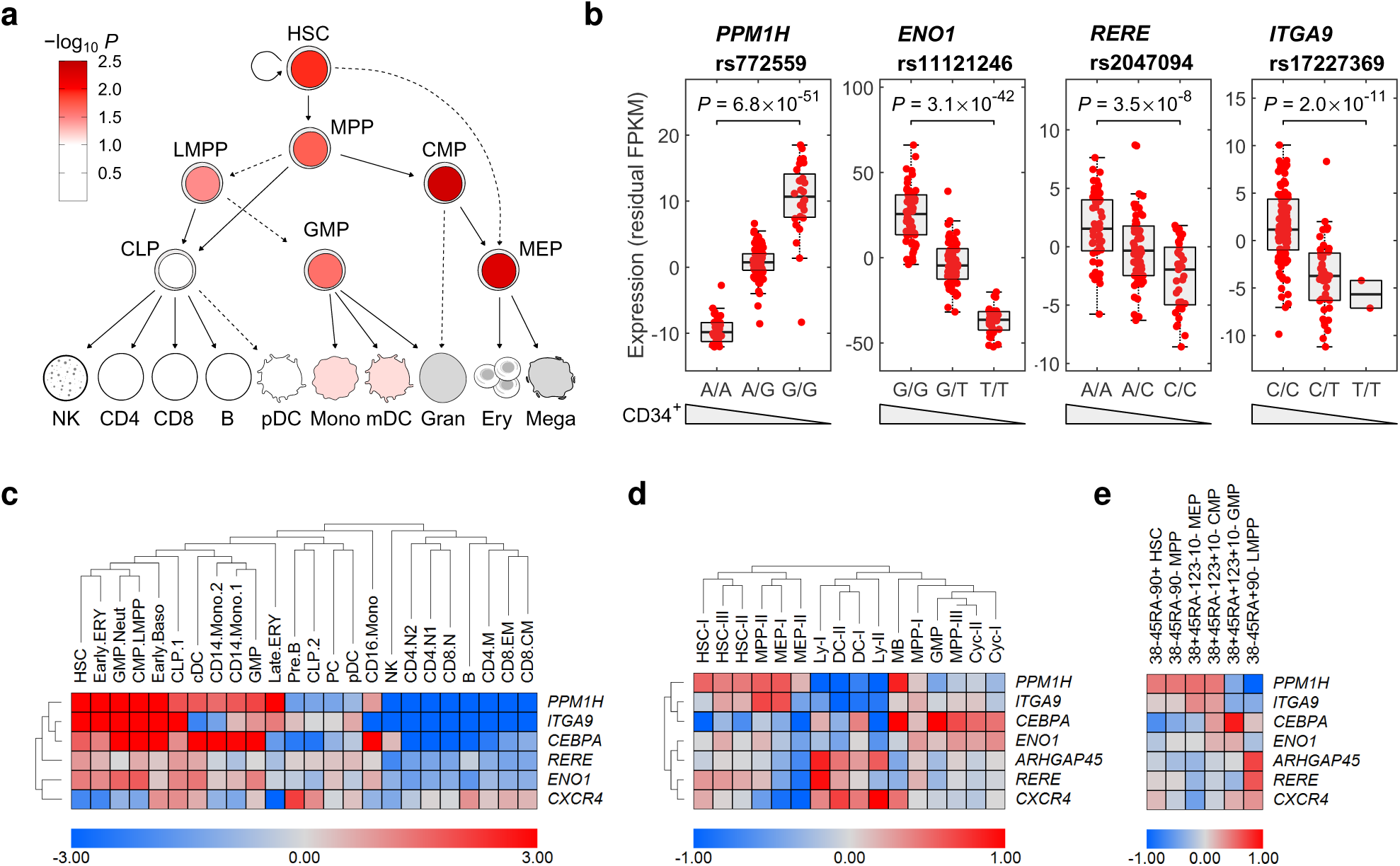
Effects on gene expression in HSPCs. **(a)** LD score regression shows enrichments of heritability in regions with accessible chromatin in HSPC subpopulations. **(b)** To identify candidate genes with HSPC-intrinsic gene-regulatory effects, we generated eQTL data for sorted CD34^+^ cells from 155 blood donors. These figures illustrate strong *cis*-eQTLs identified at *PPM1H*, *ENO1, RERE* and *ITGA9* (**Supplementary Table 5**). Data are residual FPKM values after correction for 10 expression principal components. Wedges indicate directions of effects on blood CD34^+^ cell levels for the same variant. Notably, we detected an anti-correlation between *PPM1H* expression and CD34^+^ levels for the 12q14 variant. **(c)** Candidate gene expression in scRNA-seq data from blood and bone marrow mononuclear cells^28^, showing enriched expression in HSPC populations (**Supplementary Fig. 7** and **8**). **(c,d)** To map expression within the CD34^+^ compartment in better detail, we analyzed CITE-seq (*i.e.*, mRNA-sequencing with antibody-derived tags) data for 4,905 lineage-negative CD34^+^ cells from adult bone marrow: **(d)** bulked expression in cell clusters inferred from mRNA levels; **(e)** bulked expression in clusters inferred from antibody-derived tags representing classical HSPC surface markers (**Supplementary Fig. 9** and **10**). Abbreviations: Hematopoietic stem cells (HSC), multi-potent progenitors (MPP), common myeloid progenitors (CMP), granulocyte-monocyte progenitors (GMP), common lymphoid progenitors CLP), lymphoid-primed multipotent progenitors (LMPP), erythroid progenitors (ERY), megakaryocyte-erythrocyte progenitors (MEP), mast cell/basophil progenitors, (MB), dendritic cells (DC), plasma cells (PC), CD4^+^ T-cells (CD4), CD8^+^ T-cells (CD8), B-cells (B), pre B-cells (PreB), lymphoid progenitors (Ly), natural killer cells (NK), basophil (Baso), neutrophil (Neut), monocyte (Mono), cycling cells (Cyc).

### Identification of candidate genes

To identify candidate genes based on HSPC-intrinsic gene-regulatory effects, we generated expression quantitative locus (eQTL) data for sorted CD34^+^ cells from 155 blood donors by mRNA-sequencing (average 122,000 cells per sample). Additionally, to identify regulatory links between variants and genes that are not detectable at the mRNA level in peripheral blood, we retrieved promoter capture Hi-C (PCHi-C) data^16^ for CD34^+^ cells and ATAC-sequencing data for 19 sorted blood cell populations, including 7 HSPC populations^11^. We then defined the 99% credible sets of probable causal variants (**Supplementary Table 4**), and prioritized genes as candidate genes if they: (i) had a non-synonymous coding variant within the credible set, (ii) had a *cis*-eQTL in CD34^+^ cells (**Fig. 2b** and **Supplementary Table 5**), or (iii) the credible set contained a regulatory variant that maps either to the promoter region, or to a region with a chromatin looping interaction with the promoter in CD34^+^ cells, as determined by PCHi-C^19^ (**Supplementary Fig. 6**). As regulatory variants, we considered variants in genomic regions with accessible chromatin in HSPCs, as determined by ATAC-sequencing^13^ (**Supplementary Table 4**). If none of these criteria were fulfilled, we prioritized the closest gene. Using these criteria, we identified 7 candidate genes at the significant loci, including two known HSPC-relevant genes (*CXCR4, CEBPA*) and five genes that have never previously been implicated in hematopoietic stem cell biology (*PPM1H, ENO1, RERE, ARHGAP45)* or only studied minimally in this area (*ITGA9*)^20^ (**Fig. 1**).

### Genetic overlap with blood disorders and other blood cell traits

To investigate further their impact, we asked if the variants that influence blood CD34^+^ cell levels also associate with other traits and diseases. To address this, we searched for coincident associations among variants in high LD (*r*^2^ > 0.8) with the lead variants using Phenoscanner^21, 22^. Consistent with HSPCs producing blood and immune cells, we detected coincident associations with variants known to influence mature blood cell traits (6 significant signals, at 12q14/*PPM1H,* 1p36/*ENO1-RERE*, 19p13/*ARHGAP45*, 19q13/*CEBPA* and two signals at 2q22/*CXCR4*), as well as with hematologic malignancies and autoimmune disorders (2 signals, at 2p22/*CXCR4* and 19p13/*CEBPA*) (**Supplementary Table 6**).

Of note, some of the candidate genes underlie autosomal-dominant blood disorders characterized by aberrant HSPC function. Gain-of-function mutations in *CXCR4* cause the Warts, Hypogammaglobulinemia, Immunodeficiency and Myelokathexis (WHIM) syndrome, marked by impaired egression of HSPCs and other white blood cells out of the bone marrow^23, 24^. *CEBPA* mutations cause familial acute myeloid leukemia^25^. *TERT* mutations cause dyskeratosis congenital, where impaired telomere maintenance leads to problems with HSPC regeneration and increased risk of MDS^26^. Furthermore, somatic acquired mutations in *CXCR4*^27^*, CEBPA*^9, 25^, and *TERT*^11^ have been reported in several hematologic malignancies.

### Gene expression in human hematopoiesis

We next explored the expression of the candidate genes across hematopoietic cell types. First, in single-cell mRNA-sequencing data for 35,582 mononuclear blood cells from adult blood and bone marrow^28^, we observed significant enrichment of expression in HSPC *vs* non-HSPC populations (Wilcoxon *P* = 2.1**×**10**^-^**^10^ for the candidate genes at the significant loci; **Fig. 2c** and **Supplementary Fig. 7**). The same observation was made in bulk mRNA-sequencing data for sorted blood cell populations (Wilcoxon *P* = 7.1**×**10^-5^; **Supplementary Fig. 8**).

To map gene expression within the CD34^+^ compartment in better detail, we analyzed single-cell Cellular Indexing of Transcriptomes and Epitopes by sequencing (CITE-seq) data for 4,905 lineage-negative CD34^+^ adult bone marrow cells^29^. This revealed distinct expression biases for *PPM1H*, *ITGA9, ENO1* and *RERE* (HSC, MPP, CMP and MEP bias), *CXCR4* and *ARHGAP45* (lymphoid bias), and *CEBPA* (GMP bias) (**Fig. 2d,e** and **Supplementary Fig. 9** and **10**). These data further support that the identified genes are relevant to HPSCs.

### Associations with *CXCR4*

Strikingly, four of our most significant signals map to *CXCR4* (C-X-C chemokine receptor type 4) at 2q22 (**Fig. 1**). This receptor binds stromal-derived-factor-1 (SDF-1; also called CXCL12). Among its functions, the CXCR4/SDF-1 axis regulates HSPC and immune cell migration. Particularly, internalization of CXCR4 is required for HSPC egression^30^. In the WHIM syndrome, gain-of-function mutations in the C-terminal region result in an inability to internalize CXCR4 after stimulation, leading to retention of HSPCs and other white blood cells in the bone marrow and low white blood cell counts in blood^31^. CXCR4 inhibitors (Plerixafor/AMD3100) are one of the current methods to mobilize CD34^+^ cells in stem cell donors for leukapheresis^32^. Thus, the fact that we find associations with *CXCR4* provides compelling proof-of-principle for the idea that bona fide regulators of blood CD34^+^ cell levels can be found *in vivo* in humans by GWAS.

The four 2q22 signals represent three common (rs309137, rs11688530, rs10193623) and one rare variant (rs555647251), with a total of 69 variants within their 99% credible sets of probable causal variants (**Supplementary Table 4**). Within each of these sets, we identified a single regulatory variant that either maps to, or has a looping interaction with, the *CXCR4* promoter, making them plausible causal variants (**Fig. 3b**, **Supplementary Fig. 6a**, and **Supplementary Table 4**). In the credible sets of rs11688530 and rs555647251, we identified rs59222832 and rs770321415 as regulatory promoter variants, located 4.7 and 1.3 kb upstream of the transcription start site, respectively. In the other two credible sets, we identified the lead variants rs309137 and rs10193623 as regulatory variants with chromatin looping interactions with the *CXCR4* promoter in CD34^+^ cells^19^.

**Figure 3:**
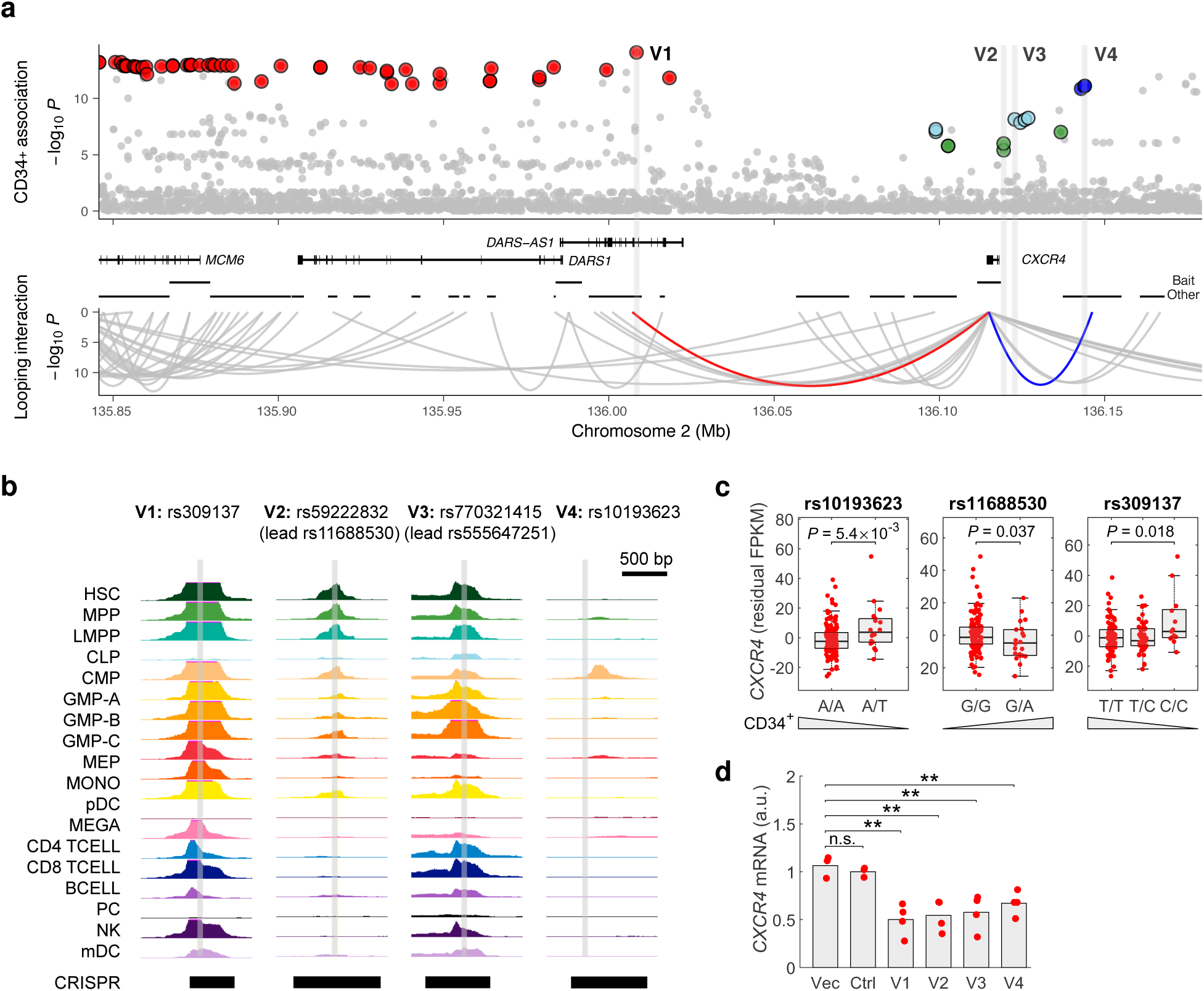
Associations with *CXCR4*. **(a)** We detected four conditionally independent associations at 2p22, represented by a total of 69 credible set variants clustered around *CXCR4* (**Supplementary Table 4**). The credible sets are indicated in red (lead variant rs309137), green (rs11688530), cyan (rs555647251) and blue (rs10193623). The rs309137, rs11688530 and rs10193623 credible sets represent common variants, whereas rs555647251 represents a rare variant. By integrating ATAC-sequencing and PCHi-C data for CD34^+^ cells, we identified a single plausible causal variant within each credible set. rs309137 (“V1”) and rs10193623 (“V4”) have chromatin looping interactions with the *CXCR4* promoter (red and blue arches; *y*-axis indicates PCHi-C *P*-score). rs59222832 (“V2”; credible set of rs11688530), and rs770321415 (“V3”; credible set of rs555647251) map to the *CXCR4* promoter. **(b)** Chromatin accessibility at the four plausible causal variants; *y*-axis indicates ATAC-sequencing signal. **(c)** Using multi-variate regression, we detected conditional *CXCR4 cis*-eQTLs for the three common variants in our CD34^+^ cell mRNA-sequencing data from blood donors. Wedges indicate directions of effects on blood CD34^+^ cell levels. Of note, the effects of these three variants on *CXCR4* expression are anti-directional to their effects on blood CD34^+^ cell levels. Data are residual FPKM values after correction for covariate SNPs and 10 principal components. **(e)** Using dual-sgRNA CRISPR/Cas9, we deleted 587 to 1421- bp regions harboring the four putative causal variants in MOLM-13 cells, resulting in downregulation of *CXCR4*.

We asked if the 2p22 associations are caused by altered *CXCR4* expression in connection to egression, similar to the gain-of-function mutations in WHIM syndrome. In agreement with this hypothesis, analysis of our CD34^+^ cell mRNA-sequencing data for blood donors unveiled conditional *CXCR4 cis*-eQTLs for the three common variants (**Fig. 3c**), while the rare variant was not polymorphic in this data set. Notably, in all three cases, the effect on *CXCR4* expression was anti-directional to the effects on blood CD34^+^ cell levels, consistent with CXCR4 being a negative regulator of egression. For further functional analysis, we used dual-sgRNA CRISPR/Cas9 to delete 587 to 1421-bp regions harboring each of the four putative causal variants in the acute myeloid leukemia cell line MOLM-13 (**Supplementary Table 7; Supplementary Fig. 11** and **12**). In all four cases, we observed downregulation of *CXCR4* (**Fig. 3d**). These data indicate that the 2q22 associations are caused by sequence variation in genomic regions that regulate *CXCR4* expression in HSPCs.

### Association with *PPM1H*

Our most significant association signal maps to *PPM1H* (protein phosphatase, Mg^2+^/Mn^2+^ dependent 1H) at 12q14. This gene encodes an evolutionarily conserved^33^serine/threonine phosphatase in the phosphatase 2C (PP2C) family^34^. Its biological role is unknown, and the few studies that have been done suggest that *PPM1H* could be involved in cellular signaling^35–37^, trastuzumab resistance^38^, lupus^39^, and colon cancer^40^.

The 12q14 association is represented by 32 credible set variants in *PPM1H* intron 1 (**Fig. 4a**, **Supplementary Fig. 6b** and **Supplementary Table 4**). Consistent with the *PPM1H cis*-eQTL in our blood donor CD34^+^ cell mRNA-sequencing data (**Fig. 2b**), the associated region shows chromatin looping interactions in CD34^+^ cells, both with the standard *PPM1H* promoter and internal promoter (**Fig. 4a** and **Supplementary Fig. 6b**). Moreover, by integrating ATAC- and mRNA-sequencing data for 16 sorted blood cell populations^13^, we discovered an approximately 500-bp-long chromosomal segment within the associated region where chromatin accessibility shows strong positive correlation with *PPM1H* expression, suggesting a regulatory role (**Fig. 4b**). This segment harbors four of the 32 credible set variants (rs772555, rs772556, rs772557, rs772559). Congruent with the expression pattern of *PPM1H* across hematopoietic cell types (**Fig. 2c-e** and **Supplementary Fig. 7** to **10**), the segment is selectively accessible in HSCs, MPPs, CMPs, and MEPs (**Fig. 4b**).

**Figure 4:**
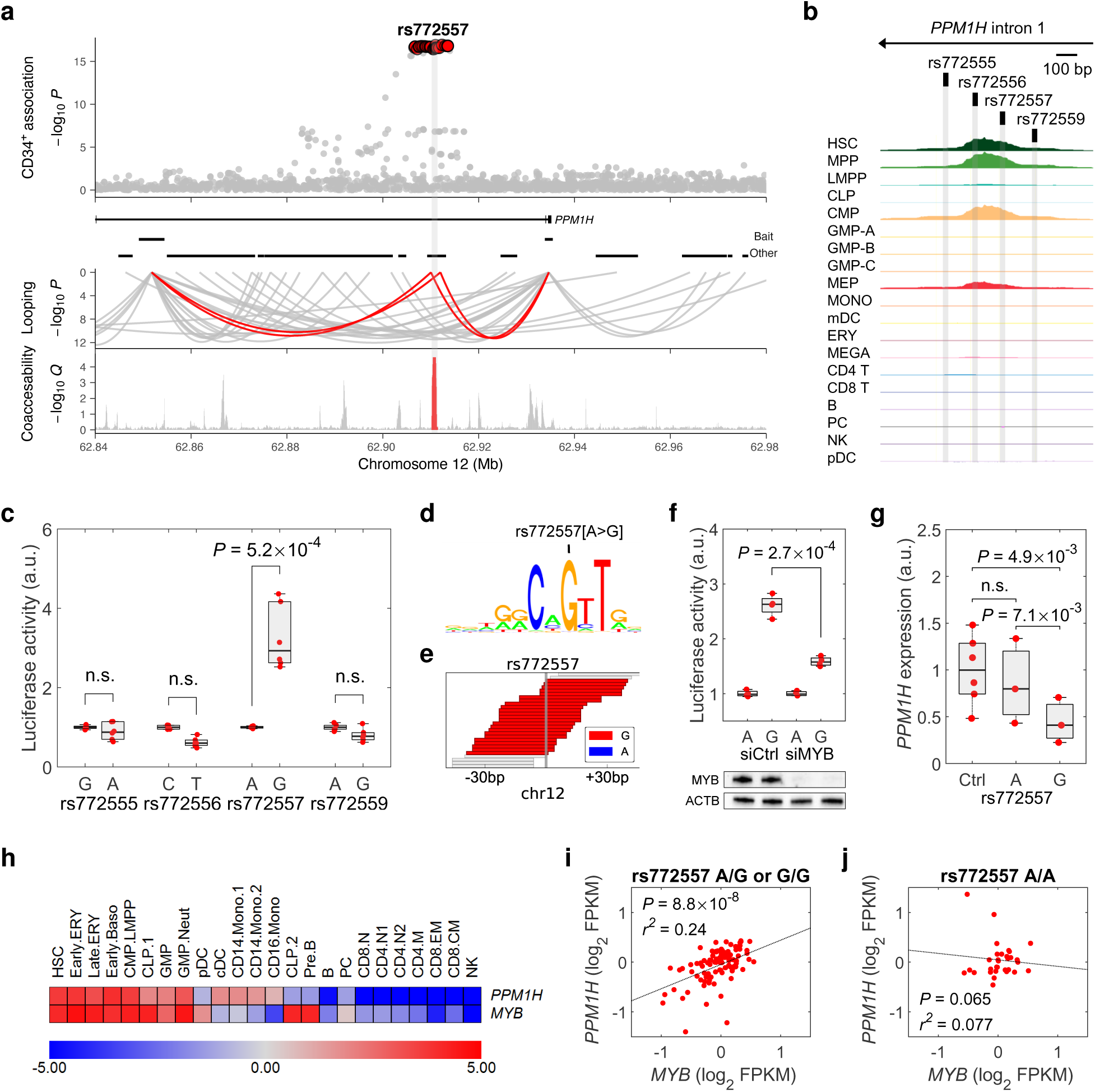
Association with *PPM1H*. **(a)** Top: the 12q14 signal is represented by a credible set of 32 variants in *PPM1H* intron 1. Middle: chromatin looping interactions in CD34^+^ cells with standard and internal promoter (red arches; *y*-axis indicates PCHi-C *P*-score). Bottom: we identified an approximately 500-bp-long chromosomal segment (red peak) where ATAC- sequencing signal (100-bp sliding window) shows strong positive correlation with *PPM1H* expression across 16 sorted blood cell populations^13^ (*y*-axis indicates false discovery rate for Pearson correlation). **(b)** Four credible set variants map to the identified regulatory segment that is accessible in HSC, MPP, CMP and MEPs (*y*-axis indicates ATAC-seq signal). **(c)** Luciferase analysis revealed higher activity with rs772557-G compared to rs772557-A constructs in the *cis*-eQTL direction (*P*-value for one-sided Student’s t-test; Fig. 2b). Signals normalized to hg38 reference (left) alleles. **(d)** MYB binding site that is altered by rs772557; rs772557-G creates binding site, while rs772557-A abrogates it, by changing a critical recognition base (arrow). **(e)** ChIP-seq data for MYB in Jurkat cells (rs772557-heterozygous) show exclusive pull-down of reads harboring rs772557-G. **(f)** siRNA knockdown of MYB in K562 cells selectively attenuates rs772557-G luciferase activity. **(g)** Allele-specific CRISPR- Cas9 disruption at rs772557 (**Supplementary Fig. 14**) in K562 cells, heterozygous for rs772557. We observed *PPM1H* downregulation with rs772557-G, but not with rs772557-A, sgRNA. **(h)** *PPM1H* and *MYB* are co-expressed in hematopoiesis^28^. **(i,j)** Analysis of our CD34^+^ mRNA-sequencing data for blood donors revealed a correlation between *MYB* and *PPM1H* expression in rs772557-G carriers, and no correlation in non-carriers. Data are log_2_- transformed FPKM values, median-centered per genotype group.

We hypothesized that the 12q14 association is caused by one of the four variants in the identified regulatory segment. For functional fine-mapping, we carried out luciferase experiments with constructs representing their reference and alternative alleles in the acute erythroleukemia cell line K562. We observed significantly higher luciferase signal with rs772557-G constructs compared with rs772557-A (**Fig. 4c**), consistent with the direction of the *PPM1H cis*-eQTL (**Fig. 2b**; rs772557[A>G] is in LD with the 12q14 lead variant rs669585[T>G] with *r*^2^ = 0.98). Additionally, we observed higher chromatin accessibility at rs772557-G than at rs772557-A in ENCODE^41^ DNAse-sequencing data for heterozygous primary adult CD34^+^ cells (125 versus 47 reads; Binomial test *P* = 2.0×10^-8^).

Motif analysis predicted that rs772557 alters a binding site for the transcription factor MYB by altering a critical recognition base (**Fig. 4d** and **Supplementary Table 8**). Consistent with the *cis*-eQTL and luciferase data, the rs772557-G allele (which confers high *PPM1H* expression) creates the site, whereas rs772557-A abrogates it. This mechanism-of-action was also supported by chromatin immunoprecipitation sequencing (ChIP-seq) data for MYB in Jurkat cells (heterozygous for rs772557), showing exclusive pull-down of reads harboring the rs772557-G allele (**Fig. 4e** and **Supplementary Fig. 13**). Moreover, luciferase experiments with co-transfected MYB siRNA showed selective attenuation of luciferase signal from rs772557-G but had no impact on rs772557-A, further supporting that MYB only drives transcription at rs772557 in the presence of the high-expressing allele (**Fig. 4f**).

To demonstrate causality, we perturbed rs772557 by CRISPR/Cas9 in K562 cells. These cells are triploid at *PPM1H*, having two copies of the rs772557-G allele and one copy of the rs772557-A allele. Serendipitously, we identified sgRNA sequences that overlap rs772557 and enable allele-specific disruption. These sgRNAs cut DNA only one bp upstream of rs772557, and within the MYB recognition site (**Supplementary Fig. 14** and **Supplementary Table 7**). Consistent with the other data, we observed downregulation of *PPM1H* with rs772557-G sgRNA, but no effect of rs772557-A sgRNA (**Fig. 4g**). Finally, we observed co-expression of *MYB* and *PPM1H* in hematopoietic cell types (**Fig. 4h**), and in our CD34^+^ mRNA-sequencing data for blood donors, we found a highly significant correlation between *MYB* and *PPM1H* among rs772557-G carriers (**Fig. 4i**), but no correlation among rs772557-A homozygotes (**Fig. 4j**). These data identify rs772557 as a causal variant.

Because of the anti-correlation between *PPM1H* expression and blood CD34^+^ cell levels (**Fig. 2b**), we hypothesized that *PPM1H* downregulation by rs772557-A increases the proportion of HPSC subpopulations in which the MYB site is active (*i.e.*, where rs772557 is accessible; **Fig. 4b**). To test this, we first calculated the correlation between rs772557 genotype and gene expression in our CD34^+^ cell mRNA-sequencing data, and tested for enrichment of correlation among genes that are highly expressed in each CD34^+^ subpopulation (**Supplementary Fig. 15**). We observed enrichments of positive correlations in the direction of rs772557-A for CMP, HSC, MEP, and MPP marker genes, and enrichments of negative correlations among marker genes for lymphoid precursors (**Fig. 5a** and **Supplementary Fig. 15**). Second, we quantified 8 HSPC subpopulations in 642 umbilical cord blood samples (**Supplementary Fig. 16**). rs772557-A conferred increased proportion of CMPs, lower proportion of B/NK progenitors (**Fig. 5b**), and unchanged proportions of HSC, MEP and MPP, possibly because these populations are smaller. Third, also consistent with the enrichment analysis, shRNA-knockdown of *PPM1H* in primary CD34^+^ cord blood cell cultures resulted in higher proportions of CD34^+^ and CD34^+^90^+^ cells relative to control shRNA over time (**Fig. 5c** and **Supplementary Fig. 17**). Taken together, our analysis of the 12q14 signal indicates that this association is driven by rs772557, with the rs772557-A allele abrogating a MYB binding site, leading to *PPM1H* downregulation and an increase of HSPC subpopulations in which the MYB site is active.

**Figure 5:**
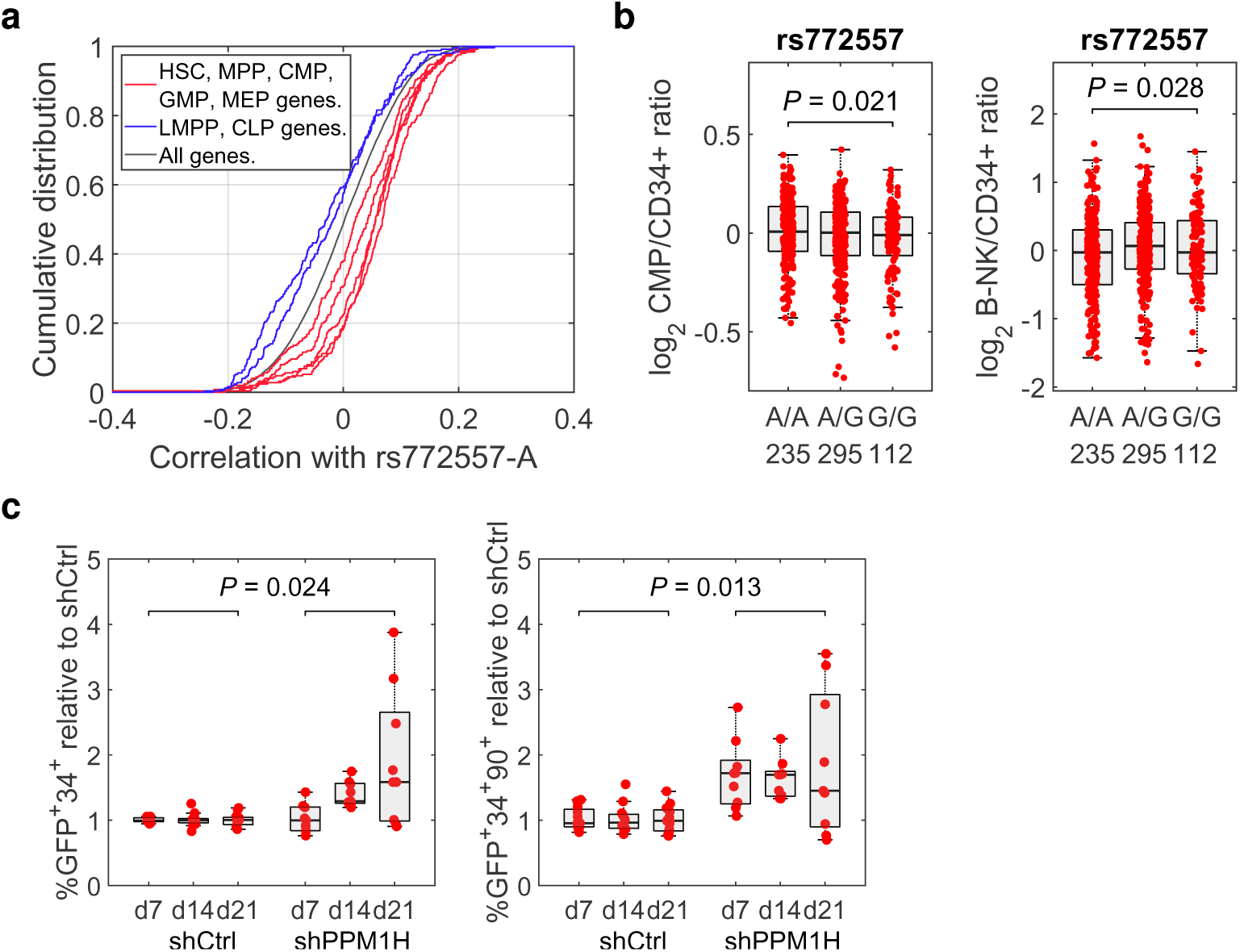
Effects of *PPM1H* downregulation. Since our *cis*-eQTL analysis identified an anti-correlation between blood CD34^+^ cell levels and *PPM1H* expression (Fig. 2), we explored the effects of *PPM1H* downregulation on HSPC levels in better detail. **(a)** We first searched for effects of rs772557 on cell type composition within the CD34^+^ compartment in adults. For this, we calculated correlations between rs772557 genotype and gene expression in our CD34^+^ mRNA-sequencing data for blood donors, and tested for enrichment of correlation within sets of marker genes for HPSC subpopulations (Supplementary Fig. 15). This composite plot shows the distributions of Pearson correlation coefficients for the top 250 marker genes inferred using mRNA-sequencing data for sorted cells^13^. We detected enrichments of positive correlations in the direction of rs772557-A allele for CMP, HSC, MEP, and MPP (red), and of negative correlations for CLP and LMPP (blue), compared to other genes in the genome (black) (details in **Supplementary Fig. 15**). This finding is consistent with rs772557-A increasing the relative abundance of CD34^+^ subpopulations in which the rs772557 region is accessible (Fig. 4b). **(b)** Quantifying HSPC subpopulations in 642 umbilical cord blood samples (**Supplementary Fig. 16**), we observed association between rs772552-A and increased proportion of CMP and lower proportion of B/NK progenitors. **(c)** shRNA-knockdown of *PPM1H* induced an increase in the proportions of CD34^+^ and CD34^+^90^+^ primary cord blood cells out of green fluorescent protein (GFP)- positive. Data are proportion at day 7, 14 and 21 after transduction, normalized to shRNA- control. *P*-value is for permutation testing, taking into account the structure of the experimental design (**Online Methods**).

### Additional associations

Among the remaining significant associations, 1p36/*ENO1-RERE* and 3p22/*ITGA9* displayed strong *cis*-eQTLs and preferential expression of candidate genes in CD34^+^ cells (**Fig. 2b-c**). First, *ENO1* encodes alpha-enolase, a glycolytic enzyme. However, alternative translation also produces a shorter isoform that binds the *MYC* promoter and has been reported to act as a tumor suppressor^42^. *RERE* encodes a transcription factor that forms a complex with the retinoic acid receptor and increases transcription of its target genes^43^. Retinoic acid signalling has been linked to HSC self-renewal in mice^44^. In acute promyelocytic leukemia (APL), genetically perturbed retinoic acid signalling blocks HSPCs from differentiating into mature white blood cells; this block can be overcome by treatment with all-trans-retinoic acid^45^. Concordant with the *cis*-eQTLs, we identified putative causal regulatory variants in the region between *ENO1* and *RERE*, showing chromatin looping interaction with *ENO1* and *RERE* promoters (**Supplementary Fig. 6c** and **Supplementary Table 4**). Second, *ITGA9* encodes the cell surface protein integrin alpha-9. Although one previous study^20^ has linked *ITGA9* expression to HSPCs, its precise function in human hematopoiesis remains unclear. Our analysis now identifies ITGA9 as a functional cell surface marker that participates in the regulation of blood CD34^+^ cell levels. The associated region maps to *ITGA9* introns 3 and 4, and several regulatory variants show looping interactions with the promoter (**Supplementary Fig. 6d** and **Supplementary Table 4**). In addition to the *cis*-eQTL data (**Fig. 2b**), flow cytometry analysis in 458 adults confirmed that the 3p22 variant impacts ITGA9 protein expression on blood CD34^+^ cells (**Supplementary Fig. 18**).

Finally, among the suggestive associations, we particularly noted the 8q24 locus. While this signal primary maps to *CCDC26*, PCHi-C analysis in blood CD34^+^ cells revealed a long-distance looping interaction with the *MYC* promoter, located 1.86 Mb away (**Supplementary Fig. 6h**). The 8q24 signal is represented by 31 variants located between position 129,590,035 and 129,612,415 on chromosome 8 (**Supplementary Table 4**). This region was recently identified as an evolutionarily conserved “super-enhancer” required for *Myc* expression in mouse HSPCs^46^. The super-enhancer comprises eight enhancer modules (A to H; **Supplementary Table 9**), collectively called the Blood ENhancer Cluster (BENC). The 8p24 signal spans BENC module D. Deletion of this module in mice has been found to affect *Myc* expression mainly in HSCs and MPP cells^46^. Our data now identify BENC module D as critical for the regulation of blood CD34^+^ cell levels in humans.

## Discussion

We conducted a large-scale genome-wide association study on blood CD34^+^ levels. Unlike studies in model systems, we exploit natural genetic variation to expose HSPC regulators *in vivo* in humans. The validity of our approach is confirmed by several observations, including enrichment of heritability and gene expression in HSPCs, and discovery of variants at loci that are known to be HSPC-relevant, particularly *CXCR4*.

We identify five novel regulators (*PPM1H*, *ENO1, RERE*, *ITGA9*, and *ARHGAP45*) based on *cis*-eQTLs, chromatin looping, or coding variants. For our strongest signal, we identify an anti-correlation between *PPM1H* expression and CD34^+^ cell levels, indicating that PPM1H could be exploitable clinically as an inhibition target to facilitate stem cell harvesting by leukapheresis. Hence, intriguing challenges ahead will be to map the signalling networks within the PPM1H pathway, and to search for PPM1H inhibitors. Towards this, a first step will be to map PPM1H dephosphorylation targets. While initial proteomic studies in non-hematopoietic cells have identified RAB8A, RAB10, RAB35, SMAD1/2 as tentative targets^35, 37^, the dephosphorylation network in HSPCs remains unexplored. Other notable findings include ENO1 as a regulatory enzyme with links to the MYC pathway, RERE as a transcription factor with links to retinoic acid signalling, and ITGA9 as a functional cell surface marker involved in the regulation of blood CD34^+^ levels.

In all, we report the first large-scale GWAS on a stem cell trait, prove that HSPC regulators can be exposed *in vivo* in humans through genetic variation, and identify several novel regulatory pathways that can be investigated further with the ultimate aim of improving the treatment of blood disorders and stem cell transplantation.

## Acknowledgements

This work was supported by grants from the European Research Council (ERC 770992- BloodVariome), Knut and Alice Wallenberg Foundation (2012.0193, 2014.0071 and 2017.0436), the Swedish Research Council (2017-02023 and 2018-00424), the Swedish Cancer Society (2017/265 and 20.0694), the Swedish Children’s Cancer Fund (PR2018-0118 and TJ2017-0042), and Inga-Britt and Arne Lundberg’s Foundation (2017-0055). We thank Ellinor Johnsson, Christina Jönsson, and Camilla Streimer for their assistance. We are indebted to the personnel at the Clinical Chemistry and Clinical Immunology and Transfusion Medicine units in Region Skåne for their help with sample collection, and to the patients and blood donors who participated in the study.

## Author contributions

A.L.d.L.P, U.T., J.L., K.S., and B.N. conceived the study. A.L.d.L.P, L.E., C.C., Z.A., M.P., K.Ž., U.T., J.L., K.S., and B.N. designed experiments. A.L.d.L.P., C.C., and N.M. developed the flow cytometry platform. L.E. and A.L.A. developed software for flow cytometry data analysis. A.L.d.L.P., L.E., C.C., Z.A., N.M., N.U.D., and D.B. carried out large-scale phenotyping. A.L.d.L.P., C.C., G.T., M.P., U.T. and K.S. carried out genotyping and genomic analyses. C.C., Z.A. N.M., and K.Ž. carried out functional experiments. A.L.d.L.P., L.E., C.C., Z.A., N.M., G.T., K.Ž., A.L.A., M.T., M.P., P.D., E.B., A.N., V.S., G.K., and B.N. carried out bioinformatic and statistical analyses. A.L.d.L.P., L.E., C.C., Z.A. and B.N. drafted the manuscript. All authors contributed to the final manuscript.

## Online methods

### Subjects

For the genome-wide association study, we collected 16,931 peripheral blood samples from random blood donors (n=7,773) and primary care patients (n=9,158) during three time periods (November 2015 to April 2016, “Phase I”, January 2017 to November 2017, “Phase II”, and August 2018 to April 2019 “Phase III”; **Supplementary Table 1**). Samples from blood donors were obtained from Clinical Immunology and Transfusion Medicine, Skåne University Hospital, Lund, Sweden. By Swedish law, men are allowed to donate blood every 3 months, women every 4 months. Hence, by collecting samples for slightly longer periods of time, we ensured collection of repeat donors samples, allowing us to assess reproducibility (**Supplementary Fig. 4**). Samples from primary care patients were surplus material from the clinical routine blood chemistry pipeline at Clinical Chemistry, Skåne University Hospital, Lund, Sweden. By programming the sample processing robot at Clinical Chemistry, samples from patients between 18 and 71 years-of-age from primary care clinics in the Lund area were selected automatically. Making use of the fact that the processing robot had a 100,000- sample history buffer, we minimized the number repeat samples from primary care patients. Samples were collected subject to ethical approval (Lund University Ethical Review Board; dnr 2018/2). Except for information about age and sex (**Supplementary Fig. 3**), the samples were irreversibly anonymized by clinical personnel before being forwarded to us for analysis. Following genotype and phenotype data quality control, and removal of duplicate individuals, a total of 13,167 unique individuals remained (**Supplementary Table 1**). Out of the 9 significant variants, only the rare variant rs555647251 showed Bonferroni-significant heterogeneity between blood donors and primary care patients (**Supplementary Table 10**).

### Flow-cytometry phenotyping for the association study

Collected samples were first diluted 1:10 in 0.84% ammonium chloride and incubated at room temperature for 10 min for erythrocyte lysis. After centrifugation at 1,200 g x 5 min and supernatant removal, leukocytes were resuspended in 20 mL of wash buffer (PBS with 6 mM EDTA) and centrifuged at 1,200 g x 5 min. Washing was done twice. After supernatant removal, cells were resuspended in residual wash buffer, and 60 uL of cell suspension were plated in 96-well plates and centrifuged at 1,200 g x 1 min. Supernatant was removed and pelleted cells were resuspended in 30 uL of staining buffer (PBS containing 6 mM EDTA and 0.1% BSA) including 0.2 µL of APC-H7 mouse anti-human CD45 (clone 2D1; BD #560178) and 0.09 µL of PE-CF594 mouse anti-human CD34, clone 563 (BD #562449), for 15 min at room temperature, in the dark. After a wash step with 100 µL of wash buffer, plates were centrifuged at 1200 g x 1 min, supernatant was removed and pelleted stained cells were re- suspended in 250 µL of staining buffer. Flow-cytometry analysis was performed using a BD FACS Canto II™ (Phase I), BD LSR Fortessa™ (Phase II), and BioRad ZE5™ (Phase III). The numbers of events recorded per sample are summarized in **Supplementary Table 2**.

### Gating of flow-cytometry data for association study

To quantify blood CD34^+^ cell levels, we first gated singlet cells based on forward scatter area (FSC-A) and forward scatter height (FSC-H). From singlets, we then gated peripheral blood mononuclear cells (PBMC) based on FSC-A and side scatter area (SSC-A). Finally, from PBMCs, we gated CD34^+^45^low^ cells and CD45^+^ cells. In CD34 and CD45 intensity space, CD34^+^ cells form a discrete cluster. We defined the CD34^+^ level (frequency) as the number of CD34^+^45^low^ cells divided by the number of CD45^+^ PBMCs (**Supplementary Fig. 1**).

To facilitate the analysis of our flow cytometry data, we developed pattern recognition software (https://github.com/LudvigEk/HSPC-regulators-in-human-blood) that mimics the manual gating strategy. First, to gate singlets, we used an ellipsoid boundary calculated using principal component analysis in FSC-A and FSC-H space (**Supplementary Fig. 1a**). Second, to gate PBMCs, we employed Dijkstraa’s shortest path algorithm to infer an optimal FSC-A-dependent SSC-A boundary between PBMCs and granulocytes by searching for the lowest total event density path from the left edge (minimal FSC-A) to right edge (maximal FSC-A). Having set the Dijkstraa boundary, the definition of PBMCs was refined by overlaying an ellipsoid gate defined by principal component analysis of FSC-A and SSC-A values (**Supplementary Fig. 1b**). Third, in CD34 and CD45 space, debris and CD45^+^ cells were separated by finding the CD45 intensity value corresponding to the lowest event density among CD34^-^ events. Having found a CD45 threshold, the CD34 intensity standard deviation of the CD45^+^ cluster was estimated. Using the estimated parameters, CD34 and CD45 intensity boundaries were inferred for the CD34^+^45^low^ cluster.

To validate our gating software, we compared blood CD34^+^ levels obtained with computer gating to blood CD34^+^ levels obtained with manual gating for 2,838 of our samples, and observed a highly significant correlation (Spearman correlation *P* < 4.94**×**10^-324^ and *r*^2^ = 0.836; **Supplementary Fig. 19**). Phase I and II samples were gated using computer gating. For extra quality control, we plotted the computer gating results (all three gating steps) and visually inspected each of these plots to verify high-quality gating. Phase III samples were gated manually using Flojwo V10.6.1 (Becton, Dickinson & Company).

### Genotyping and association analysis

The Swedish samples were genotyped with Illumina single-nucleotide polymorphism microarrays and phased together with 570,100 samples from North-Western Europe using Eagle2^47^. Samples and variants with <98% yield were excluded. We created a haplotype reference panel by phasing the whole-genome sequence (WGS) genotypes for 15,575 individuals from North-Western Europe, including 3,012 individuals of Swedish ancestry, together with the phased microarray data, and to impute the genotypes from the haplotype reference panel into the phased microarray data using methods described previously^48, 49^.

Genetic ancestry analysis was done in two stages for the Swedish sample sets. Firstly, ADMIXTURE v1.23^50^ was run in supervised mode with 1,000 Genomes populations CEU, CHB, and YRI^51^ as training samples and Swedish individuals as test samples. Input data for ADMIXTURE had long-range LD regions removed^52^ and was then LD-pruned with PLINK v.190b3a^53^ using the --indep-pairwise 200 25 0.3 option. Samples with < 0.9 CEU ancestry were excluded. Secondly, remaining samples were projected onto a principal component analysis (PCA), calculated with an in-house European reference panel to calculate the 20 first principal components for each population. UMAP^54^ was used to reduce the coordinates of test samples on 20 principal components to two dimensions. Additional European samples not in the original reference set were also projected onto the PCA and UMAP components to identify ancestries, and samples with Swedish ancestry were identified. This included 10,949 unique individuals with blood CD34^+^ cell level measurements. After excluding samples of Swedish ancestry, the process was repeated for the remaining samples with blood CD34^+^ cell level measurements and 2,218 unique individuals with > 0.7 CEU ancestry were identified and principle components were created separately for those individuals. We used linear regression to test for association between blood CD34^+^ cell levels and genotypes for the discovery and follow-up sets separately using deCODE software^48^. Prior to association the blood CD34^+^ level measurements were adjusted for gender and 20 principal components and standardized using an inverse normal transform. The association was restricted to 18,061,173 variants with MAF > 0.05%, imputation info > 0.9 and that passed various quality test. We used LD score regression to account for distribution inflation due to cryptic relatedness and population stratification^55^.

For the meta-analysis we used a fixed-effects inverse variance method to combine the results from the discovery and follow-up dataset^56^. We tested for heterogeneity by comparing the null hypothesis of the effect being the same in both sets to the alternative hypothesis of each set having a different effect using a likelihood ratio test (Cochrańs Q) reported as *P*_het_. Genome-wide significance was determined using class-based Bonferroni significance thresholds adjusting for the 18 million variants tested. Sequence variants were split into classes based on their genome annotation, and the significance threshold for each class was based on the number of variants in that class^57^. The adjusted significance thresholds used are 4.5×10^−7^ for variants with high impact (including stop-gained and stop-loss, frameshift, splice acceptor or donor and initiator codon variants), 9.0×10^−8^ for variants with moderate impacts (including missense, splice-region variants, in-frame deletions and insertions), 8.2×10^−9^ for low-impact variants (including synonymous, 3′ and 5′ UTR, and upstream and downstream variants) and 1.4×10^−9^ for all other variants, including those in intergenic regions.

We used step-wise conditional analysis using genotype data to look for additional signals at each locus. The conditional analysis was done for the discovery and follow-up dataset separately, and the results combined. At each locus, we tested all variants in a 2 Mb region centred at the lead variant and significance threshold was set at 1×10^-6^ for a secondary signal. We calculated 99% credible sets of plausible causal variants for each independent association signal^58^, weighting the variants using the same weights as were used to define the class specific genome-wide significance thresholds.

### Chromatin accessibility data

Previously published ATAC-seq data for sorted hematopoietic cell types were downloaded from the Sequence Read Archive^17, 18^. Reads were processed as the MM ATAC-seq libraries using the hg38 reference genome. Next, we created a master peak file by aggregating the summits of each population and enumerating the fragments overlapping each peak for each population as previously described. In addition to the ATAC-seq data for sorted blood cells, we used DNA DNAse-sequencing data for heterozygous primary adult CD34^+^ cellsfrom ENCODE Phase 3 (accession no. ENCSR098PTC, ENCFF971ZOL, ENCFF814EOK).

For MOLM-13 cells, we generated ATAC-seq libraries from 50,000 cells as described in ref^59^. Libraries were prepared using the Nextera DNA Library Prep kit and sequenced on an Illumina Novaseq 6000 Sequencing System with a read length of 50 bases in paired-end mode. Adapter sequences in sequence reads were removed using Trimmomatic (v0.36^60^) and aligned using Bowtie2 to hg38. Duplicate and mitochondrial reads were filtered out using SAMtools^61^ and Picard (http://broadinstitute.github.io/picard).

### ChIP-seq data

To test of allele-specific binding of MYB to rs772557, we used MYB ChIP-sequencing data for Jurkat cells^62^. Raw SRA files were obtained from the Sequence Read Archive (accession no. SRR1603653) and converted to FASTQ format using fastq-dump. The raw FASTQ file was aligned using bowtie2, and the resulting SAM file was sorted and indexed using samtools. For Jurkat WGS, a SAM file containing the region of interest was downloaded from the Sequence Read Archive (accession no. SRR5349449^63^).

### Heritability estimation

Total SNP heritability was estimated with the LDSC software^55, 64^ with default settings, intercept 1.0, and Baseline model v1.1 as control. To estimate cell type-specific partitioned heritability based on chromatin accessibility, we used LD-scores based on ATAC-sequencing data for sorted blood cell types available for LDSC^55^, extended with ATAC-sequencing data for mDC and pDC^18^ (NCBI Gene Expression Omnibus accession no. GSE119453).

### Promoter capture Hi-C data analysis

A table of significant PCHi-C contacts for CD34+ cells was obtained from the ArrayExpress repository (accession number: E-MTAB-2323^65^). These data had been previously processed using the GOTHiC Bioconductor package and filtered to interactions with FDR < 0.05 in two biological replicates. The log of the observed/expected ratio shown in locus plots is an estimate of effect size or interaction frequency.

### Expression quantitative trait locus (eQTL) analysis in blood CD34^+^ cells

We purified CD34^+^ cells from 155 leukocytes depletion filters (Reveos^TM^) from random blood donors within 8 h of donation. We first enriched CD34^+^ cells using the MACSprep CD34^+^ MicroBead kit (Miltenyi Biotech; #130-120-673). The enriched cells were stained using 5 uL of APC-H7 mouse anti-human CD45, clone 2D1 (BD #560178) and 22 uL of PE- CF594 mouse anti-human CD34, clone 563 (BD #562449) for 15 min at room temperature. Cells were washed with 500 µL of wash buffer and centrifuged at 900 rpm g x 5 min. Pelleted stained cells were re-suspended in 350 µL of staining buffer. CD34^+^CD45^dim^ mononuclear cells were sorted using a BD FACSAria^TM^ III (BD Biosciences) into lysis buffer (Qiagen; RLT buffer #74004; containing 1% β-mercaptoethanol), vortexed, and snap frozen. From each sample, 14,000 to 358,000 cells (average 122,000) were sorted. The samples were thawed on ice and RNA was extracted using the RNeasy plus micro kit (Qiagen; 74034) and treated with RNAse-Free DNase Qiagen; #79256).

cDNA libraries were prepared using the SMARTer Stranded Total RNA-Seq Kit v2 Pico Input Kit (Clontech, Takara) and sequenced on an Illumina NovaSeq 6000 Sequencing System with a read length of 100 bases in paired-end mode. After trimming the first three bases using Trimmomatic v0.36^60^, the sequenced reads were aligned to the human reference genome (human GRCh38 primary assembly) using STAR (v2.5.2b^66^). Fragments aligned to human genes were quantified using featureCounts^67^, with the following settings: paired-end mode (-p), strand specific (-s 2), no multimapping reads counted, counting exonic reads. Raw gene counts were transformed to FPKM using in-house Python scripts.

To test for associations between variants and gene expression, we used linear modeling with the variant genotype is independent variable and 10 principal-component covariates, calculated using genes with average FPKM > 1.0 in our mRNA-sequencing data set. For *CXCR4*, we included the three common variants as independent variables in the same model. *P*-values were calculated using Student’s *t*-test for the independent variables.

### Gene expression data and analysis

To map candidate gene expression in hematopoietic cell types, we used published single-cell mRNA sequencing data for 35,582 mononuclear cells from blood and bone marrow^28^, and bulk mRNA-sequencing data for sorted blood cell populations^18^. The definitions of the cell populations in these data sets are detailed in the original publications. Additionally, we used CITE-seq data on 4,905 CD34^+^ cells from adult bone marrow. Definitions of cell populations are detailed in ref.^29^. For dimension-reduction of single-cell data, we used uniform manifold approximation and projection (UMAP)^68^, and imputed gene expression using the MAGIC tool^69^. Plots were generated using SCANPY^70^.

### CRISPR/Cas9 perturbation of variant regions

We used dual-sgRNA CRISPR/Cas9 to delete the regions harboring the four candidate causal variants at *CXCR4* (**Supplementary Table 7**), and an allele-specific single-sgRNA CRISPR/Cas9 approach to abrogate the MYB binding site at rs772557 at *PPM1H* (**Supplementary Fig. 14**). sgRNAs were cloned into the pSpCas9(BB)-2A-GFP PX458 vector (Addgene; #48138), and transfected into MOLM13 cells (*CXCR4* variants) or K562 cells (*PPM1H* rs772557 variant) using the Neon system (Thermo Fisher Scientific). At 24 hours post-transfection, GFP-positive cells were isolated by fluorescence-activated cell sorting. To verify CRISPR/Cas9 deletion in MOLM13 cells, DNA was extracted, the targeted regions amplified by PCR (**Supplementary Table 7**), purified using the NucleoSpin Gel and PCR Clean-up kit, loaded onto a 1% agarose gel. The deletion efficiency was estimated as the intensity of the deletion band divided by the sum of the intensity of the deletion band and the intensity of the wild type band. For allele-specific CRISPR/Cas9 towards rs772557 in K562 cells, perturbation was verified using Sanger sequencing and the Tracking of Indels by Decomposition (TIDE, https://tide.nki.nl/). As control, we used sgRNAs targeting a random non-coding region on a different chromosome (here an intronic region in the *WAC* gene on chromosome 10). RNA was extracted from the same cells using the RNeasy plus micro kit (Qiagen; #74034), reverse-transcribed using Sensiscript RT Kit (Qiagen; #205213) and Oligo-dT Primers (Qiagen; #79237), and quantified by Taqman assays (Thermo Fisher; #Hs00324748_m1 for *PPM1H* and Hs01060665_g1 for *ACTB*, as control).

### Luciferase experiments

For the *PPM1H* variants, 200-bp sequences representing the reference and alternative allele of rs772555, rs772556, rs772557 and rs772559 in their genomic contexts were synthesized as gBlocks and cloned into the pGL3-Basic plasmid using KpnI and BglII restriction sites. Each sequence was centered on the variant and the two constructs differed only for the variant. 240 ng of Renilla luciferase construct were co-transfected with 10 ug of firefly construct in 5 million K562 ells, using the Neon electroporation system (Thermo Fisher Scientific). At 24 h post-electroporation, luciferase and renilla activity were measured using DualGlo Luciferase (Promega; #E1960) on a GLOMAX 20/20 Luminometer. Based on luciferase/renilla readings, log_2_ scores were calculated for each variant reflecting the luciferase activity of the alternative allele relative to the corresponding reference allele. For co-electroporation experiments with *MYB* siRNA, luciferase plasmids were co-transfected with 400 nM Qiagen FlexiTube siRNA solution targeting *MYB* (Qiagen; #1027415) or 400nM Qiagen Negative Control siRNA (Qiagen; #1022076). Lysates for luciferase activity measurement and Western blot were collected simultaneously, 24 hours after electroporation. Electroporation conditions were not modified for co-transfection with siRNA and knockdown was confirmed by Western Blot using a recombinant c-Myb antibody (Abcam; #ab109127).

### Motif analysis

To identify differentially binding transcription factors, we used the PERFECTOS-APE tool (http://opera.autosome.ru/perfectosape) with the HOCOMOCO-10, JASPAR, HT-SELEX, SwissRegulon and HOMER motif databases.

### Enrichment analysis

To identify effects of rs772557 on specific CD34^+^ populations, we calculated the correlation between rs772557 genotype and the expression of all genes with median FPKM > 1.0 in our CD34^+^ mRNA-sequencing data set for blood donors. We then tested for enrichment for correlations within sets of marker genes for different CD34^+^ HSPC subpopulations, inferred by comparing the expression profile of each cell type to other CD34^+^ cell types in three data sets: **(a)** bulk RNA-sequencing data for sorted blood cells^18^; **(b)** single-cell RNA-sequencing data for CD34^+^ cells^29^; and **(c)** single-cell RNA sequencing data for mononuclear white blood cells from adult blood and bone marrow^28^. For analysis we used RenderCat^71^ with default settings. For completeness, we carried out the analysis with varying numbers of marker genes (25 to 500), with agreeing results.

### Quantification of CD34^+^ cell populations in cord blood

Umbilical cord blood (CB) samples were obtained from newborns at Skåne University Hospital (Lund and Malmö, Sweden) and Helsingborg Hospital (Helsingborg, Sweden), in compliance with regulations set by the regional ethical committee and informed consent. Samples were collected in Dulbecco’s modified Eagle medium (DMEM) (GE Healthcare Life Sciences; #SH30022.01) with 1% fetal bovine serum (Fisher Scientific; #SV30160), 1% penicillin-streptomycin (Cytiva; #SV30010) and 4% heparin (Leo Pharmaceutics; 20U/ml). Samples were processed within 48 hours by isolating mononuclear cells using the density-gradient method (Abbott; #1019817). Mononuclear cells were isolated and frozen in FBS (Fisher Scientific; #SV30160) with 10% dimethyl sulfoxide (Sigma-Aldrich; #D5879), and kept at -80°C until analysis. For flow cytometry analysis, cells were thawed, incubated with 0.8% ammonium chloride (Stem Cell Technologies; #07850) for 5 minutes at room temperature to remove remaining red blood cells, washed, filtered (BD #340632), and transferred to a polypropylene 96-well plate. After pelleting by centrifuging at 300 g for 5 min, cells were resuspended in 30ul antibody mix (**Supplementary Table 11**), diluted in staining buffer (PBS with 4mM EDTA and 2% BSA), and incubated for 60 min at 4°C in the dark with shaking. Cells were then washed, resuspended in 200 µL PBS with 4mM EDTA and 2% BSA and analyzed using a four-laser BioRad ZE5^TM^ .

### Gating of flow-cytometry data for cord blood

Cord blood samples were gated using pattern recognition software developed in-house (https://github.com/LudvigEk/HSPC-regulators-in-human-blood) that implements the strategy described in **Supplementary Fig. 16**. Similar to the strategy used in adult blood, singlets were separated using principal component analysis, and the PBMC gate also used the Dijkstraa’s shortest path algorithm to separate granulocytes from PBMCs. Debris with low scatter values were also removed at this gate. The CD34^+^ cluster was identified by finding the highest intensity value among the CD45^+^34^+^ cells, which often corresponds to the center of the cluster. The multiple gates involving lineage markers used the lowest density point in the corresponding marker-axis to separate the cells into two groups each time. The CD38 gate was fixed, considering the 30% of cells with the lowest CD38 values to be CD38- cells, while the rest were determined to be CD38^+^. B/NK progenitors, CMPs, GMPs, MEPs, HSCs, MLPs and MPPs were determined in each case by different thresholds on CD10, CD135 and CD45RA values. These thresholds were determined by first identifying the highest density point in the distribution of the relevant marker values, and then locating a point in a predefined interval where the density dropped below a specific fraction of that value.

### shRNA-knockdown in umbilical cord blood cells

Umbilical cord blood samples were collected as in the previous section. Thawed CB-derived CD34^+^ cells were sorted and cultured in tissue-treated plates. Cells were maintained in serum-free expansion medium (SFEM) (Stem Cell Technologies; #09650) with penicillin-streptomycin (Cytiva; #SV30010), supplemented with stem cell factor, thrombopoietin, and FLT3-ligand at 100 ng/ml (Peprotech; #130-096-696, #130-095-754 and #130-096-480). pLKOpuro clones containing these shRNA sequences targeting *PPM1H* were obtained: CCCTCATTGTGATTTGCCTTT (TRCN0000331708), GCTTGGAAAGAAGATGCTCTA (TRCN0000052768), CCCAAACAGGAACTCATCCAA (TRCN0000052771) (Sigma), and recloned into pLKO.1-EGFP vector. Lentiviruses were produced in 293T cells as previously described^72^. Knockdown efficiency was assessed in K562 cells cultured in RPMI media with 10% FBS. GFP^+^ cells were sorted 2 days after transduction using a FACSAria^TM^ III (BD Biosciences). RNA was extracted with the RNeasy Plus Micro Kit (Qiagen; #74034). cDNA was generated using the Sensiscript RT Kit (Qiagen; #205211) with poly-dT primers.

*PPM1H* expression was quantified with TaqMan probes from ThermoFisher Scientific with ACTB as an internal control (*PPM1H* assay ID: Hs00324748_m1; *ACTB* assay ID: Hs01060665_g1). In the subsequent experiments on primary umbilical cord blood cells, transduced cells were analysed using a BD LSR Fortessa flow cytometer after staining with 1:200 CD34-PeCy7 (BioLegend, #343516), 1:100 CD90-APC (BioLegend, #328114), and resuspended with CountBright™ Absolute Counting Beads (Fisher Scientific; #C36950) and 7-AAD for dead cell exclusion. As readout, we measured the percentage and absolute count (using the CountBright beads) of CD34^+^ and CD34^+^90^+^ cells, the latter known to be highly enriched for hematopoietic stem cells. Cell enrichment was calculated by dividing CD34^+^ and CD34^+^90^+^ cells counts at each time point by initial cell counts, and then normalizing the enrichment values to the control shRNA transduced cells at the same time point. At each time point, three transduction replicates were recorded. The experiment was repeat three times. For statistical analysis, we compared normalized enrichment values at day 7, 14 and 21 for shRNA-transduced cells vs non-targeting shRNA control. To ensure conservative statistical analysis, we first averaged the transduction replicates for each experiment, then calculated *P*- values using permutation testing with the data the day 7, 14, and 21 data for the same experiment being permuted together (100,000 permutations).

### Protein quantitative trait locus (pQTL) analysis

Using flow-cytometry, we quantified ITGA9 protein expression on CD34^+^ cells in blood samples from 458 primary care patients, not included in the association study. Samples were obtained and processed as described above. Cells were stained with CD34-PE-CF594, CD45- APC-H7 (**Supplementary Table 11**), and ITGA9-PE (Biolegend # 351606) at concentration 0.02µl of antibody per 30µl of total staining volume. Gating and calculcation of median fluorescence intensity was done with FlowJo V10.6.1 (Becton, Dickinson & Company).

