## Supplementary Tables and Figures for "Genome-wide association study on 13,167 individuals identifies regulators of hematopoietic stem and progenitor cell levels in human blood"

### Supplementary Figure 1

Quantification of $CD34^+$ cells in peripheral blood. In each sample, we analyzed up to 1 million cells using high-throughput flow cytometry. **(a)** First, we gated single cells based on forward scatter area and forward scatter height. **(b)** Second, we gated mononuclear cells based on side scatter area and forward scatter area. **(c)** Third, we gated $CD34^+$ cells (red) and $CD45^+$ cells (top left population). $CD34^+$ cells form a discrete, well-defined cluster in the CD34-CD45 intensity space. We defined the $CD34^+$ level as the number of $CD34^+$ cells divided by the number of $CD45^+$ cells.

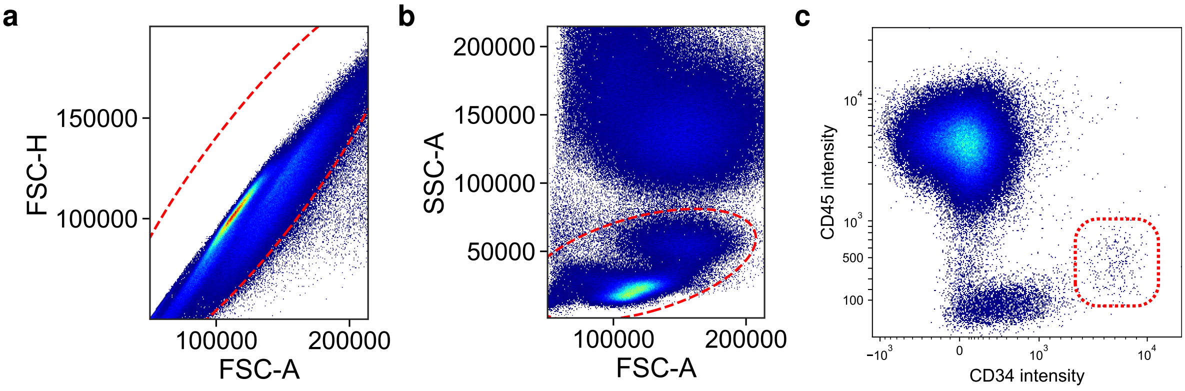

#### Supplementary Figure 2

Representative example of flow cytometry data illustrating the distribution of the different HSPC subpopulations within the CD34$^+$ compartment in peripheral blood. A total of 2 million flow-cytometry events were analyzed. Peripheral blood mononuclear cells (PBMC) were gated on forward (FSC-A) and side scatter area (SSC-A). After excluding doublets based on FSC-A and forward scatter height (FSC-H), HSPC were defined as CD34$^+$45$^{low}$, and divided into CD38$^-$ and CD38$^+$ cells. Among CD38$^+$ cells, B-NK progenitors were defined as CD45RA$^+$10$^+$ cells, while CD10$^-$ cells were subdivided into common myeloid progenitors (CMP), granulocyte-monocyte progenitors (GMP) or megakaryocyte-erythrocyte progenitors (MEP) by CD135 and CD45RA expression. CD38$^-$ cells were classified as hematopoietic stem cells (HSC), multipotent progenitors (MPP), and multilymphoid progenitors (MLP) by CD90 and CD45RA expression.

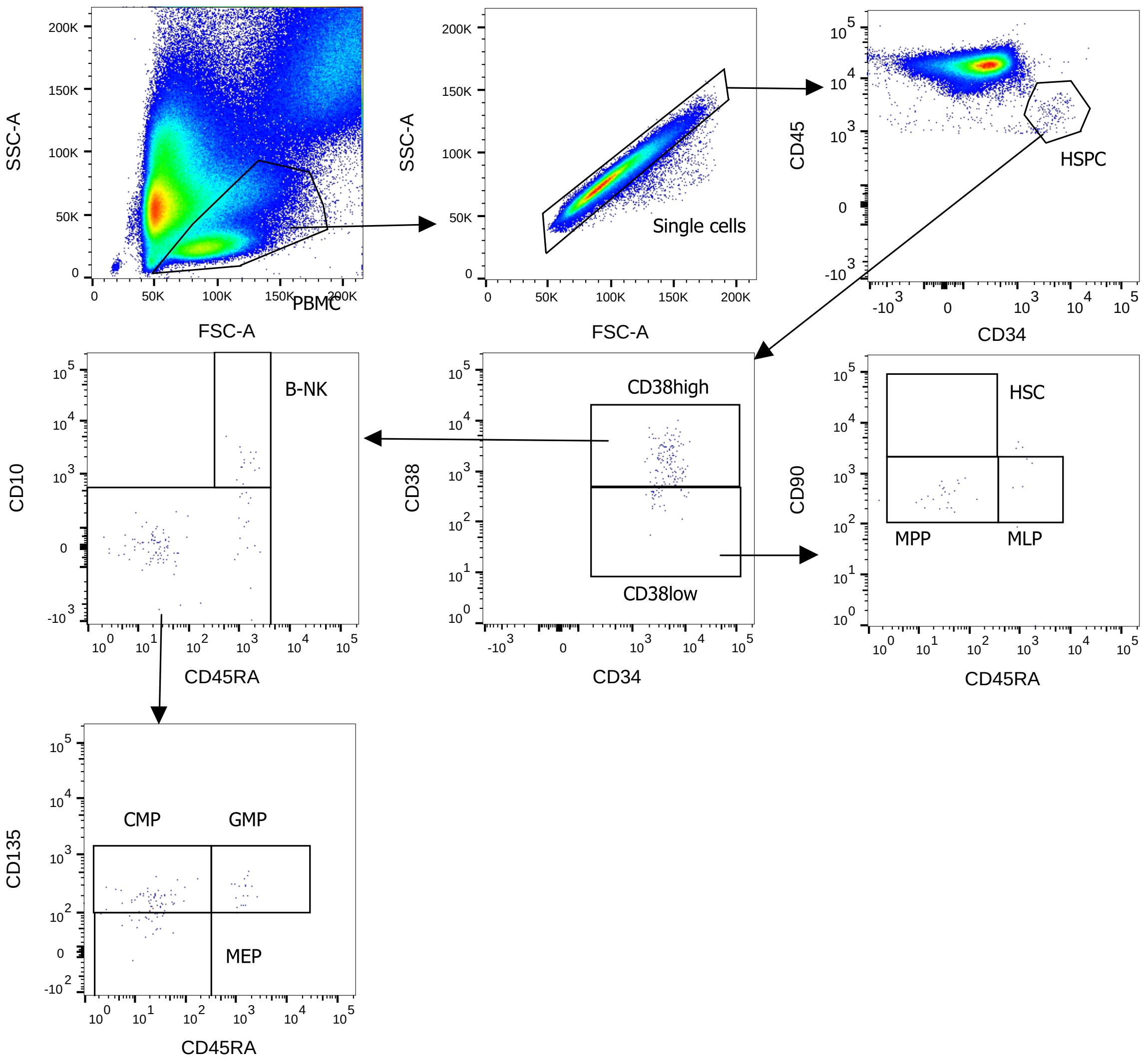

#### Supplementary Figure 3

Distribution of ages. Age information was available for 8,294 unique participants.

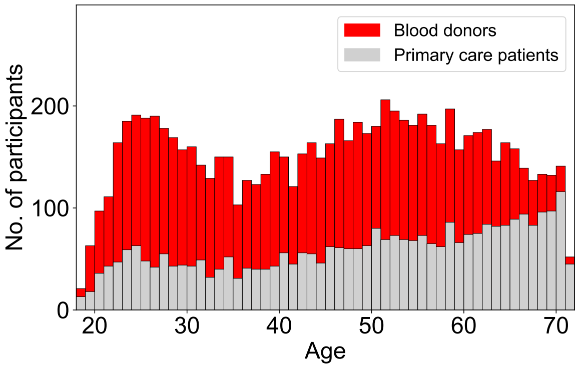

#### Supplementary Figure 4

To assess reproducibility, we analyzed CD34$^+$ levels twice in 660 individuals, with 3 to 36 months between samplings. This figure illustrates the correlation in CD34$^+$ level between replicates. The *r* and *P*-values are for Spearman correlation.

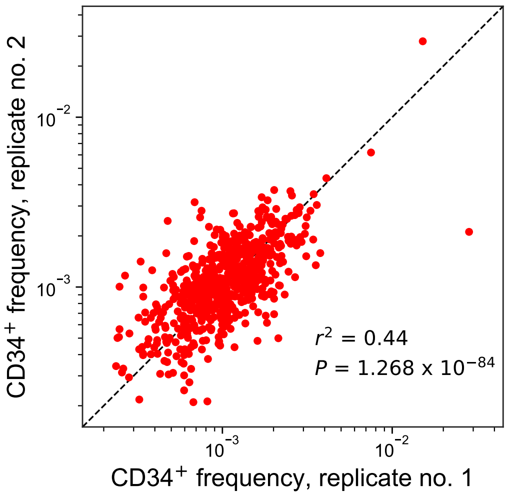

### Supplementary Figure 5

Differences in $CD34^{+}$ levels between men and women, and between age groups.

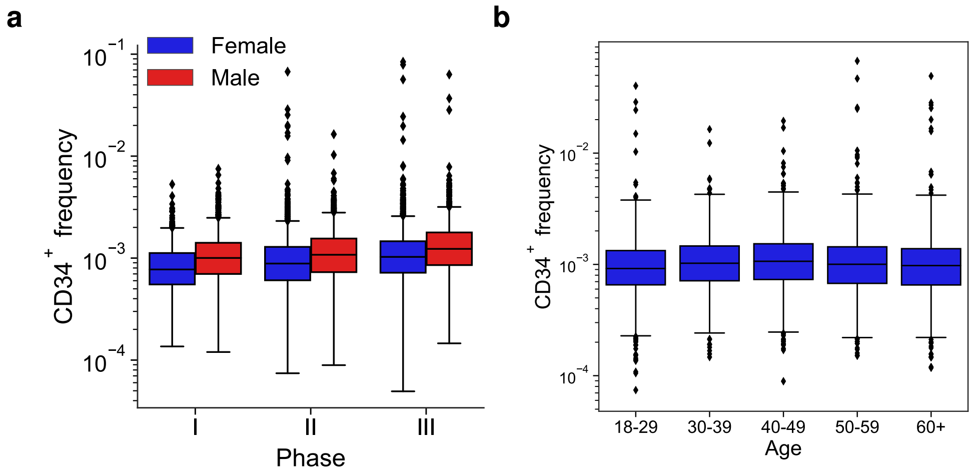

### Supplementary Figure 6

Locus plots for 6 significant (**a** to **f**) and 2 suggestive loci (**g** to **h**; also indicated by *). **Top panels:** Association with $CD34^+$ levels. For 2p22/*CXCR4*, the plot shows the four independent signals, in the order they were identified in the step-wise conditional association analysis. **Mid panels:** Chromatin looping interactions. Red arches indicate interactions between regulatory variants that map to a gene promoter or a genomic region with a looping interaction with a promoter, as determined by by PCHi-C in primary adult $CD34^+$ cells (data from Mifsud *et al.*, *Nature Genetics* 2015;47(6):598-606). As regulatory variants, we considered variants mapping to open chromatin in HSPCs, as determined by ATAC-sequencing. **Bottom panels:** Correlation between ATAC-sequencing signal (100-bp sliding window) and mRNA-sequencing signal (both data sets from Ulirsch *et al.*, *Cell* 2016;165(6):1530-1545).

**(a)**

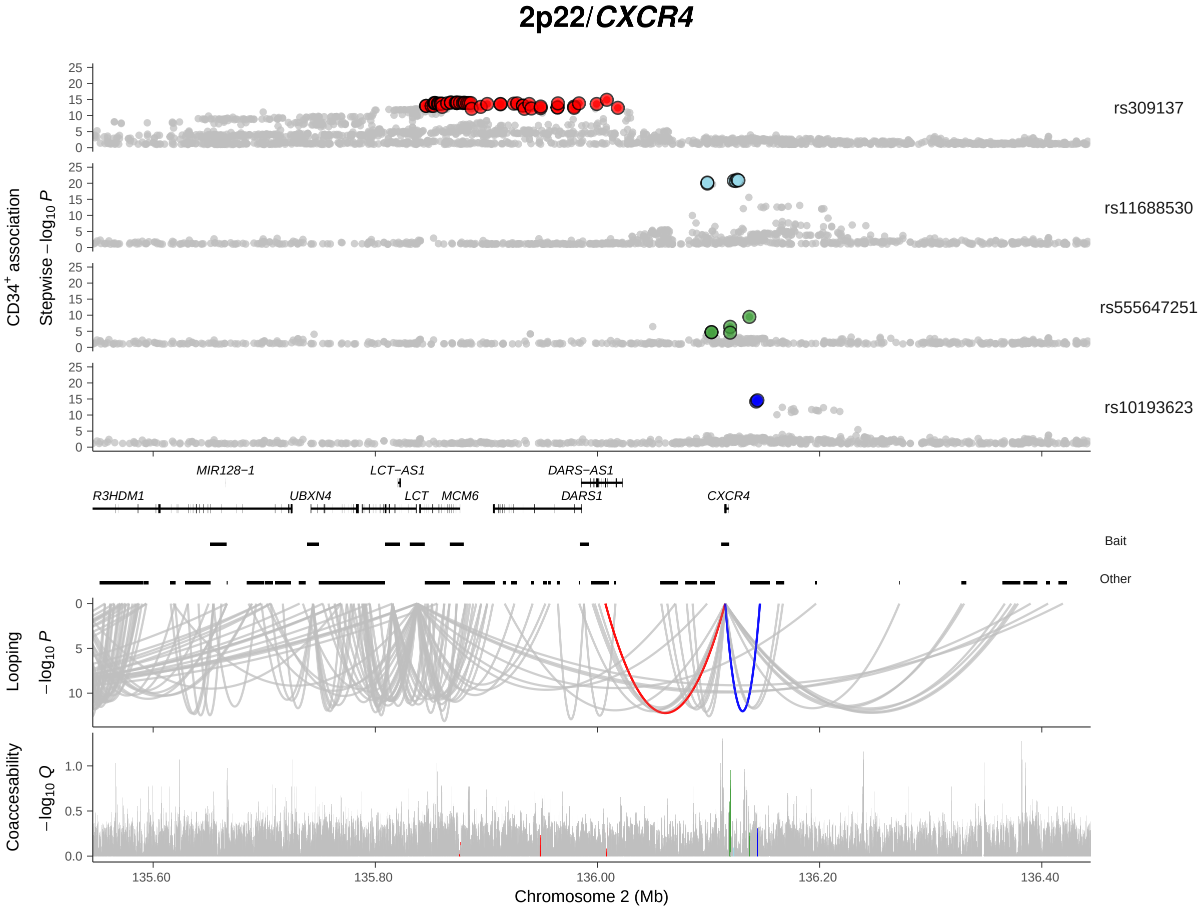

(b)

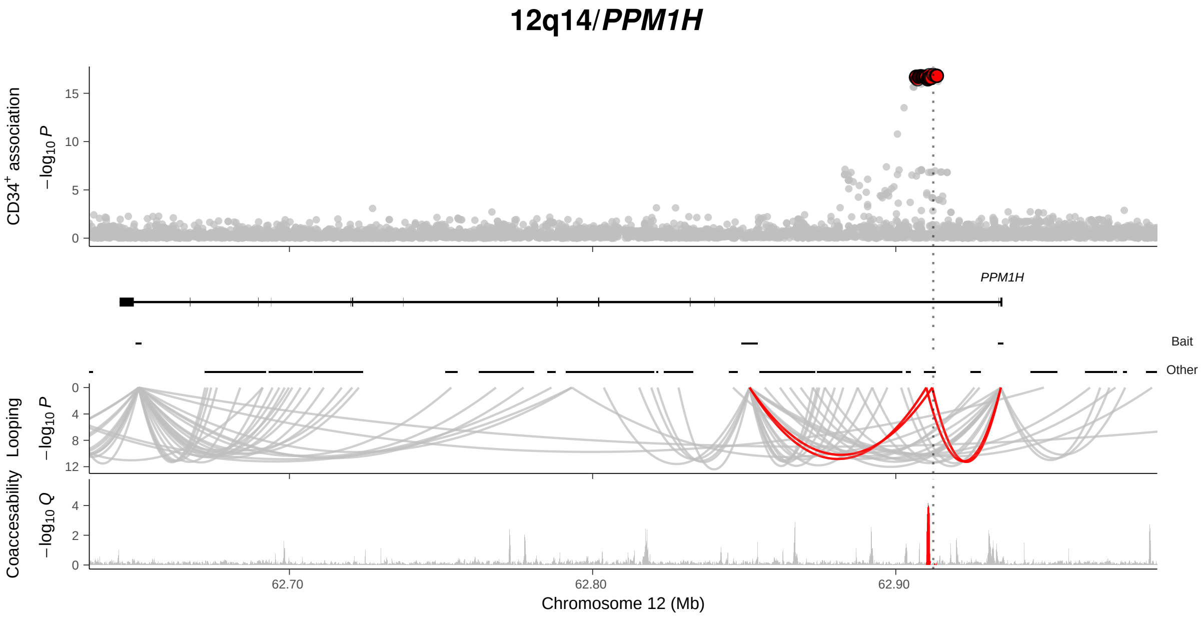

12q14/PPM1H
CD34+ association
−log10 P
PPM1H
Bait
Other
Looping
−log10 P
Coaccesability
−log10 Q
62.70
62.80
62.90
Chromosome 12 (Mb)


(c)

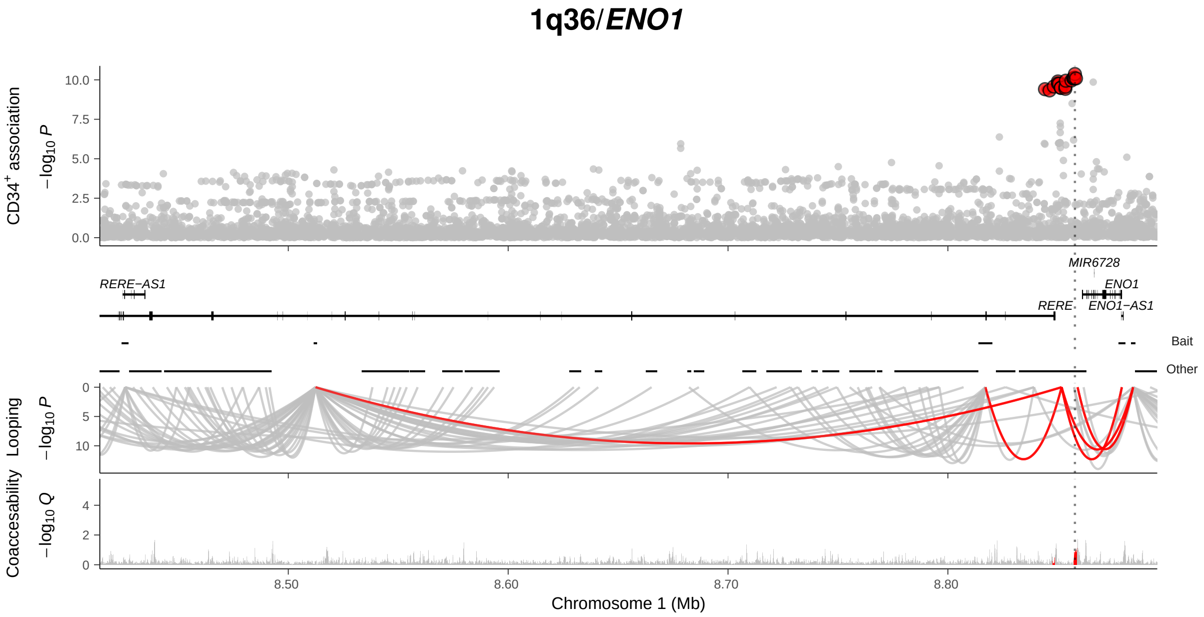

1q36/ENO1
CD34+ association
−log10 P
MIR6728
RERE−AS1
ENO1
RERE
ENO1−AS1
Bait
Other
Looping
−log10 P
Coaccesability
−log10 Q
8.50
8.60
8.70
8.80
Chromosome 1 (Mb)

**(d)**

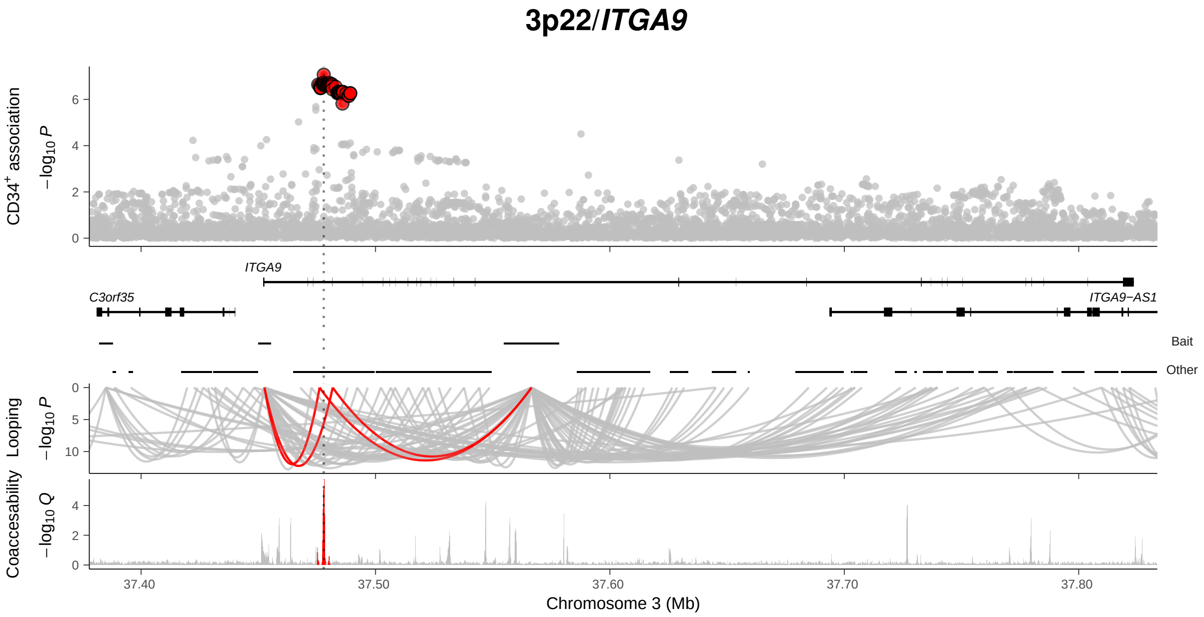
3p22/ITGA9
CD34+ association
−log10 P
ITGA9
C3orf35
ITGA9−AS1
Bait
Other
Looping
−log10 P
Coaccesability
−log10 Q
37.40
37.50
37.60
37.70
37.80
Chromosome 3 (Mb)

**(e)**

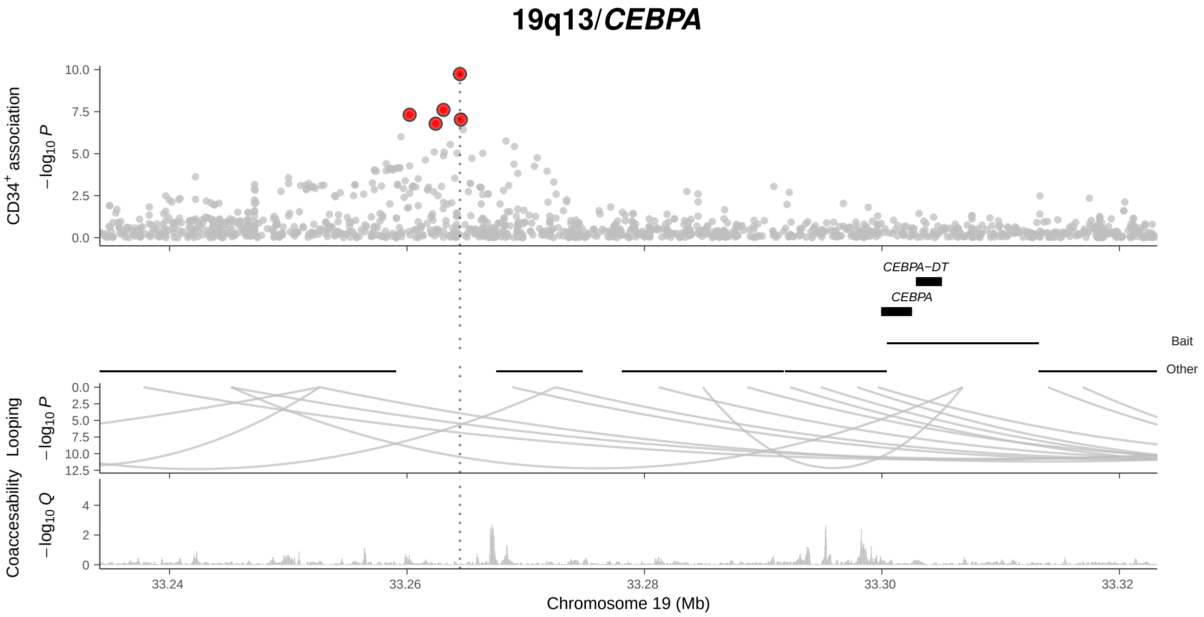
19q13/CEBPA
CD34+ association
−log10 P
CEBPA−DT
CEBPA
Bait
Other
Looping
−log10 P
Coaccesability
−log10 Q
33.24
33.26
33.28
33.30
33.32
Chromosome 19 (Mb)

(f)

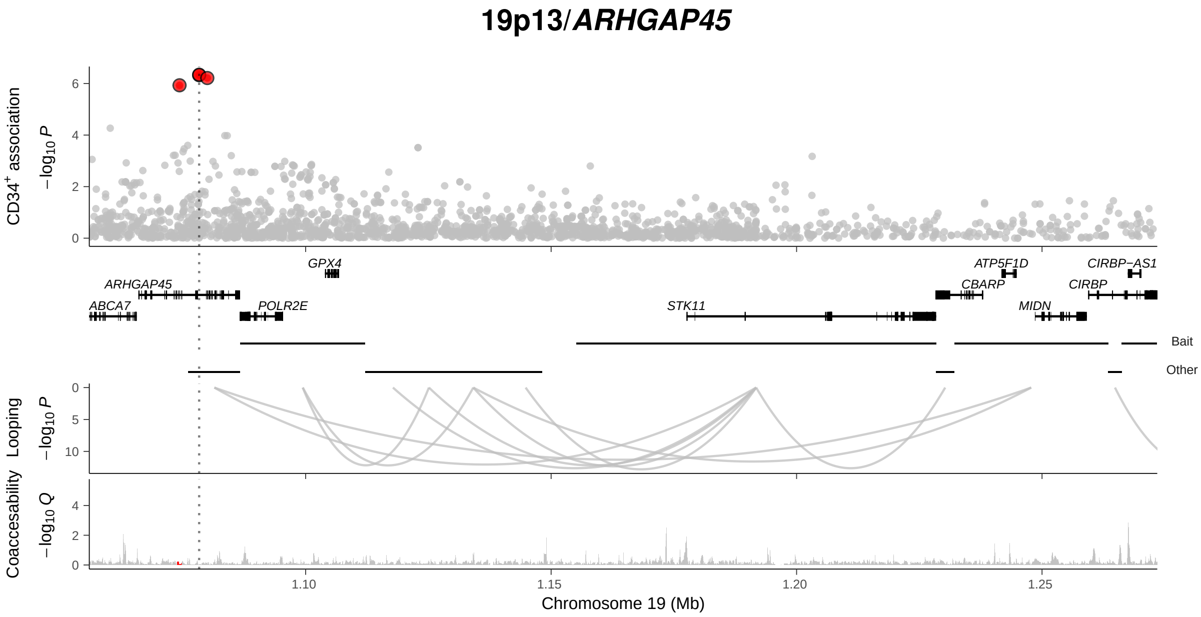
19p13/ARHGAP45
CD34+ association
−log10 P
GPX4
ATP5F1D
CIRBP−AS1
ARHGAP45
CBARP
CIRBP
ABCA7
POLR2E
STK11
MIDN
Bait
Other
Looping
−log10 P
Coaccesability
−log10 Q
1.10
1.15
1.20
1.25
Chromosome 19 (Mb)

(g)

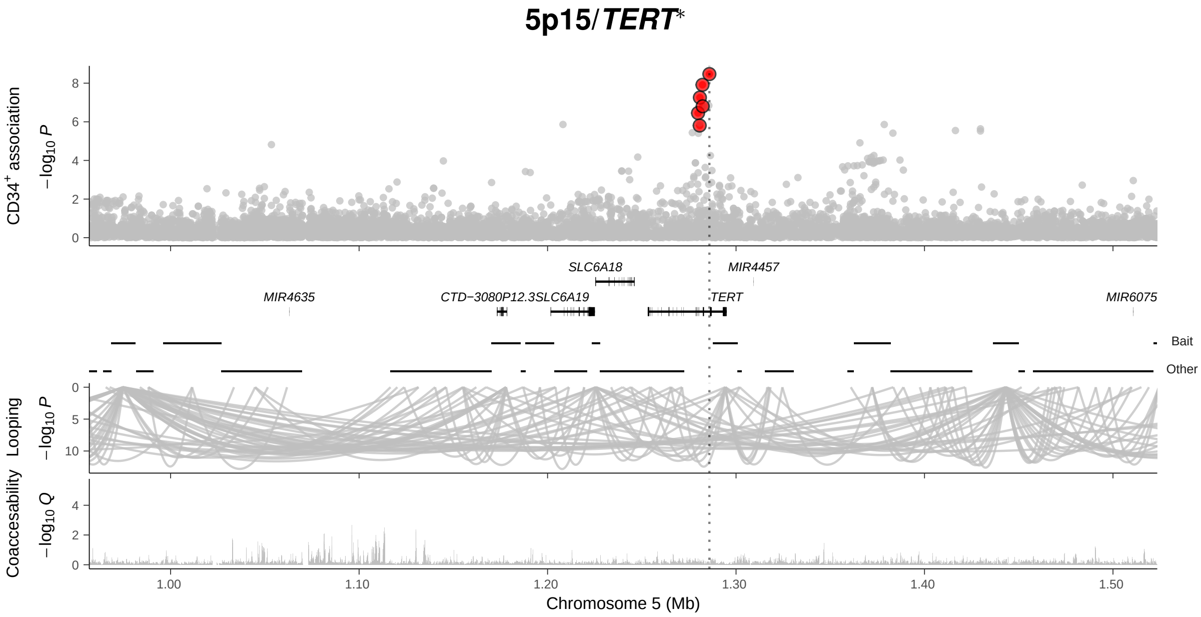
5p15/TERT*
CD34+ association
−log10 P
SLC6A18
MIR4457
MIR4635
CTD−3080P12.3
SLC6A19
TERT
MIR6075
Bait
Other
Looping
−log10 P
Coaccesability
−log10 Q
1.00
1.10
1.20
1.30
1.40
1.50
Chromosome 5 (Mb)

(h)

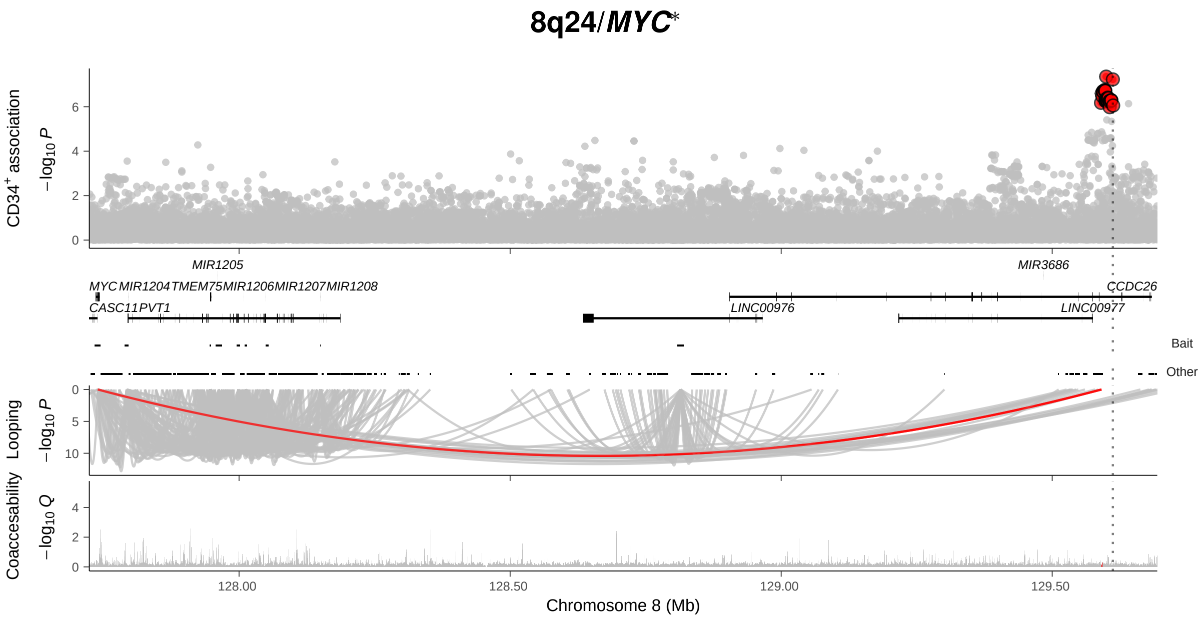

8q24/MYC*
CD34+ association
−log10 P
MIR1205
MIR3686
MYC MIR1204 TMEM75 MIR1206 MIR1207 MIR1208
CCDC26
CASC11 PVT1
LINC00976
LINC00977
Bait
Other
Looping
−log10 P
Coaccesability
−log10 Q
128.00
128.50
129.00
129.50
Chromosome 8 (Mb)

#### Supplementary Figure 7

Single-cell mRNA sequencing data for 35,582 mononuclear cells from blood and bone marrow. Data from Granja *et al.* (Nature Biotechnology, 2019 Dec;37(12):1458-1465). Dimension-reduction using uniform manifold approximation and projection (UMAP). The x- and y-axes indicate the projection of the expression pattern of each cell on the first and second UMAP components, respectively. Out of our 11 candidate genes, 10 were represented in this data set. Data are Magic-imputed expression values, shown per individual cell (left) or per cluster (right).

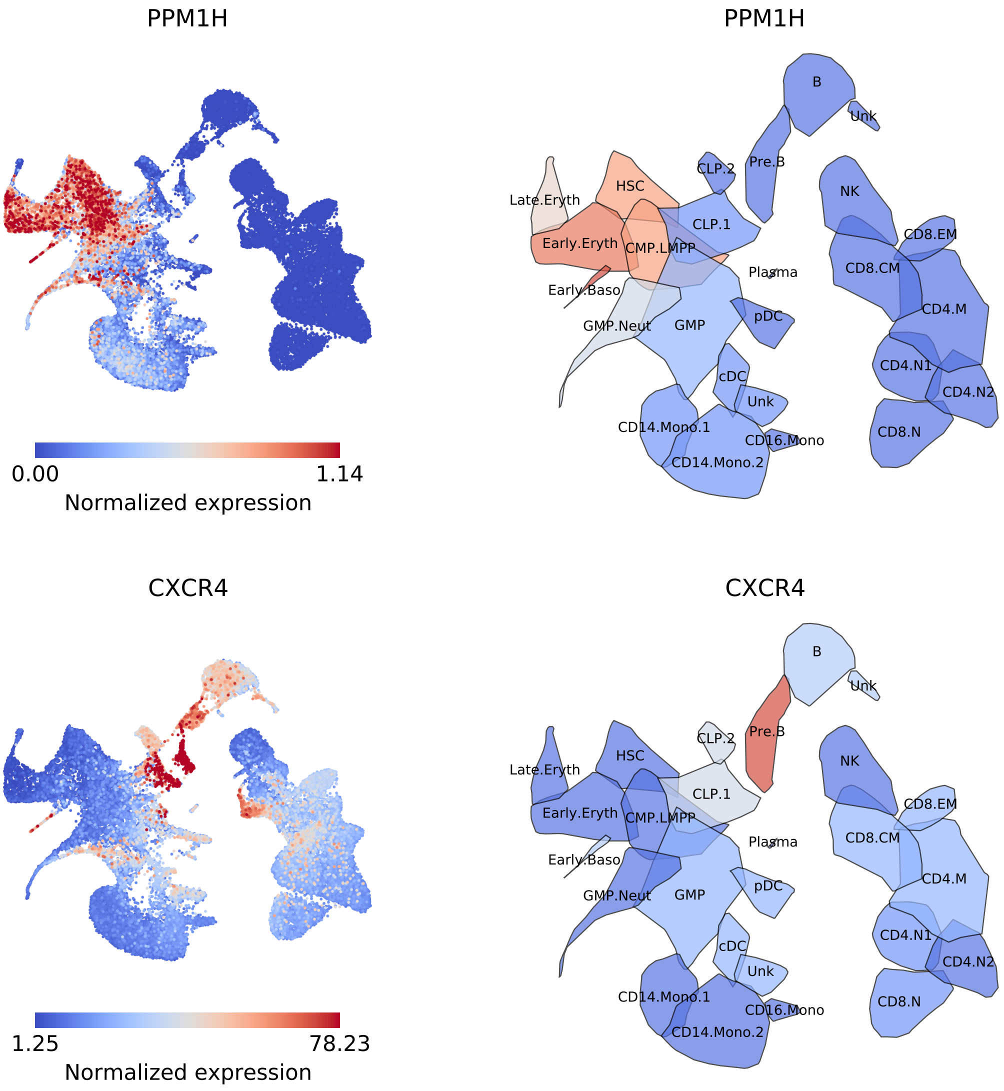

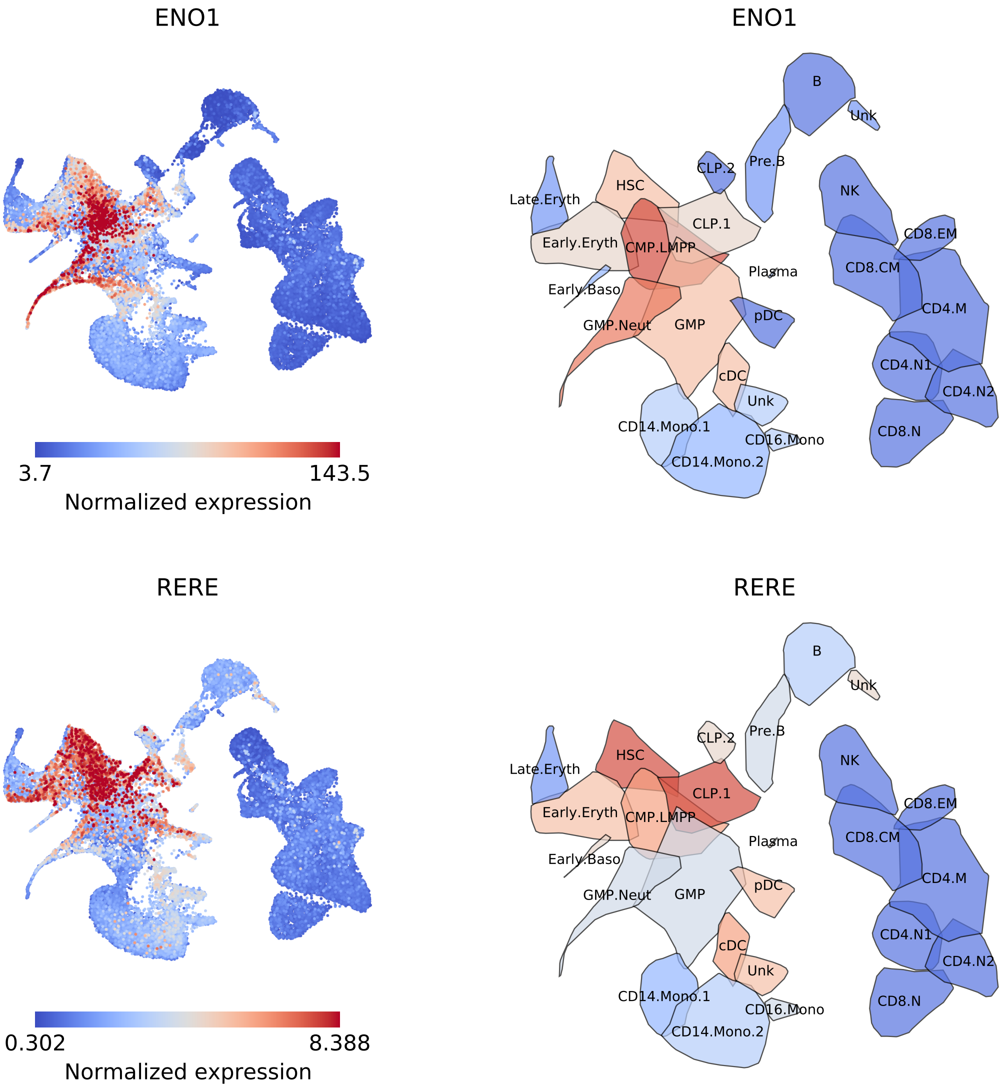
ENO1
3.7
143.5
Normalized expression
ENO1
B
Unk
CLP.2
Pre.B
NK
Late.Eryth
HSC
CLP.1
CD8.EM
Early.Eryth
CMP.LMPP
Plasma
CD8.CM
Early.Baso
CD4.M
GMP.Neut
GMP
pDC
CD4.N1
cDC
CD4.N2
Unk
CD14.Mono.1
CD16.Mono
CD8.N
CD14.Mono.2
RERE
0.302
8.388
Normalized expression
RERE
B
Unk
CLP.2
Pre.B
NK
Late.Eryth
HSC
CLP.1
CD8.EM
Early.Eryth
CMP.LMPP
Plasma
CD8.CM
Early.Baso
CD4.M
GMP.Neut
GMP
pDC
CD4.N1
cDC
CD4.N2
Unk
CD14.Mono.1
CD16.Mono
CD8.N
CD14.Mono.2

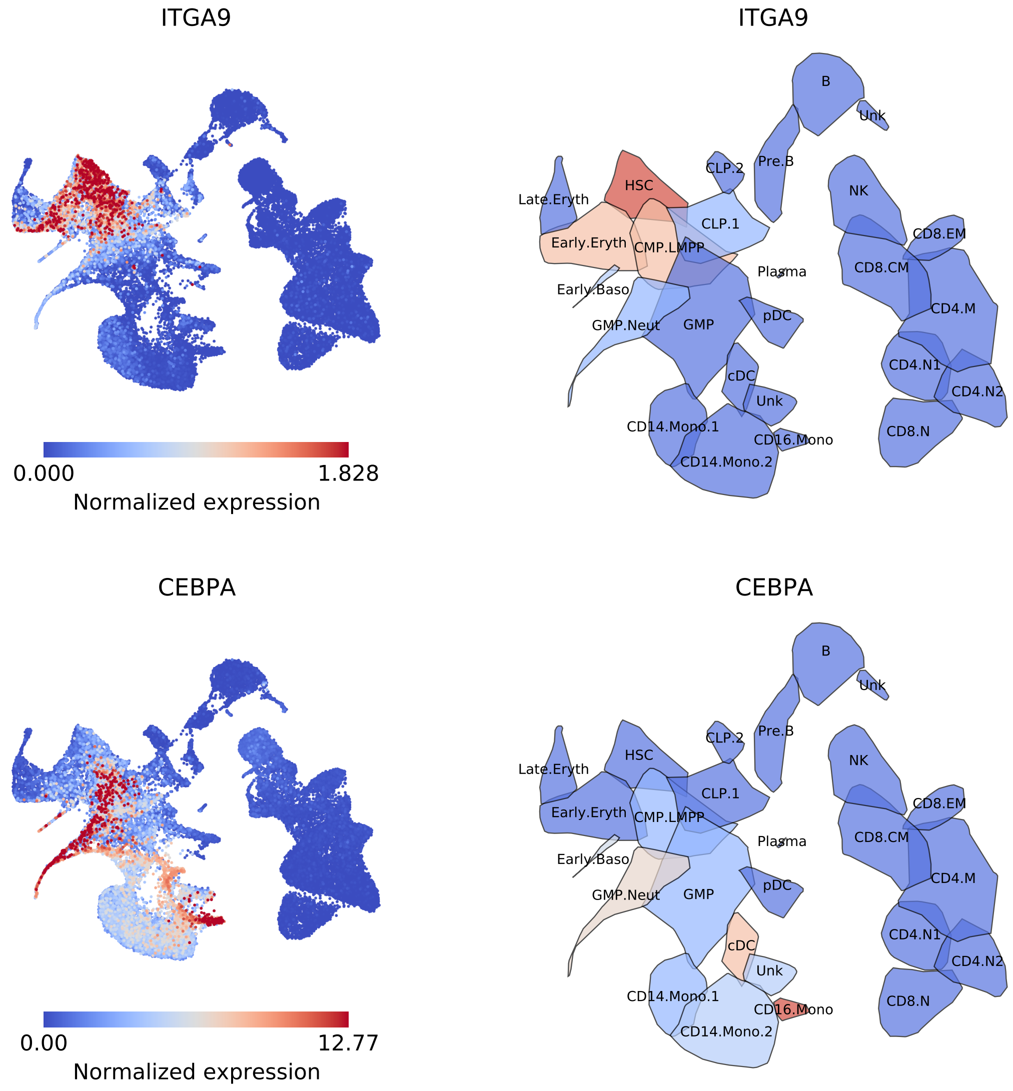
ITGA9
0.000
1.828
Normalized expression
ITGA9
B
Unk
CLP.2
Pre.B
HSC
Late.Eryth
NK
CLP.1
Early.Eryth
CMP.LMPP
CD8.EM
Plasma
CD8.CM
Early.Baso
CD4.M
pDC
GMP.Neut
GMP
CD4.N1
cDC
CD4.N2
Unk
CD14.Mono.1
CD8.N
CD16.Mono
CD14.Mono.2
CEBPA
0.00
12.77
Normalized expression
CEBPA
B
Unk
CLP.2
Pre.B
HSC
Late.Eryth
NK
CLP.1
Early.Eryth
CMP.LMPP
CD8.EM
Plasma
CD8.CM
Early.Baso
CD4.M
pDC
GMP.Neut
GMP
CD4.N1
cDC
CD4.N2
Unk
CD14.Mono.1
CD8.N
CD16.Mono
CD14.Mono.2

#### Supplementary Figure 8

Expression of candidate genes at the significant loci across hematopoietic blood cell types isolated by fluorescence-activated cell sorting. Data are $\log_2$-transformed, median-centered bulk mRNA sequencing data from Ulirsch *et al.*. (*Nature Genetics* 2019 Apr 51(4):683-693).

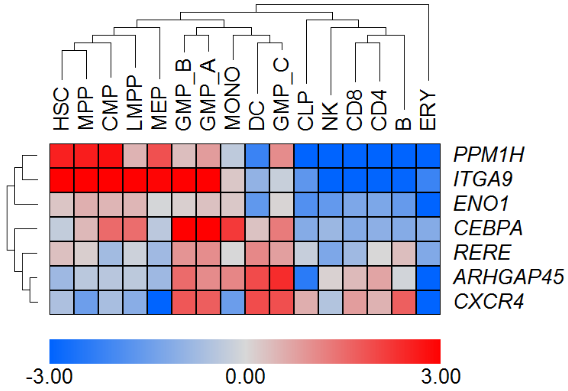

#### Supplementary Figure 9

Single-cell mRNA sequencing data for 4,905 CD34$^{+}$ cells from adult bone marrow. Dimension-reduction using uniform manifold approximation and projection (UMAP). x- and y-axes indicate projection of cell expression patterns on the first and second UMAP components, respectively. Data are Magic-imputed expression values, shown per individual cell (left) or per cluster (right).

Cluster descriptions: **HSC-I and HSC-II:** These two HSC clusters combined constituted more than 60% of surface marker defined HSC. **HSC-III:** This HSC cluster had lowest MEP (see below) fate potential. This cluster constituted only 10-15% of antibody-derived tag (ADT)-defined HSCs. **MPP-I:** Multi-potential progenitors I. This is the most primitive MPP cluster. About 90% ADT-defined MPPs consisted of HSCI-III and MPPI-III. MPP-I had highest representation in the ADT-defined MPP population (about 20%). **MPP-II:** Multi-potential progenitors I. This cluster is biased towards MEP fate (see MEP-I). **MPP-III:** Multi-potential progenitors III. This MPP cluster is biased towards GMP (see GMP) fate. **Ly-I:** Lymphoid I. More than 60% of ADT-defined LMPP cells are present in this cluster. **Ly-II:** Lymphoid II. This cluster contains about 70% of ADT-defined CLPs. **GMP:** Granulocyte-Monocyte progenitors. This cluster contains more than 50% of ADT-defined GMP and CMP cells. **MB:** Bone marrow mast cell/basophil cluster. **DC-I:** Dendritic cell progenitors. **DC-II:** Dendritic cell progenitors further towards DC fate than DC-I. **MEP-I:** Early Megakaryote-Erythroid progenitors. This cluster contains more than 50% of ADT-defined MEPs. **MEP-II:** This cluster is composed of cells that have further lineage-committed than MEP-I. **Cyc-I and II:** These clusters are cycling cells that are not in G1 phase.

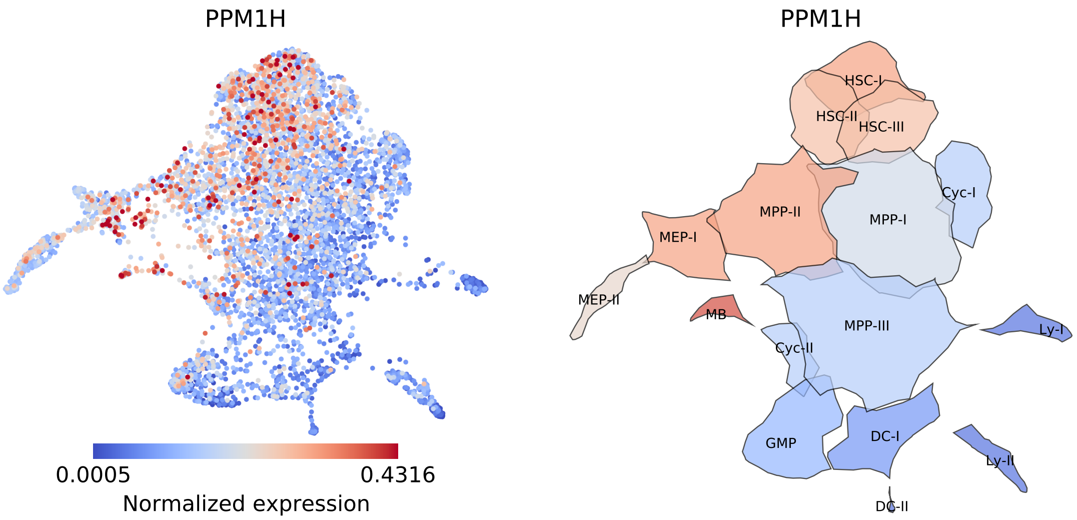

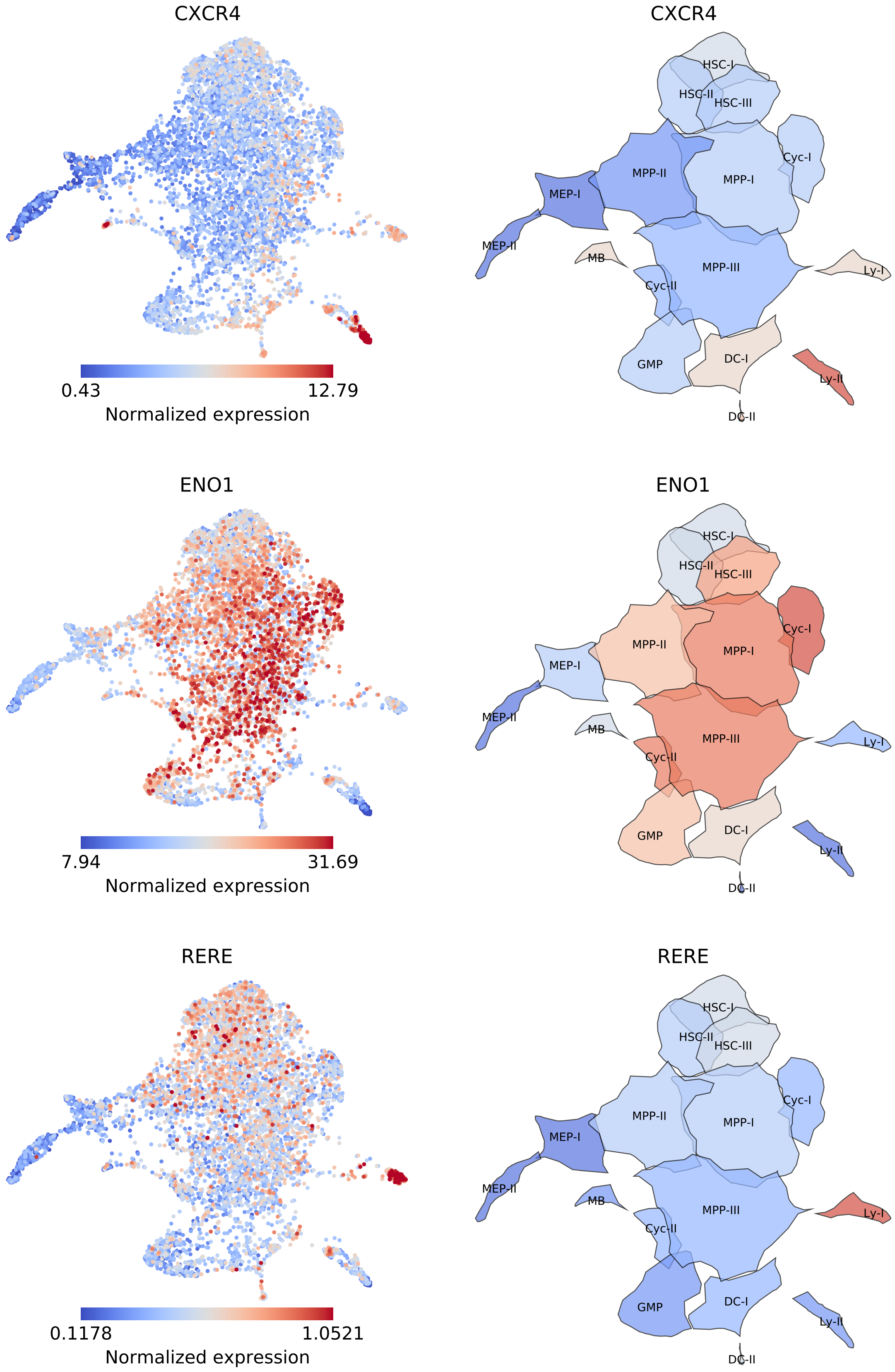
CXCR4
0.43
12.79
Normalized expression
CXCR4
HSC-I
HSC-II
HSC-III
Cyc-I
MPP-II
MPP-I
MEP-I
MEP-II
MB
MPP-III
Ly-I
Cyc-II
GMP
DC-I
Ly-II
DC-II
ENO1
7.94
31.69
Normalized expression
ENO1
HSC-I
HSC-II
HSC-III
Cyc-I
MPP-II
MPP-I
MEP-I
MEP-II
MB
MPP-III
Ly-I
Cyc-II
GMP
DC-I
Ly-II
DC-II
RERE
0.1178
1.0521
Normalized expression
RERE
HSC-I
HSC-II
HSC-III
Cyc-I
MPP-II
MPP-I
MEP-I
MEP-II
MB
MPP-III
Ly-I
Cyc-II
GMP
DC-I
Ly-II
DC-II

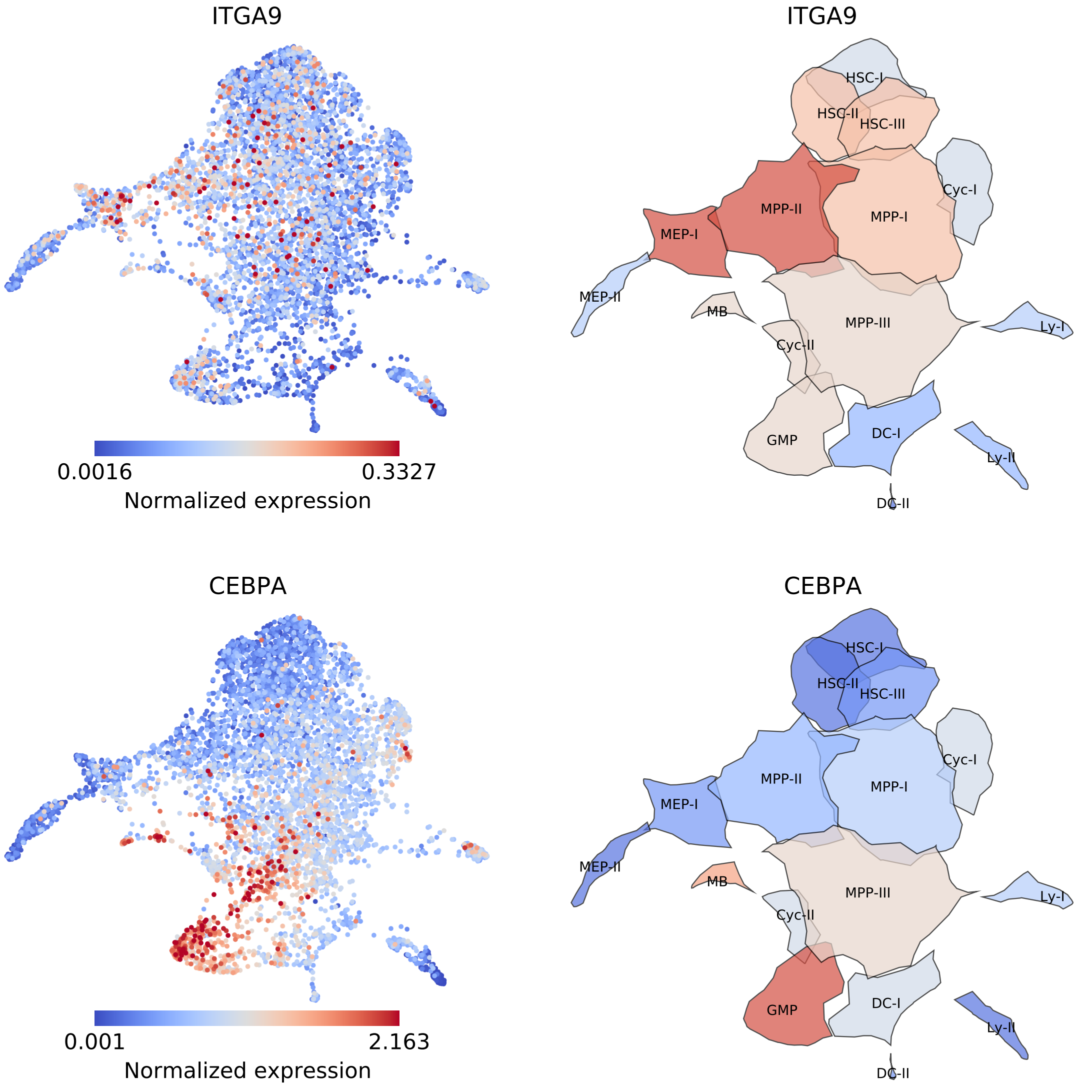
ITGA9
0.0016
0.3327
Normalized expression
ITGA9
HSC-I
HSC-II
HSC-III
Cyc-I
MPP-II
MPP-I
MEP-I
MEP-II
MB
MPP-III
Ly-I
Cyc-II
GMP
DC-I
Ly-II
DC-II
CEBPA
0.001
2.163
Normalized expression
CEBPA
HSC-I
HSC-II
HSC-III
Cyc-I
MPP-II
MPP-I
MEP-I
MEP-II
MB
MPP-III
Ly-I
Cyc-II
GMP
DC-I
Ly-II
DC-II

### Supplementary Figure 10

Expression of the 11 candidate genes in single-cell CITE-seq data for $CD34^+$ bone marrow cells stained with 44 nucleotide-tagged antibodies towards classical hematopoietic cell surface markers. From 4,905 input cells, we gated 2,887 cells belonging to the following $Lin^-CD34^+$ HSPC subsets: HSC ($CD38^-45RA^-90^+$; n = 501), MPP ($38^-45RA^-90^-$; n = 304), MEP ($38^+45RA^-123^-10^-$; n = 515), CMP ($38^+45RA^-123^+10^-$; n = 994), GMP ($38^+45RA^+123^+10^-$; n = 501) and LMPP ($38^-45RA^+90^-$; n = 72) subpopulations. Data are Magic-imputed expression values per gated cell. The data are median-summarized in **Fig. 3e**.

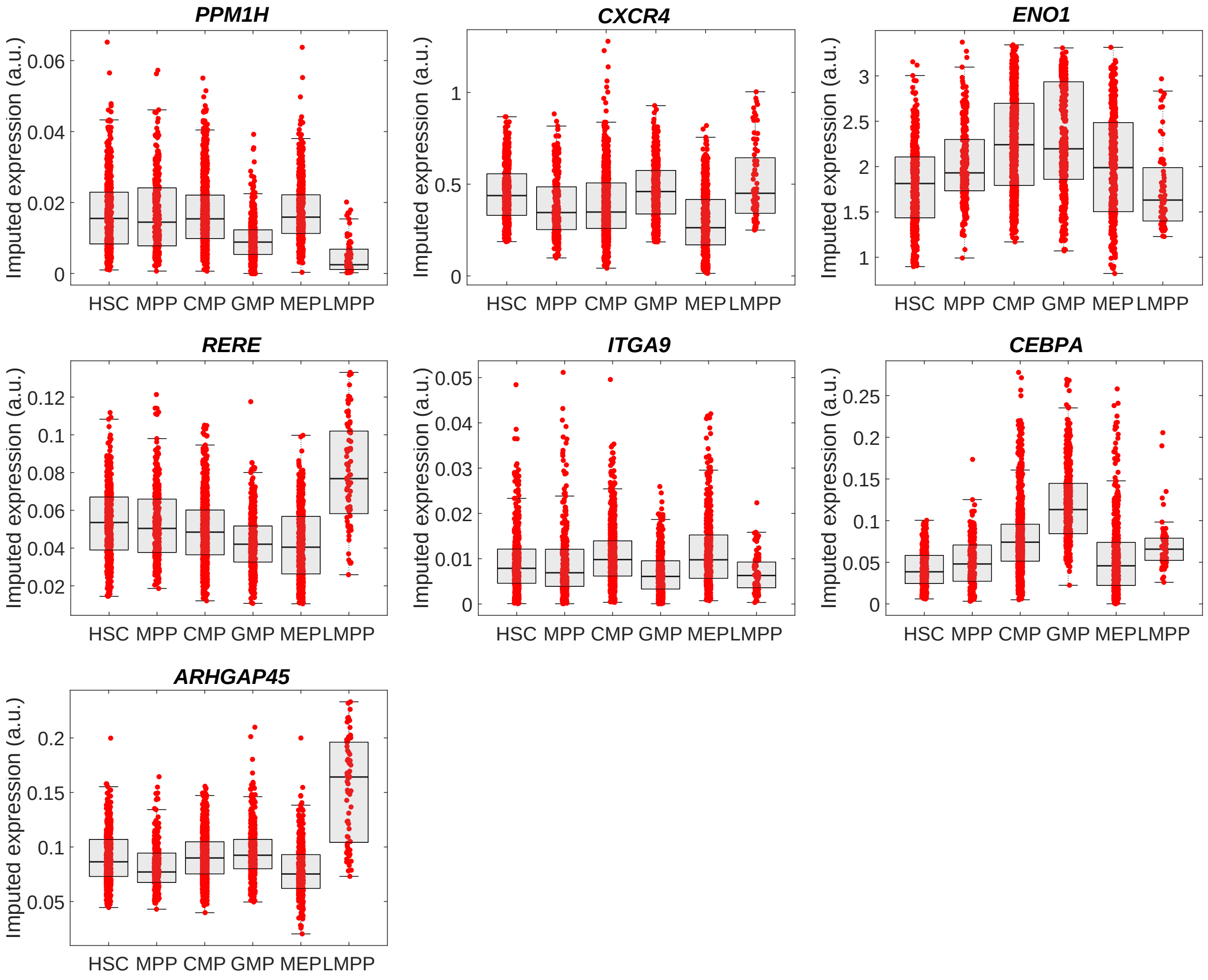

#### Supplementary Figure 11

Agarose gels confirming successful deletion of genomic DNA by dual-sgRNA CRISPR/Cas9 at the sites of the four putative causal variants at *CXCR4* on chromosome 2: rs309137 (**V1**), rs59222832 (**V2**), rs770321415 (**V3**), rs10193623 (**V4**) (red boxes). The percentages indicate deletion efficiency, calculated as intensity of the deletion band (red boxes) divided by the sum of the intensities of the deletion and wild type bands (blue boxes). As control, we used a sgRNA pair targeting a randomly selected region on a different chromosome, in this case an intronic region in the *WAC* gene on chromosome 10. The blue transparent areas are other PCR products that were run simultaneously in the gels. The latter are not a part of the data in **Fig. 3d**, and are included here for the sole purpose of showing the original gels.

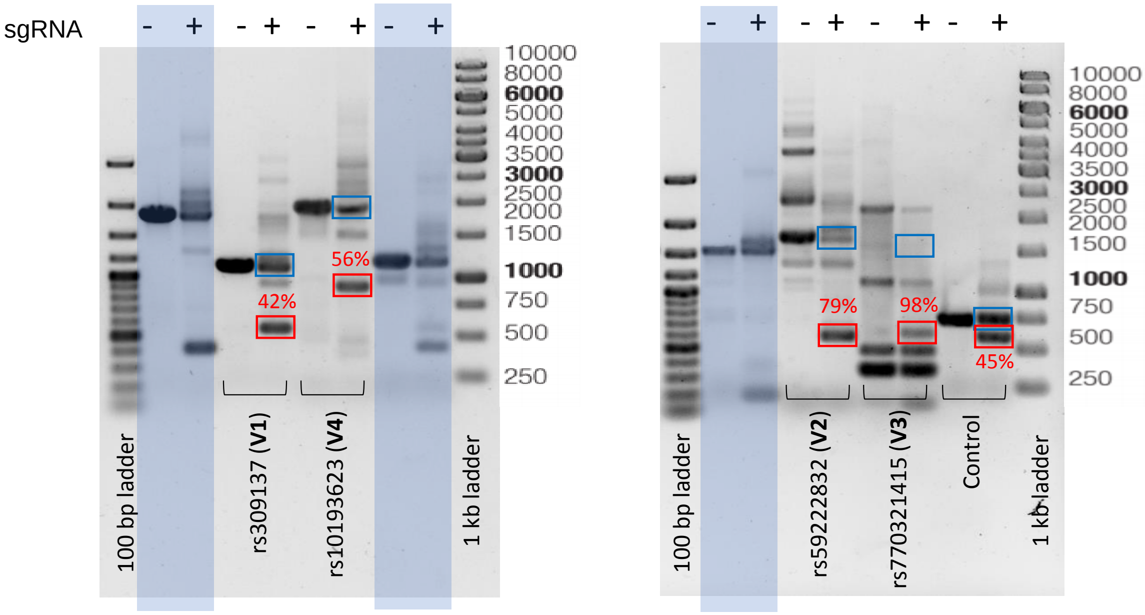

### Supplementary Figure 12

ATAC-seq data for the MOLM13 cell line around the four putative causal variants at *CXCR4*. Black bars indicate regions deleted by use of dual-sgRNA CRISPR/Cas9. The accessibility patterns are largely consistent with the ones observed in the same regions in sorted HSPCs (**Fig. 3b).**

V1: rs309137

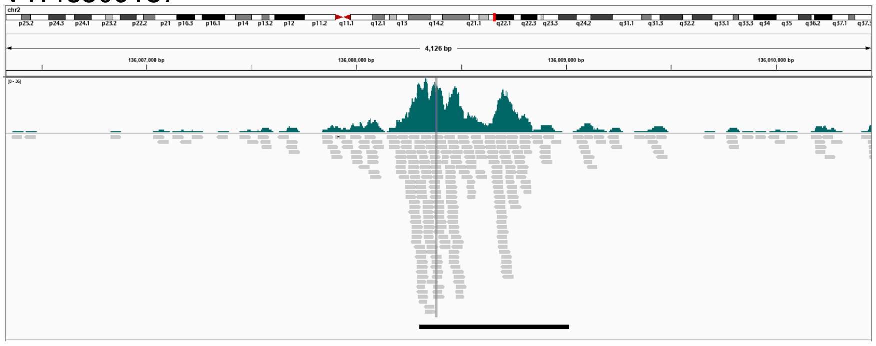


V2: rs59222832

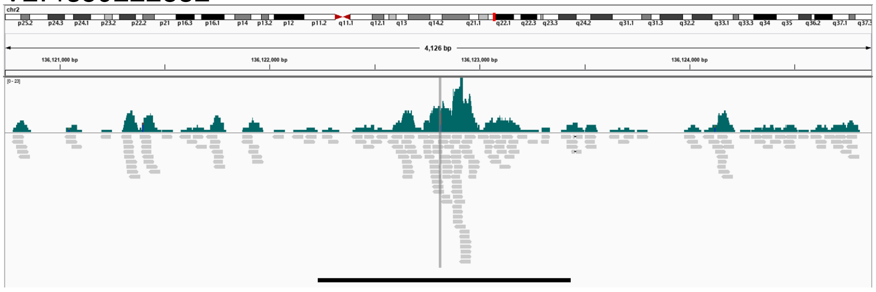


V3: rs770321415

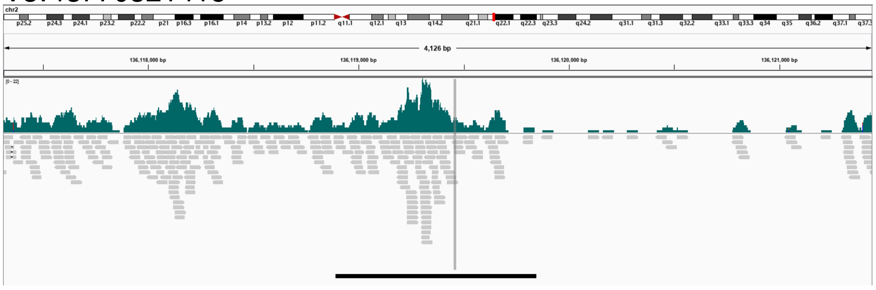


V4: rs10193623

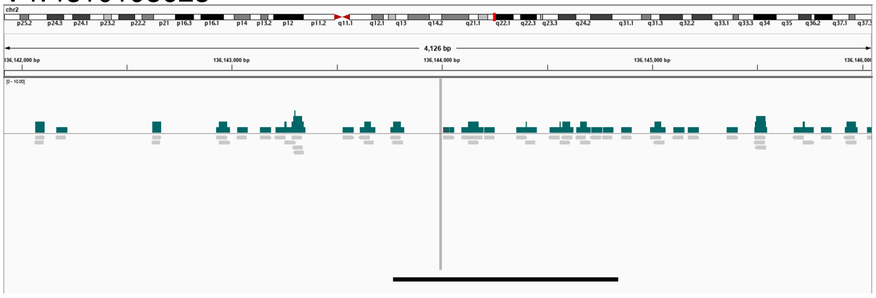

### Supplementary Figure 13

Sequence reads for Jurkat cells aligning to the rs772557 region: **(a)** whole-genome sequencing (WGS) data, showing heterozygosity for rs772557; and **(b)** MYB ChIP-sequencing data, showing exclusive pull-down of reads harboring rs772557-G. Data from Mansour MR *et al.*, *Science* 2014 Dec 12;346(6215):1373-7. Red reads harbor rs772557-G. Blue reads harbor rs772557-A. Grey reads do not overlap rs772557 or any other variant in the *PPM1H* credible set. The vertical grey bar indicates the position of rs772557.

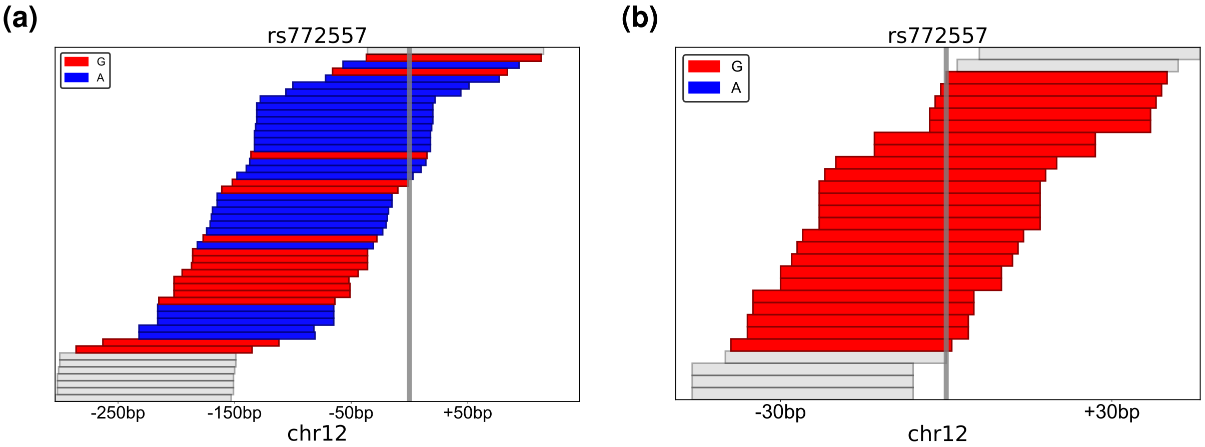

#### Supplementary Figure 14

Structure of sgRNAs used to perturb rs772557. Critical MYB recognition bases underlined.

5’ - CATTTTATCTGAAGAATCAGACCAAGC - C[A/G]TTGGAGCACCAGCC - 3'

sgRNA

rs772557

- Cas9 cut site

#### Supplementary Figure 15

To identify effects of rs772557 on specific CD34$^+$ populations, we calculated the correlation between rs772557 genotype and the expression of all genes with median FPKM > 1.0 in our CD34$^+$ mRNA-sequencing data set for blood donors. We then tested for enrichment for correlations within sets of marker genes for different CD34$^+$ HSPC subpopulations, inferred by comparing the expression profile of each cell type to other CD34$^+$ cell types in three data sets: **(a)** bulk RNA-sequencing data (Ulirsch *et al.*, *Nature Genetics* 2019 Apr;51(4):683-693); **(b)** single-cell RNA-sequencing data for CD34$^+$ cells (**Online Methods**), or **(c)** single-cell RNA-sequencing data for mononuclear cells (Granja *et al.*, *Nature Biotechnology* 2019 Dec;37(12):1458-1465.). For analysis we used RenderCat (Nilsson *et al.*, *Genome Biology*, 2007;8(5):R74) with default settings (Zhang-C statistic). For completeness, we carried out the analysis with varying numbers of marker genes, with similar results. The results shown here were obtained with the top 100 and 250 marker genes.

#### Bulk RNA-sequencing sort cells (Ulirsch *et al.*), 100 marker genes

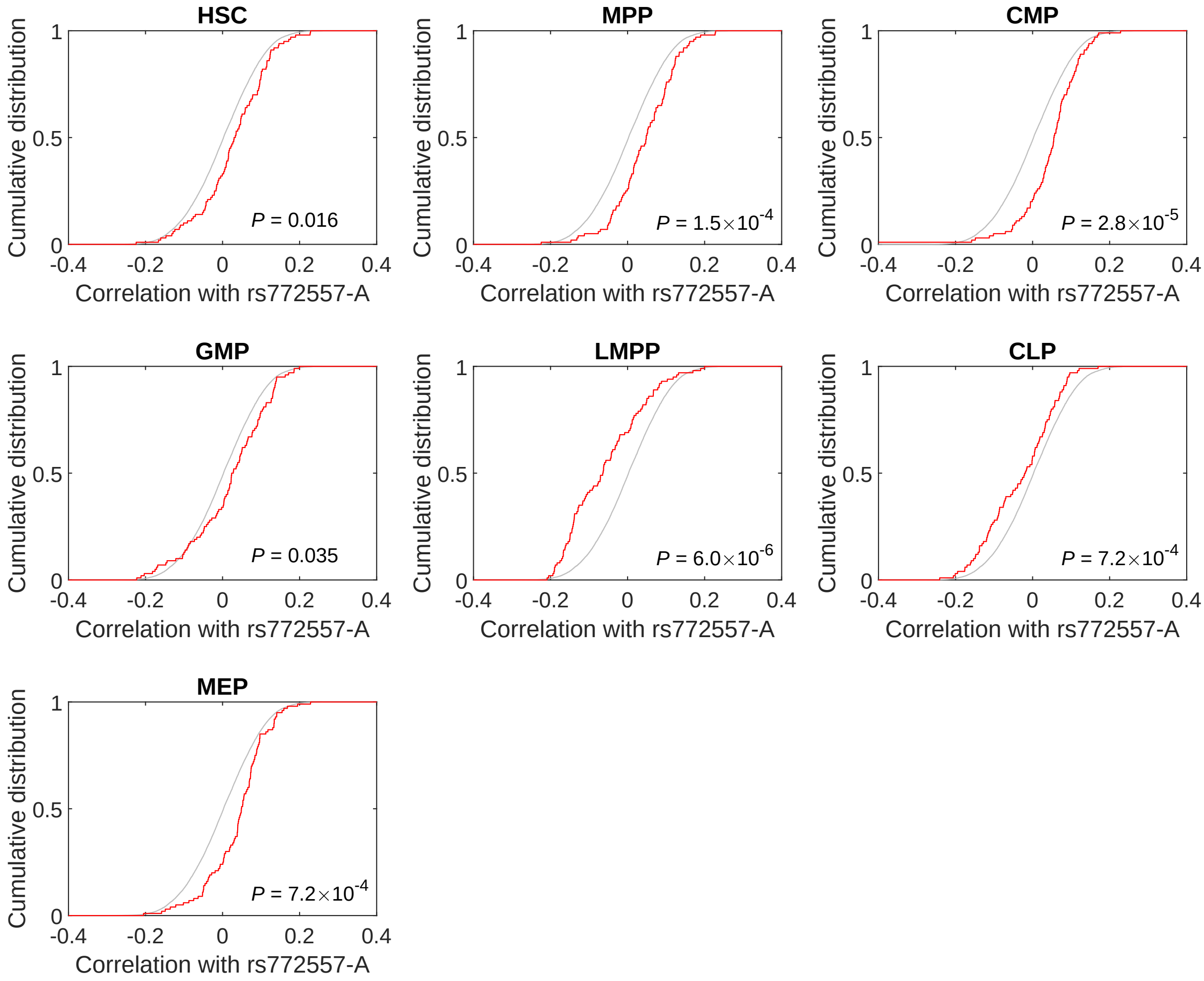

#### Bulk RNA-sequencing sort cells (Ulirsch *et al.*), 250 marker genes

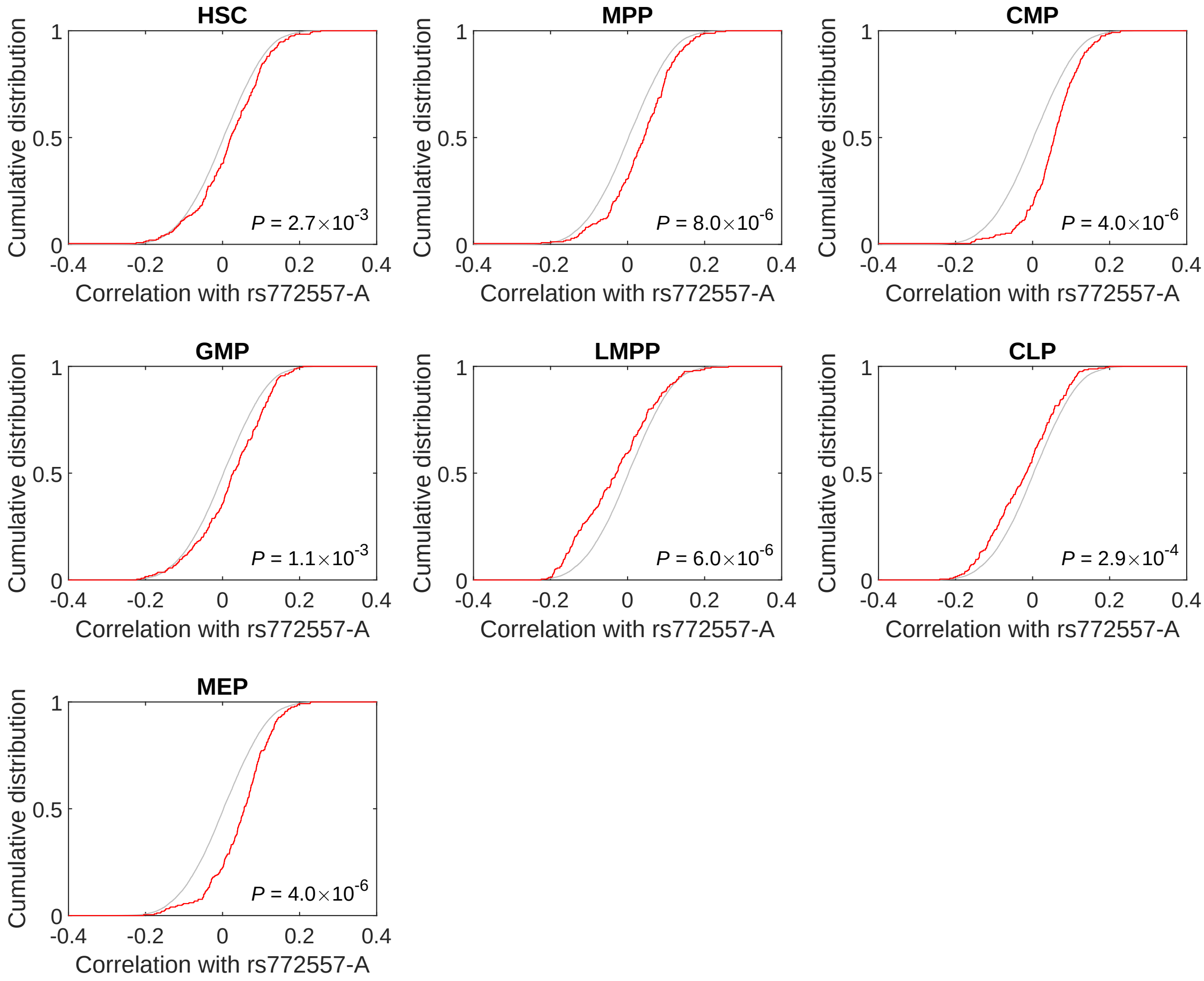

#### Single-cell RNA-sequencing CD34$^+$ cells, 100 marker genes

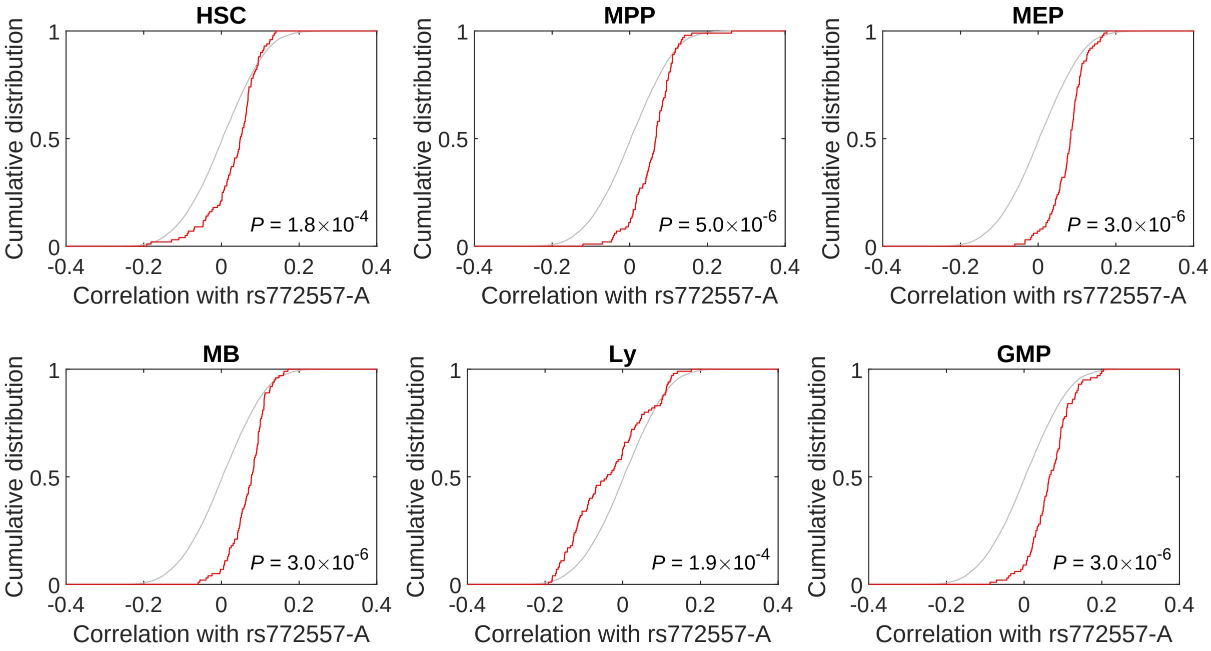

#### Single-cell RNA-sequencing CD34$^{+}$ cells, 250 marker genes

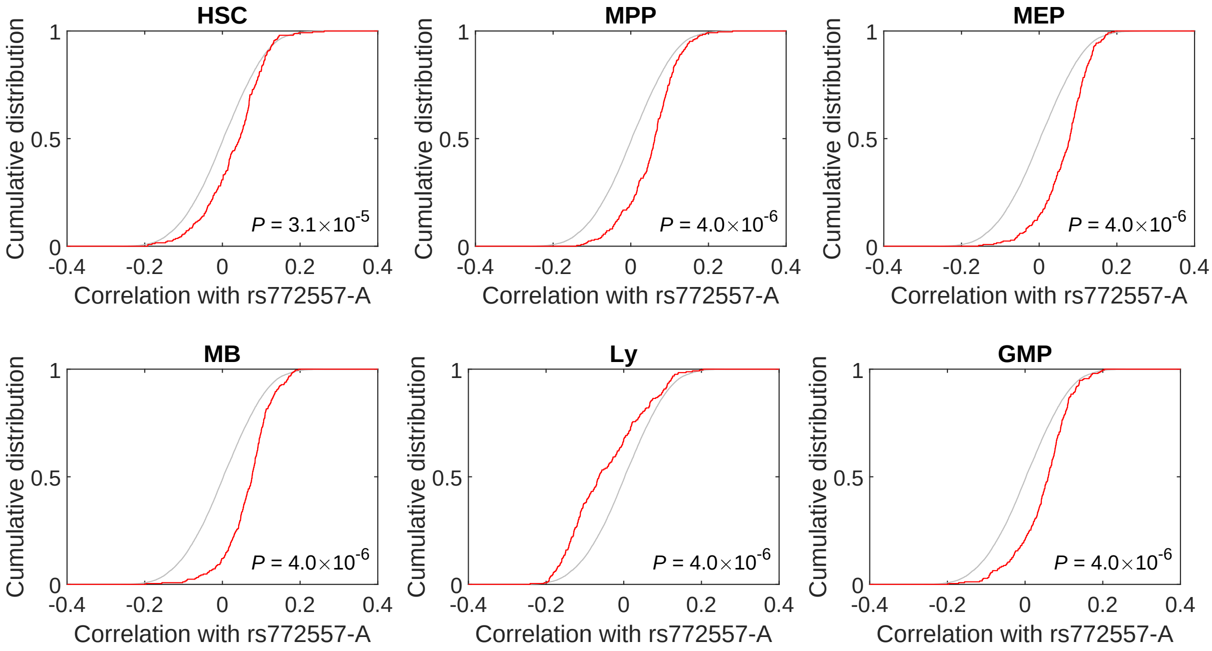

#### Single-cell RNA-sequencing mononuclear cells (Granja *et al.*), 100 marker genes

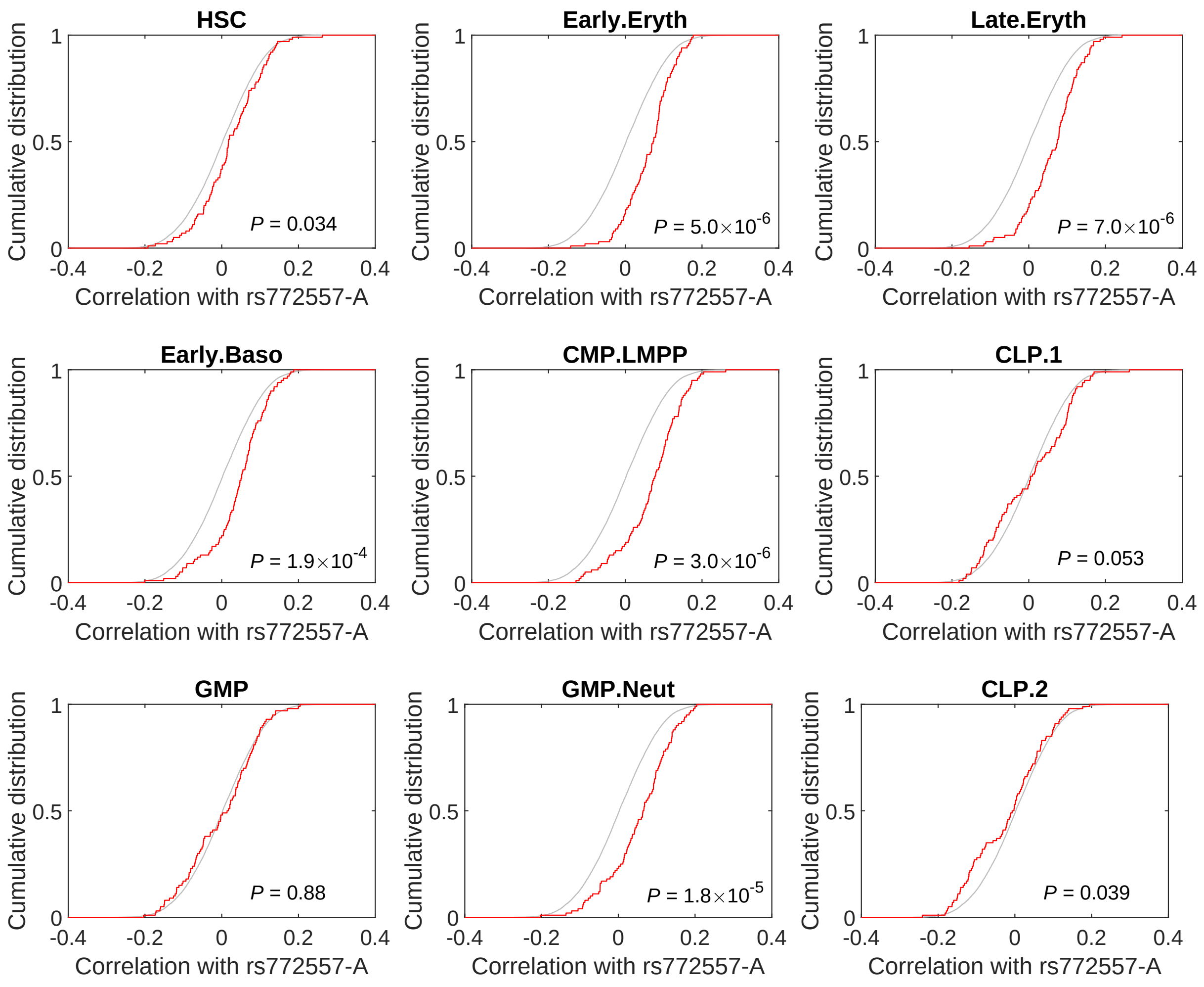

#### Single-cell RNA-sequencing mononuclear cells (Granja *et al.*), 250 marker genes

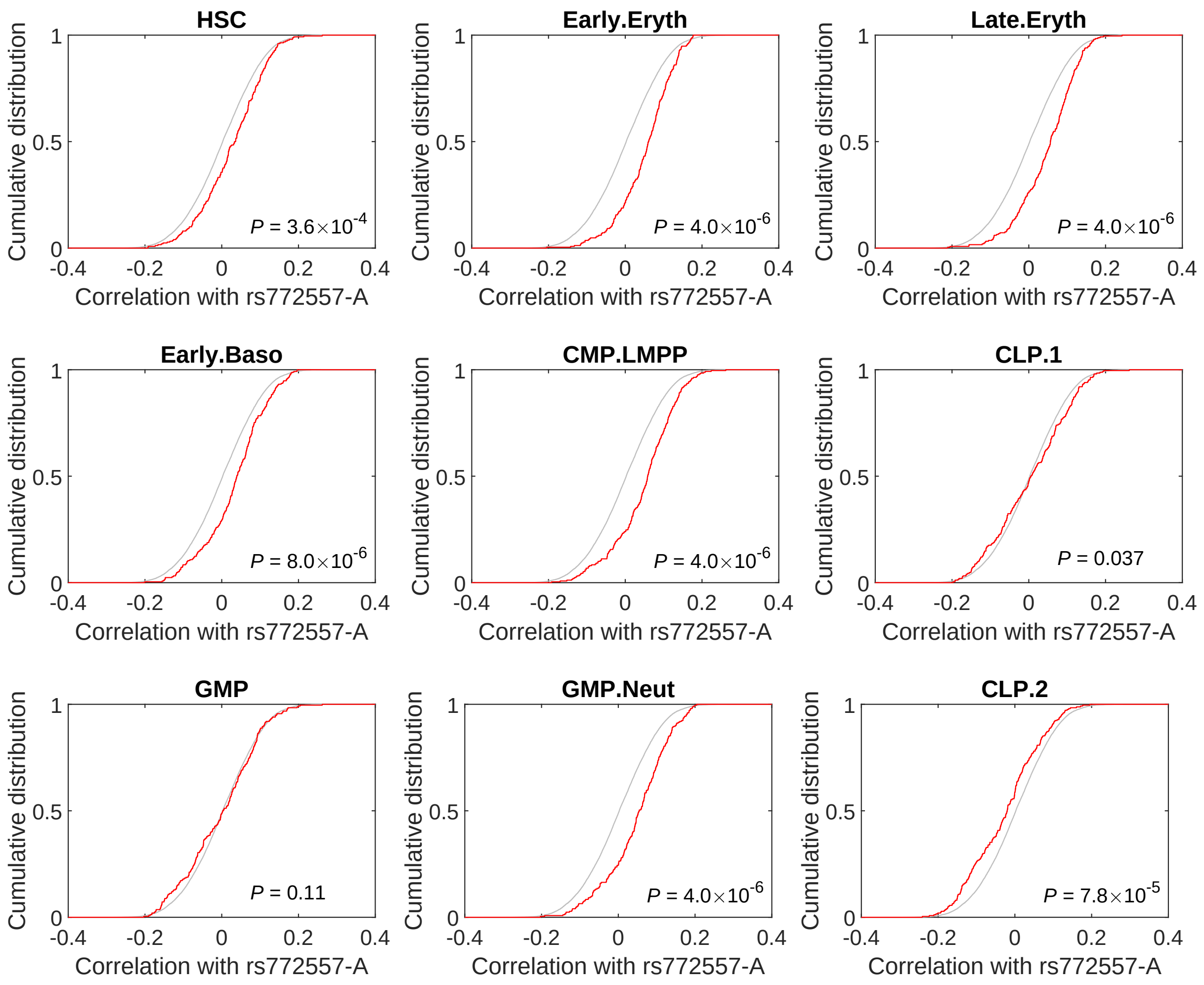

### Supplementary Figure 16

Gating of cord blood samples. After excluding dead cells using 7-AAD, Ficoll-enriched mononuclear cells were gated on forward (FSC-A) and side scatter area (SSC-A). After doublet exclusion based on FSC-A and forward scatter height (FSC-H), we gated CD34$^{+}$CD45$^{\text{low}}$ cells, negative for lineage markers CD3, CD19, CD14, CD16, and CD56. These cells were divided into CD38$^{-}$ and CD38$^{+}$. Among CD38$^{+}$ cells, we defined CD45RA$^{+}$CD10$^{+}$ cells as B-NK progenitors, while CD10$^{-}$ cells were subdivided into common myeloid progenitors (CMP), granulocyte-monocyte progenitors (GMP) or megakaryocyte-erythrocyte progenitors (MEP) based on CD135 and CD45RA expression. CD38$^{-}$ were defined as hematopoietic stem cells (HSC), multipotent progenitors (MPP) and multilymphoid progenitors (MLP) based on CD90 and CD45RA expression. Finally, MLP were subdivided into CD10$^{+}$ and CD10$^{-}$. For each of the Lin$^{-}$CD34$^{+}$ populations, we calculated their ratio out of total CD34$^{+}$CD45$^{\text{low}}$, and out of Lin$^{-}$CD34$^{+}$CD45$^{\text{low}}$.

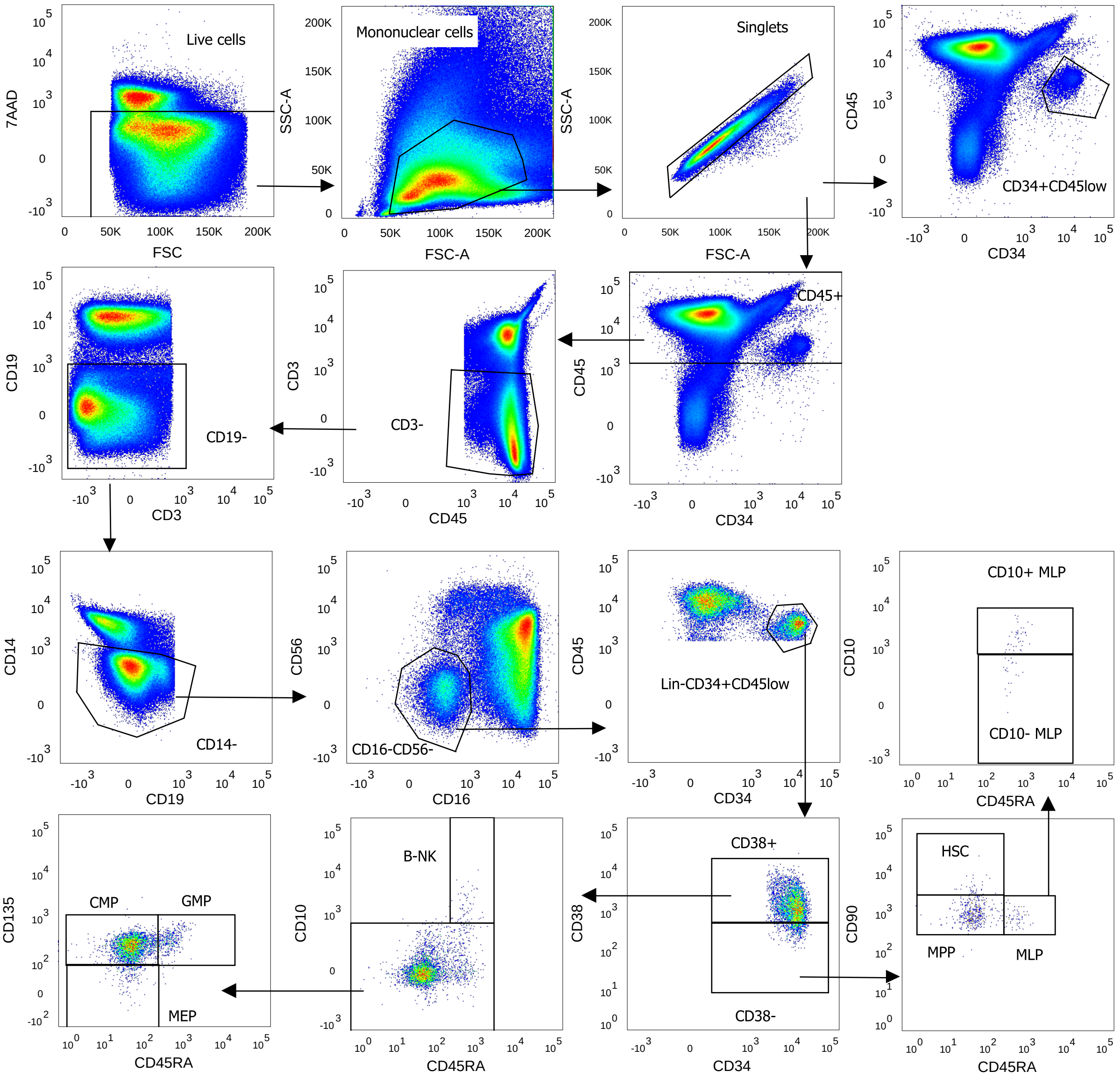

#### Supplementary Figure 17

We screened three shRNA sequences in the leukemia cell line K562. One sequence (sh1) yielded deep knockdown. Expression values are $2^{-\Delta\Delta Ct}$ quantitative PCR readings normalized to shCtrl.

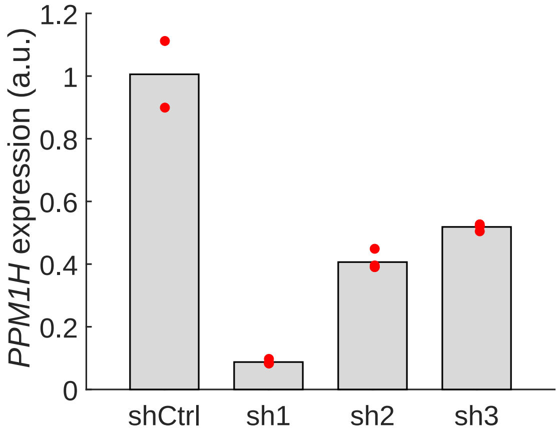

#### Supplementary Figure 18

To test further the effects of the *ITGA9* variant, we quantified ITGA9 protein expression on CD34$^+$ cells. Consistent with the *cis*-eQTL, we identified a significant correlation between rs17227369 genotype (credible set proxy for rs201494641 used for TaqMan genotyping) and median fluorescence intensity (MFI), as quantified using phycoerythrin (PE)-conjugated monoclonal antibody towards ITGA9. Statistics are for Pearson correlation.

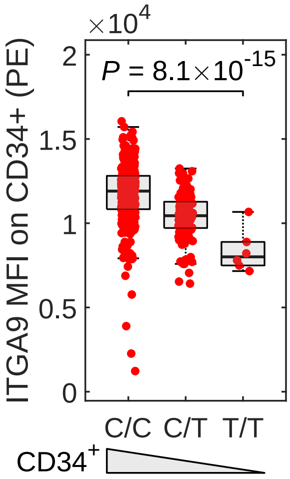

#### Supplementary Figure 19

To validate the AliGater software, we gated 2,838 samples both with AliGater and manually. This plot shows the correlation between manual and AliGater-assisted quantification of $CD34^+$ levels in these samples. The *r* and *P*-values are for Spearman correlation.

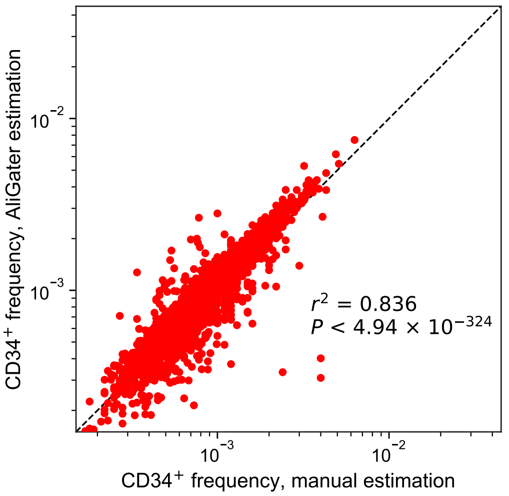

### Supplementary Table 1

Using blood samples from blood donors and primary care patients from Skåne Country, Sweden, we generated three association data sets (denoted Phase I, II and III). Out of 16,931 collected samples, a total of 13,167 samples from unique participants yielded high-quality phenotype and genotype data. Through ancestry analysis, we identified individuals of Swedish and non-Swedish European origin based on the genotype data. For discovery, we used the Swedish samples. For follow-up, non-Swedish European samples. These tables list the numbers of unique individuals in each sample set.

| Swedes (discovery) | | | | |
|---|---|---|---|---|
| **Phase** | **N** | **Male (%)** | **Age (years±SD)** | **Blood donors (%)** |
| I | 2 408 | 59,5 | 40.9 (±14.3) | 100,0 |
| II | 3 826 | 42,4 | 45.1 (±14.8) | 30,0 |
| III | 4 715 | 43,4 | 46.3 (±14.7) | 34,1 |
| Total | 10 949 | 45,1 | 45.1 (±14.8) | 47,2 |

| Caucasian non-Swedes (follow-up) | | | | |
|---|---|---|---|---|
| **Phase** | **N** | **Male (%)** | **Age (±SD)** | **Blood donors (%)** |
| I | 344 | 59,3 | 39.4 (±14.2) | 100,0 |
| II | 753 | 35,9 | 43 (±14.2) | 12,1 |
| III | 1 122 | 40,4 | 43 (±14.2) | 40,9 |
| Total | 2 218 | 41,8 | 43 (±14.2) | 33,5 |

| All | | | | |
|---|---|---|---|---|
| **Phase** | **N** | **Male (%)** | **Age (±SD)** | **Blood donors (%)** |
| I | 2 752 | 59,5 | 44.9 (±14.7) | 100,0 |
| II | 4 578 | 41,3 | 44.8 (±14.7) | 27,0 |
| III | 5 837 | 42,9 | 44.8 (±14.7) | 32,9 |
| Total | 13 167 | 45,8 | 44.8 (±14.7) | 44,9 |

### Supplementary Table 2

Number of cells analyzed per sample (median, 10th and 90th percentile for each collection phase).

| Phase | No. of input cells | | | No. of $CD45^+$ PBMCs | | | No. of $CD34^+$ PBMCs | | | $CD34^+$ level | | |
|---|---|---|---|---|---|---|---|---|---|---|---|---|
| | median | 10th | 90th | median | 10th | 90th | median | 10th | 90th | median | 10th | 90th |
| I | 500000 | 292444 | 500000 | 160923 | 100619 | 215479 | 142 | 59 | 281 | 9.11E-04 | 4.00E-04 | 1.82E-03 |
| II | 457150 | 197285 | 500000 | 136951 | 63134 | 201621 | 124 | 52 | 254 | 9.66E-04 | 4.50E-04 | 1.99E-03 |
| III | 1000000 | 934300 | 1000000 | 326804 | 214426 | 437000 | 354 | 178 | 702 | 1.00E-03 | 5.30E-04 | 2.10E-03 |

#### Supplementary Table 3

Lead variants for the 9 significant and two suggestive association signals with blood CD34$^+$ levels, along with their statistics.

| | | | | | | Discovery | | Follow-up | | Combined | | | |
|---|---|---|---|---|---|---|---|---|---|---|---|---|---|
| Position | rsID | Gene | OA | EA | EAF (%) | β | *P* | β | *P* | β | *P* | $P_{Bonferroni}$ | $P_{het}$ |
| chr1:8857756 | rs2047094 | *ENO1, RERE* | A | C | 49.2 | -0.093 | 4.3E-11 | -0.141 | 2.10E-06 | -0.102 | 1.3E-15 | 7.93E-09 | 0.15 |
| chr2:136008381 | rs309137 | *CXCR4* | T | C | 25.5 | -0.146 | 1.7E-15 | -0.068 | 1.10E-02 | -0.121 | 1.3E-15 | 7.93E-09 | 0.016 |
| chr2:136125862 | rs11688530 | *CXCR4* | G | A | 7.1 | 0.234 | 1.2E-15 | 0.228 | 1.10E-07 | 0.232 | 7.9E-22 | 4.82E-15 | 0.91 |
| chr2:136136760 | rs555647251 | *CXCR4* | C | T | 0.083 | 1.731 | 3.3E-10 | 1.794 | 1.00E-02 | 1.740 | 1.2E-11 | 7.32E-05 | 0.93 |
| chr2:136143990 | rs10193623 | *CXCR4* | TA | T,TT | 5.9 | -0.181 | 1.4E-08 | -0.235 | 2.30E-08 | -0.200 | 2.8E-15 | 1.71E-08 | 0.30 |
| chr3:37477922 | rs201494641 | *ITGA9* | TTT | T,TT,TTTT | 13.0 | -0.113 | 8.6E-08 | -0.158 | 7.80E-05 | -0.123 | 4.7E-11 | 2.87E-04 | 0.32 |
| chr12:62912352 | rs699585 | *PPM1H* | T | G | 46.1 | -0.121 | 1.2E-17 | -0.151 | 2.00E-07 | -0.126 | 2.2E-23 | 7.86E-16 | 0.35 |
| chr19:1079960 | rs36084354 | *ARHGAP45* | G | A | 8.8 | 0.140 | 6.1E-07 | 0.222 | 8.70E-03 | 0.148 | 2.7E-08 | 1.50E-02 | 0.36 |
| chr19:33264458 | rs12975577 | *CEBPA* | T | C | 49.0 | -0.090 | 1.8E-10 | -0.047 | 1.10E-01 | -0.082 | 1.2E-10 | 4.29E-03 | 0.18 |
| chr5:1285859 | rs7705526 | *TERT* | C | A | 34.0 | 0.089 | 3.4E-09 | 0.051 | 9.10E-02 | 0.081 | 1.5E-09 | 5.36E-02 | 0.26 |
| chr8:129599498 | rs60379390 | *CCDC26* | ! | ACTCTCTTTT | 36.9 | -0.08 | 4.4E-08 | -0.053 | 6.70E-02 | -0.075 | 1.1E-08 | 3.93E-01 | 0.42 |

**Abbreviations:** Other allele (OA); Effect Allele (EA); Effect allele frequency (EAF).

### Supplementary Table 4

Table of 99% credible sets of probable causal variants. Red lines represent variants that both: **(a)** map to a regulatory region with accessible chromatin in HSPCs (ATAC-seq column); and **(b)** map to a gene promoter or have a chromatin looping interaction with a gene promoter in CD34$^+$ cells (PCHi-C column). Using this two-criterion filter, we prioritize variants that likely regulate the expression in HSPCs. We identified such variants at *PPM1H* , *ENO1* , *ITGA9, CXCR4* , and *CCDC26* (at the latter a long-range interaction with *MYC* ).

| Position | rsID | EA | OA | EAF | β | *P* | postProb | ATAC-seq | PCHi-C |
|---|---|---|---|---|---|---|---|---|---|
| **1p36, *ENO1 RERE*** | | | | | | | | | |
| chr1:8857756 | rs2047094 | C | A | 49.22 | -0.102 | 1.3E-15 | 0.201 | | |
| chr1:8857040 | rs11121246 | ! | CTCG | 49.19 | -0.101 | 2.3E-15 | 0.317 | | *ENO1-AS1;RERE* |
| chr1:8857040 | . | CTCT | ! | 49.19 | -0.101 | 2.3E-15 | 0.432 | | *ENO1-AS1;RERE* |
| chr1:8857339 | rs59246144 | T | TTA | 49.19 | -0.101 | 2.3E-15 | 0.547 | | |
| chr1:8857610 | rs11590606 | C | T | 49.20 | -0.101 | 2.3E-15 | 0.662 | | |
| chr1:8858254 | rs10864368 | T | C | 49.20 | -0.101 | 2.6E-15 | 0.764 | Yes | |
| chr1:8856165 | rs11121245 | T | C | 49.23 | -0.100 | 3.6E-15 | 0.838 | | *ENO1-AS1;RERE* |
| chr1:8853596 | rs12752838 | ! | GGAAAAAAAAAAG | 49.03 | -0.099 | 7.5E-15 | 0.874 | | *ENO1-AS1;RERE* |
| chr1:8850047 | . | TAAAA | ! | 48.52 | -0.099 | 1.0E-14 | 0.902 | | *ENO1-AS1;RERE* |
| chr1:8850047 | rs6577536 | ! | TAAAG | 48.56 | -0.098 | 1.2E-14 | 0.924 | | *ENO1-AS1;RERE* |
| chr1:8850483 | rs6661010 | C | T | 48.57 | -0.098 | 1.3E-14 | 0.945 | | *ENO1-AS1;RERE* |
| chr1:8853423 | rs548497051 | ! | AA | 48.53 | -0.098 | 1.8E-14 | 0.961 | | *ENO1-AS1;RERE* |
| chr1:8848153 | rs2065603 | G | A | 48.41 | -0.098 | 2.3E-14 | 0.973 | Yes | *ENO1-AS1;RERE* |
| chr1:8851314 | . | CA | ! | 47.82 | -0.096 | 5.3E-14 | 0.978 | | *ENO1-AS1;RERE* |
| chr1:8851314 | rs34971118 | ! | CAA | 47.91 | -0.096 | 5.5E-14 | 0.983 | | *ENO1-AS1;RERE* |
| chr1:8853423 | . | AAAAAAAAAAAAAAAAA | ! | 48.18 | -0.096 | 6.2E-14 | 0.988 | | *ENO1-AS1;RERE* |
| chr1:8844149 | rs60454710 | ! | C | 47.99 | -0.096 | 6.7E-14 | 0.992 | | *RERE* |
| **2q22, *CXCR4* rs10193623** | | | | | | | | | |
| chr2:136143990 | . | ! | TA | 5.87 | -0.200 | 2.8E-15 | 0.399 | Yes | *CXCR4* |
| chr2:136143990 | rs10193623 | TT | ! | 5.87 | -0.200 | 2.8E-15 | 0.798 | Yes | *CXCR4* |
| chr2:136142899 | rs6726457 | T | C | 5.85 | -0.199 | 5.8E-15 | 0.995 | | |
| **2q22, *CXCR4* rs11688530** | | | | | | | | | |
| chr2:136125862 | rs11688530 | A | G | 7.12 | 0.232 | 7.9E-22 | 0.195 | | |
| chr2:136126891 | rs11679328 | T | C | 6.98 | 0.231 | 1.3E-21 | 0.316 | | |

| | | | | | | | | | |
|---|---|---|---|---|---|---|---|---|---|
| chr2:136122815 | rs59222832 | C | T | 6.99 | 0.231 | 1.7E-21 | 0.868 | Yes | In *CXCR4* promoter |
| chr2:136124535 | rs6751768 | T | C | 6.99 | 0.230 | 1.9E-21 | 0.951 | | |
| chr2:136098926 | rs61528333 | GTTTTTTTT | ! | 6.97 | 0.229 | 5.7E-21 | 0.979 | | *LCT* |
| chr2:136098926 | . | ! | GTTTTTTT | 6.98 | 0.227 | 1.3E-20 | 0.992 | | *LCT* |
| **2q22, *CXCR4* rs309137** | | | | | | | | | |
| chr2:136008381 | rs309137 | C | T | 25.53 | -0.121 | 1.3E-15 | 0.183 | Yes | *CXCR4* |
| chr2:135868032 | . | A | ! | 24.12 | -0.118 | 8.7E-15 | 0.212 | | |
| chr2:135873664 | rs3835799 | ACAGAGGTAC | A | 24.17 | -0.118 | 8.9E-15 | 0.240 | | |
| chr2:135879981 | . | ! | ATTTTTT | 24.21 | -0.118 | 9.6E-15 | 0.266 | | *LCT* |
| chr2:135872341 | rs309128 | G | A | 24.16 | -0.117 | 1.1E-14 | 0.289 | | |
| chr2:135876201 | rs191079 | C | T | 24.17 | -0.118 | 1.1E-14 | 0.311 | Yes | |
| chr2:135868032 | rs309811 | ! | G | 24.15 | -0.117 | 1.1E-14 | 0.334 | | |
| chr2:135878754 | rs55830620 | G | A | 24.17 | -0.117 | 1.2E-14 | 0.459 | | |
| chr2:135873461 | rs309130 | C | A | 24.16 | -0.117 | 1.2E-14 | 0.480 | | |
| chr2:135873419 | rs188680 | A | G | 24.16 | -0.117 | 1.2E-14 | 0.501 | | |
| chr2:135873187 | rs680428 | A | G | 24.14 | -0.117 | 1.2E-14 | 0.521 | | |
| chr2:135880646 | rs632632 | C | T | 24.17 | -0.117 | 1.3E-14 | 0.637 | | *LCT* |
| chr2:135882663 | rs666407 | G | A | 24.17 | -0.117 | 1.3E-14 | 0.656 | | *LCT* |
| chr2:135885985 | rs309125 | T | C | 24.13 | -0.117 | 1.3E-14 | 0.675 | | *LCT* |
| chr2:135853028 | rs4988226 | A | G | 24.19 | -0.117 | 1.3E-14 | 0.695 | | *MCM6* |
| chr2:135853733 | rs57541371 | ! | GA | 24.25 | -0.117 | 1.4E-14 | 0.713 | | *MCM6* |
| chr2:135884312 | rs1435576 | T | A | 24.14 | -0.117 | 1.4E-14 | 0.731 | | *LCT* |
| chr2:135854054 | rs309178 | C | T | 24.19 | -0.117 | 1.4E-14 | 0.748 | | *RAB3GAP1* |
| chr2:135853733 | rs55820219 | GAAA | ! | 24.16 | -0.117 | 1.5E-14 | 0.765 | | *MCM6* |
| chr2:135927658 | rs309160 | T | C | 24.24 | -0.117 | 1.5E-14 | 0.782 | | *LCT* |
| chr2:135964425 | rs13404551 | C | T | 24.22 | -0.117 | 1.7E-14 | 0.797 | | |
| chr2:135983330 | rs687670 | C | T | 24.25 | -0.117 | 1.7E-14 | 0.812 | | |
| chr2:135864646 | rs309176 | T | C | 24.17 | -0.116 | 1.7E-14 | 0.827 | | *LCT* |
| chr2:135924704 | rs192822 | T | A | 24.25 | -0.117 | 1.8E-14 | 0.841 | | *LCT* |
| chr2:135858887 | . | G | ! | 24.08 | -0.116 | 1.9E-14 | 0.854 | | *LCT* |
| chr2:135856210 | rs309179 | G | A | 24.19 | -0.116 | 1.9E-14 | 0.867 | | *RAB3GAP1* |
| chr2:135856685 | rs309180 | G | A | 24.18 | -0.116 | 2.0E-14 | 0.880 | | *RAB3GAP1* |
| chr2:135858887 | rs4988202 | ! | GT | 24.19 | -0.116 | 2.2E-14 | 0.892 | | *LCT* |

| | | | | | | | | | |
|---|---|---|---|---|---|---|---|---|---|
| chr2:135859954 | rs309173 | C | T | 24.19 | -0.116 | 2.2E-14 | 0.903 | | *LCT* |
| chr2:135857243 | rs309181 | C | G | 24.19 | -0.116 | 2.3E-14 | 0.914 | | *RAB3GAP1* |
| chr2:135938568 | rs309145 | G | A | 24.25 | -0.116 | 2.5E-14 | 0.924 | | |
| chr2:135900775 | rs218174 | G | A | 24.30 | -0.116 | 2.5E-14 | 0.934 | | *LCT* |
| chr2:135912728 | rs309168 | T | ! | 24.26 | -0.116 | 2.8E-14 | 0.944 | | |
| chr2:135999276 | . | ! | CTTTTTTT | 24.60 | -0.115 | 2.8E-14 | 0.953 | | *CXCR4* |
| chr2:135912728 | . | ! | C | 24.26 | -0.116 | 3.2E-14 | 0.961 | | |
| chr2:135932847 | rs34744196 | AAAAAAAAAAAAG | ! | 24.05 | -0.114 | 7.2E-14 | 0.964 | | |
| chr2:135932847 | . | ! | AAAAAAAAAAAAAG | 24.19 | -0.114 | 7.9E-14 | 0.968 | | |
| chr2:135845706 | rs3754686 | T | C | 23.71 | -0.114 | 9.8E-14 | 0.970 | | |
| chr2:135845796 | rs3769005 | C | G | 23.70 | -0.114 | 9.8E-14 | 0.973 | | |
| chr2:135852405 | rs4954493 | T | C | 23.71 | -0.114 | 9.9E-14 | 0.976 | | *MCM6* |
| chr2:135850661 | rs4954490 | G | A | 23.71 | -0.113 | 1.0E-13 | 0.978 | | *MCM6* |
| chr2:135948872 | rs35323281 | GTCTCCAAAAAAAAAAAAA | ! | 22.38 | -0.115 | 1.6E-13 | 0.980 | Yes | |
| chr2:135978966 | . | ! | TTTCAATTTTTTTTTTTTTTT | 23.18 | -0.114 | 1.7E-13 | 0.982 | | |
| chr2:135894881 | . | ! | GTTTTTTTTTTTT | 22.79 | -0.114 | 2.0E-13 | 0.983 | | *LCT* |
| chr2:135860235 | rs160329 | C | T | 25.71 | -0.111 | 2.0E-13 | 0.984 | | *LCT* |
| chr2:135964032 | rs12615624 | TGATT | ! | 22.65 | -0.113 | 3.5E-13 | 0.985 | | |
| chr2:135964032 | . | ! | TAATT | 22.65 | -0.113 | 3.5E-13 | 0.986 | | |
| chr2:135978966 | rs5834455 | TTTCAATTTTTTTTTTTTTT | ! | 22.18 | -0.114 | 3.8E-13 | 0.987 | | |
| chr2:136018328 | rs2322818 | C | T | 24.31 | -0.112 | 4.2E-13 | 0.987 | | |
| chr2:135948872 | . | ! | GTCTCCAAAAAAAAAAAA | 22.67 | -0.112 | 5.3E-13 | 0.988 | Yes | |
| chr2:135940528 | rs309149 | T | C | 22.66 | -0.112 | 6.4E-13 | 0.988 | | *LCT* |
| chr2:135886700 | . | ! | ATGCCTC | 22.74 | -0.111 | 7.5E-13 | 0.989 | | *LCT* |
| chr2:135934255 | rs309164 | C | T | 22.67 | -0.111 | 7.6E-13 | 0.989 | | |
| **2q22, *CXCR4* rs555647251** | | | | | | | | | |
| chr2:136136760 | rs555647251 | T | C | 0.08 | 1.740 | 1.2E-11 | 0.242 | Yes | |
| chr2:136119459 | . | ! | G | 0.30 | 0.521 | 4.2E-07 | 0.831 | Yes | |
| chr2:136703620 | rs140018110 | C | A | 2.72 | -0.174 | 1.6E-05 | 0.910 | | *THSD7B* |
| chr2:136755749 | rs183299226 | G | A | 2.69 | -0.170 | 2.3E-05 | 0.966 | | |
| chr2:136119459 | rs770321415 | GCCCCCACCCCGAAGG | ! | 0.11 | 0.617 | 8.7E-05 | 0.988 | Yes | In *CXCR4* promoter |
| chr2:136102666 | rs202081231 | GTTTTTTTTTTT | ! | 0.09 | 0.812 | 1.6E-04 | 0.989 | | *LCT* |
| chr2:136102666 | . | ! | GTTTTTTTTTT | 0.09 | 0.811 | 1.6E-04 | 0.990 | | *LCT* |

| | | | | | | | | | |
|---|---|---|---|---|---|---|---|---|---|
| **3p22, *ITGA9*** | | | | | | | | | |
| chr3:37477922 | . | ! | TTT | 12.95 | -0.123 | 4.7E-11 | 0.068 | Yes | *ITGA9* |
| chr3:37477419 | rs11717950 | C | A | 12.68 | -0.121 | 9.7E-11 | 0.101 | | *ITGA9* |
| chr3:37476228 | rs73053283 | C | T | 12.67 | -0.121 | 1.0E-10 | 0.134 | | *ITGA9* |
| chr3:37477707 | rs17227369 | T | C | 12.68 | -0.121 | 1.0E-10 | 0.166 | Yes | *ITGA9* |
| chr3:37480987 | rs78502025 | T | C | 12.67 | -0.121 | 1.0E-10 | 0.199 | | *ITGA9* |
| chr3:37481301 | rs17813754 | G | T | 12.68 | -0.121 | 1.1E-10 | 0.229 | | *ITGA9* |
| chr3:37481340 | rs17036403 | C | A | 12.68 | -0.121 | 1.1E-10 | 0.258 | | *ITGA9* |
| chr3:37480319 | rs80091028 | C | T | 12.68 | -0.121 | 1.1E-10 | 0.288 | | *ITGA9* |
| chr3:37477922 | rs201494641 | T | ! | 12.68 | -0.121 | 1.1E-10 | 0.318 | Yes | *ITGA9* |
| chr3:37479134 | rs17227482 | G | A | 12.64 | -0.121 | 1.2E-10 | 0.345 | | *ITGA9* |
| chr3:37477598 | rs73053290 | T | C | 12.65 | -0.121 | 1.3E-10 | 0.370 | | *ITGA9* |
| chr3:37480497 | rs17227650 | T | C | 12.65 | -0.121 | 1.3E-10 | 0.396 | | *ITGA9* |
| chr3:37481384 | rs73055204 | T | C | 12.65 | -0.121 | 1.3E-10 | 0.421 | | *ITGA9* |
| chr3:37480326 | rs76945046 | T | C | 12.65 | -0.121 | 1.3E-10 | 0.446 | | *ITGA9* |
| chr3:37478380 | rs140918699 | GTGTTTC | ! | 12.65 | -0.120 | 1.3E-10 | 0.472 | | *ITGA9* |
| chr3:37479041 | rs17813574 | G | A | 12.65 | -0.120 | 1.3E-10 | 0.497 | | *ITGA9* |
| chr3:37479746 | rs73053300 | T | C | 12.65 | -0.120 | 1.4E-10 | 0.520 | | *ITGA9* |
| chr3:37479569 | rs73053299 | T | C | 12.64 | -0.120 | 1.4E-10 | 0.544 | | *ITGA9* |
| chr3:37479546 | rs75974193 | A | G | 12.65 | -0.120 | 1.4E-10 | 0.567 | | *ITGA9* |
| chr3:37479523 | rs73053296 | A | G | 12.64 | -0.120 | 1.4E-10 | 0.591 | | *ITGA9* |
| chr3:37479524 | rs73053297 | T | A | 12.65 | -0.120 | 1.4E-10 | 0.614 | | *ITGA9* |
| chr3:37478072 | rs17227404 | T | C | 12.65 | -0.120 | 1.4E-10 | 0.638 | Yes | *ITGA9* |
| chr3:37478380 | . | ! | GTGTTGTTTC | 12.65 | -0.120 | 1.4E-10 | 0.661 | | *ITGA9* |
| chr3:37476574 | rs61083478 | G | A | 12.72 | -0.120 | 1.4E-10 | 0.685 | | *ITGA9* |
| chr3:37476573 | . | ! | C | 12.71 | -0.120 | 1.4E-10 | 0.708 | | *ITGA9* |
| chr3:37476573 | rs57496508 | T | ! | 12.71 | -0.120 | 1.4E-10 | 0.732 | | *ITGA9* |
| chr3:37475576 | rs17227306 | T | C | 12.66 | -0.120 | 1.5E-10 | 0.754 | | *ITGA9* |
| chr3:37482970 | rs11710782 | T | C | 12.65 | -0.120 | 1.7E-10 | 0.773 | | *ITGA9* |
| chr3:37481588 | rs17227748 | A | G | 12.82 | -0.118 | 2.1E-10 | 0.789 | | *ITGA9* |
| chr3:37484928 | rs57409788 | A | G | 12.67 | -0.119 | 2.4E-10 | 0.803 | | *ITGA9* |
| chr3:37484943 | rs58527780 | A | G | 12.67 | -0.119 | 2.4E-10 | 0.817 | | *ITGA9* |
| chr3:37484582 | rs61636317 | T | C | 12.67 | -0.119 | 2.5E-10 | 0.831 | | *ITGA9* |

| | | | | | | | | | |
|---|---|---|---|---|---|---|---|---|---|
| chr3:37485015 | rs17814006 | C | ! | 12.68 | -0.118 | 2.6E-10 | 0.844 | | *ITGA9* |
| chr3:37485015 | . | ! | G | 12.68 | -0.118 | 2.6E-10 | 0.856 | | *ITGA9* |
| chr3:37485962 | rs17814180 | G | A | 12.67 | -0.118 | 2.7E-10 | 0.869 | | *ITGA9* |
| chr3:37485912 | rs17228097 | AA | ! | 12.68 | -0.118 | 2.7E-10 | 0.881 | | *ITGA9* |
| chr3:37486164 | rs73055214 | G | A | 12.68 | -0.118 | 2.8E-10 | 0.893 | | *ITGA9* |
| chr3:37484270 | rs10510691 | C | T | 12.68 | -0.118 | 2.8E-10 | 0.905 | | *ITGA9* |
| chr3:37484075 | rs10510690 | A | T | 12.67 | -0.118 | 2.8E-10 | 0.917 | | *ITGA9* |
| chr3:37485291 | rs17227972 | T | C | 12.67 | -0.118 | 2.9E-10 | 0.929 | | *ITGA9* |
| chr3:37483835 | . | ! | GAGTG | 12.67 | -0.118 | 3.5E-10 | 0.939 | | *ITGA9* |
| chr3:37483835 | rs144852530 | AGAGTG | ! | 12.67 | -0.118 | 3.5E-10 | 0.948 | | *ITGA9* |
| chr3:37487504 | rs11715360 | A | C | 12.68 | -0.118 | 3.9E-10 | 0.957 | | *ITGA9* |
| chr3:37488358 | rs11716496 | A | G | 12.69 | -0.118 | 4.1E-10 | 0.966 | | *ITGA9* |
| chr3:37488607 | rs112199844 | G | A | 12.68 | -0.118 | 4.6E-10 | 0.973 | | *ITGA9* |
| chr3:37489195 | rs11709684 | G | A | 12.76 | -0.117 | 5.7E-10 | 0.979 | | *ITGA9* |
| chr3:37485912 | . | ! | AT | 12.83 | -0.115 | 6.1E-10 | 0.985 | | *ITGA9* |
| chr3:37489253 | rs11709687 | G | A | 12.76 | -0.117 | 6.3E-10 | 0.990 | | *ITGA9* |
| **12q14, *PPM1H*** | | | | | | | | | |
| chr12:62912352 | rs699585 | G | T | 46.07 | -0.126 | 2.2E-23 | 0.062 | | *PPM1H* |
| chr12:62910977 | rs772559 | G | A | 46.09 | -0.126 | 2.3E-23 | 0.121 | Weak | *PPM1H* |
| chr12:62913073 | rs772565 | C | T | 46.06 | -0.126 | 3.0E-23 | 0.167 | | *PPM1H* |
| chr12:62913467 | . | ! | CT | 46.04 | -0.126 | 3.2E-23 | 0.210 | | |
| chr12:62913467 | rs772566 | CC | ! | 46.04 | -0.126 | 3.2E-23 | 0.252 | | |
| chr12:62911088 | rs2246214 | G | A | 45.88 | -0.126 | 3.3E-23 | 0.294 | | *PPM1H* |
| chr12:62910804 | rs772557 | G | A | 45.87 | -0.126 | 3.3E-23 | 0.335 | Yes | *PPM1H* |
| chr12:62913546 | rs772567 | C | T | 46.04 | -0.126 | 3.4E-23 | 0.376 | | |
| chr12:62911027 | rs772560 | G | A | 45.87 | -0.126 | 3.5E-23 | 0.415 | | *PPM1H* |
| chr12:62911765 | rs772563 | T | C | 45.86 | -0.126 | 3.6E-23 | 0.453 | | *PPM1H* |
| chr12:62913566 | rs772568 | G | C | 46.03 | -0.126 | 3.7E-23 | 0.490 | | |
| chr12:62911183 | rs772561 | A | C | 45.87 | -0.126 | 4.0E-23 | 0.525 | | *PPM1H* |
| chr12:62911372 | rs1666872 | G | A | 45.87 | -0.126 | 4.1E-23 | 0.558 | | *PPM1H* |
| chr12:62908362 | rs798403 | G | A | 46.38 | -0.126 | 4.3E-23 | 0.590 | | |
| chr12:62910667 | rs772556 | T | C | 45.87 | -0.126 | 4.4E-23 | 0.621 | Yes | *PPM1H* |
| chr12:62908764 | rs771986 | A | G | 46.38 | -0.126 | 4.6E-23 | 0.651 | | |

| | | | | | | | | | |
|---|---|---|---|---|---|---|---|---|---|
| chr12:62908028 | rs699584 | T | C | 46.39 | -0.126 | 4.6E-23 | 0.681 | | |
| chr12:62908956 | rs812224 | C | T | 46.38 | -0.126 | 4.8E-23 | 0.710 | | |
| chr12:62910503 | rs772555 | A | G | 45.87 | -0.125 | 4.8E-23 | 0.739 | Weak | *PPM1H* |
| chr12:62909042 | rs771985 | C | G | 46.40 | -0.125 | 5.0E-23 | 0.766 | | |
| chr12:62909404 | rs772550 | G | A | 46.38 | -0.125 | 5.1E-23 | 0.793 | | *PPM1H* |
| chr12:62909767 | rs2449865 | T | C | 46.38 | -0.125 | 5.2E-23 | 0.820 | | *PPM1H* |
| chr12:62909987 | rs812814 | C | G | 46.38 | -0.125 | 5.4E-23 | 0.845 | | *PPM1H* |
| chr12:62909852 | rs2449866 | T | C | 46.38 | -0.125 | 5.5E-23 | 0.870 | | *PPM1H* |
| chr12:62907614 | . | ! | TTGT | 46.39 | -0.125 | 6.1E-23 | 0.893 | | |
| chr12:62907614 | rs699583 | CTGT | ! | 46.39 | -0.125 | 6.1E-23 | 0.916 | | |
| chr12:62907518 | rs771988 | A | C | 46.39 | -0.125 | 6.3E-23 | 0.938 | | |
| chr12:62906716 | . | ! | CAACAATAGAAGAAGATAT | 46.48 | -0.125 | 1.1E-22 | 0.950 | | |
| chr12:62906716 | rs699580 | CAACAAAAGAAGAAGATAT | ! | 46.47 | -0.124 | 1.1E-22 | 0.963 | | |
| chr12:62907177 | rs771990 | C | T | 46.36 | -0.124 | 1.3E-22 | 0.974 | | |
| chr12:62906606 | rs1666900 | C | A | 46.52 | -0.124 | 1.4E-22 | 0.984 | | |
| chr12:62907076 | rs699582 | A | G | 46.48 | -0.124 | 1.5E-22 | 0.993 | | |
| **19p13, *ARHGAP45*** | | | | | | | | | |
| chr19:1078304 | rs35532684 | C | T | 9.53 | 0.143 | 2.6E-08 | 0.258 | | *SBNO2;C19orf26* |
| chr19:1079960 | rs36084354 | A | G | 8.81 | 0.148 | 2.7E-08 | 0.511 | | *SBNO2;C19orf26* |
| chr19:1078298 | rs35140707 | T | C | 9.50 | 0.143 | 2.7E-08 | 0.761 | | *SBNO2;C19orf26* |
| chr19:1074303 | rs12974537 | GGCCTTGTCCCAGTACCTCACAC | ! | 9.04 | 0.142 | 5.3E-08 | 0.893 | | |
| chr19:1074303 | . | ! | GGCCTTGTCCCAGCACCTCACAC | 9.11 | 0.141 | 6.7E-08 | 0.998 | | |
| **19p13, *CEBPA*** | | | | | | | | | |
| chr19:33264458 | rs12975577 | C | T | 48.96 | -0.082 | 1.2E-10 | 0.829 | | |
| chr19:33262414 | rs12150887 | G | C | 19.35 | -0.097 | 1.3E-09 | 0.928 | | |
| chr19:33263071 | rs112557032 | ! | TAAAAAAATTATT | 32.40 | -0.083 | 1.7E-09 | 0.995 | | |
| **5p15, *TERT*** | | | | | | | | | |
| chr5:1285859 | rs7705526 | A | C | 33.98 | 0.081 | 1.5E-09 | 0.468 | | |
| chr5:1282204 | rs7726159 | A | C | 34.22 | 0.080 | 1.8E-09 | 0.858 | | |
| chr5:1280823 | rs386684408 | GTGAGTCTCC | ! | 39.98 | 0.074 | 8.8E-09 | 0.939 | | |
| chr5:1282299 | rs7725218 | A | G | 35.59 | 0.075 | 1.6E-08 | 0.986 | | |
| chr5:1279849 | rs10054203 | C | G | 42.18 | 0.070 | 5.4E-08 | 1.000 | | |

| | | | | | | | | | |
|---|---|---|---|---|---|---|---|---|---|
| **8q24, *CCDC26* (long-distance interaction with *MYC*)** | | | | | | | | | |
| chr8:129611859 | rs1991866 | G | C | 44.86 | -0.075 | 3.5E-09 | 0.144 | | |
| chr8:129599498 | . | ACTCTCTTTT | ! | 36.93 | -0.075 | 1.1E-08 | 0.192 | | |
| chr8:129596822 | rs28500514 | T | C | 45.77 | -0.072 | 1.1E-08 | 0.240 | | |
| chr8:129597122 | rs4407843 | A | C | 45.80 | -0.072 | 1.1E-08 | 0.287 | | |
| chr8:129598143 | rs2163952 | C | T | 45.79 | -0.072 | 1.1E-08 | 0.334 | | |
| chr8:129594230 | rs1865223 | G | T | 45.83 | -0.072 | 1.1E-08 | 0.381 | | |
| chr8:129593625 | rs55964818 | T | C | 45.77 | -0.072 | 1.2E-08 | 0.424 | | |
| chr8:129593685 | rs12544501 | T | C | 45.79 | -0.072 | 1.2E-08 | 0.468 | | |
| chr8:129597074 | rs4480083 | C | T | 45.81 | -0.072 | 1.2E-08 | 0.511 | | |
| chr8:129602516 | rs1579555 | A | G | 45.51 | -0.072 | 1.6E-08 | 0.544 | | |
| chr8:129591389 | rs10107630 | C | T | 45.41 | -0.071 | 1.6E-08 | 0.577 | | *MYC* |
| chr8:129603606 | rs6989110 | T | C | 45.52 | -0.071 | 1.7E-08 | 0.608 | | |
| chr8:129592317 | rs13277237 | G | A | 45.87 | -0.071 | 1.8E-08 | 0.637 | Yes | *MYC* |
| chr8:129601253 | rs13280084 | T | G | 45.52 | -0.071 | 1.9E-08 | 0.665 | | |
| chr8:129603303 | rs4291235 | T | C | 45.49 | -0.071 | 2.0E-08 | 0.691 | | |
| chr8:129600947 | rs28854585 | A | G | 45.48 | -0.071 | 2.1E-08 | 0.717 | | |
| chr8:129609008 | rs35389394 | ! | T | 45.54 | -0.071 | 2.2E-08 | 0.741 | | |
| chr8:129609008 | . | C | ! | 45.54 | -0.071 | 2.2E-08 | 0.765 | | |
| chr8:129602762 | rs2217715 | C | A | 45.59 | -0.070 | 2.6E-08 | 0.785 | | |
| chr8:129598493 | rs2395905 | T | C | 45.57 | -0.070 | 2.7E-08 | 0.805 | | |
| chr8:129600857 | rs4486162 | A | G | 45.53 | -0.070 | 2.7E-08 | 0.825 | | |
| chr8:129601368 | rs10098310 | G | A | 45.58 | -0.070 | 2.8E-08 | 0.844 | | |
| chr8:129612415 | rs10092988 | A | G | 49.98 | 0.070 | 2.9E-08 | 0.862 | | |
| chr8:129605834 | rs62525616 | C | T | 45.60 | -0.070 | 2.9E-08 | 0.881 | | |
| chr8:129607873 | rs13276769 | G | A | 45.61 | -0.070 | 3.0E-08 | 0.899 | | |
| chr8:129598538 | rs2395906 | ! | CTCTTACACA | 45.60 | -0.070 | 3.0E-08 | 0.917 | | |
| chr8:129598538 | . | CTCTTACACG | ! | 45.59 | -0.070 | 3.0E-08 | 0.935 | | |
| chr8:129599841 | rs2395902 | T | G | 45.52 | -0.070 | 3.2E-08 | 0.951 | | |
| chr8:129605779 | rs62525615 | C | T | 45.60 | -0.070 | 3.2E-08 | 0.968 | | |
| chr8:129606265 | rs7341546 | A | T | 45.58 | -0.070 | 3.4E-08 | 0.984 | | |
| chr8:129590035 | rs10103048 | A | C | 43.39 | -0.068 | 8.8E-08 | 0.990 | | *MYC* |

### Supplementary Table 5

*cis*-eQTLs for the identified variants in $CD34^+$ cells, as detected in mRNA-sequencing data for $CD34^+$ isolated from 155 blood donors using fluorescence-activated cell sorting (**Online Methods**). In the analysis, we considered all genes in the credible set regions of all variants that were polymorphic in the mRNA-sequencing data set (*i.e.*, not the region of rs55564721), extended by 0.1 Mb. These regions harbor 26 genes, and 4 tests were done four *CXCR4*, yielding a Bonferroni limit of 0.05/29 = 1.7E-3. The table shows Bonferroni-significant associations, and the directed testing results for *CXCR4*, which were motivated by analysis of the PCHi-C/ATAC-seq data and known biology of *CXCR4*. Statistics are for linear modeling with 10 expression principal component covariates (calculated using all genes with median FPKM > 1.0). For the *CXCR4 locus*, we also included the other common variants at the locus as covariates.

| **Locus** | **$CD34^+$ level marker** | **Position (hg38)** | **eQTL marker** | **Position (hg38)** | **$CD34^+$/eQTL marker correlation ($r^2$)** | **Ref allele** | **Alt allele** | **eQTL gene** | ***P*-value** | **Effect (z)** |
|---|---|---|---|---|---|---|---|---|---|---|
| 7p15.3 | rs699585 | 62912352 | rs772559 | 62912352 | 0.99 | A | G | *PPM1H* | 6.79E-51 | 23.50 |
| 1p36 | rs2047094 | 8857756 | rs11121246 | 8857043 | 1.00 | T | G | *ENO1* | 3.10E-42 | -19.40 |
| 1p36 | rs2047094 | 8857756 | rs11121246 | 8857043 | 1.00 | T | G | *RERE* | 3.47E-08 | -5.68 |
| 3p22 | rs201494641 | 37477913 | rs17227369 | 37477707 | 1.00 | C | T | *ITGA9* | 2.02E-11 | -7.16 |
| 2p22 | rs10193623 | 136143991 | rs6726457 | 136142899 | 0.99 | C | T | *CXCR4* | 5.40E-03 | 2.32 |
| 2p22 | rs11688530 | 136125862 | rs11688530 | 136125862 | 1.00 | G | A | *CXCR4* | 3.70E-02 | -1.45 |
| 2p22 | rs309137 | 136008381 | rs309137 | 136008381 | 1.00 | T | C | *CXCR4* | 1.77E-02 | 1.82 |

### Supplementary Table 6

Associations with human diseases and traits (r2 > 0.8 with a CD34+ lead variant) identified by UK BioBank PhenomeScanner.

| $CD34^+$ rsID | hg38 position | trait rsID | hg38 position | a1 | a2 | $r^2$ | trait | *P* | β | dataset |
|---|---|---|---|---|---|---|---|---|---|---|
| rs11688530 | chr2:136125862 | rs11688530 | chr2:136125862 | A | G | 1.00 | Granulocyte percentage of myeloid white cells | 3.20E-11 | 0.05 | A |
| rs11688530 | chr2:136125862 | rs11688530 | chr2:136125862 | A | G | 1.00 | Monocyte count | 4.34E-09 | -0.04 | A |
| rs11688530 | chr2:136125862 | rs11688530 | chr2:136125862 | A | G | 1.00 | Monocyte percentage of white cells | 4.49E-13 | -0.05 | A |
| rs11688530 | chr2:136125862 | rs77880275 | chr2:136103697 | C | A | 0.94 | Platelet count | 1.29E-06 | 0.04 | A |
| rs11688530 | chr2:136125862 | rs77880275 | chr2:136103697 | C | A | 0.94 | Plateletcrit | 1.17E-07 | 0.04 | A |
| rs12975577 | chr19:33264458 | rs10401672 | chr19:33269055 | C | T | 0.61 | Basophil count | 1.88E-42 | 0.05 | A |
| rs12975577 | chr19:33264458 | rs10401672 | chr19:33269055 | C | T | 0.61 | Basophil percentage of granulocytes | 1.00E-39 | -0.05 | C |
| rs12975577 | chr19:33264458 | rs10401672 | chr19:33269055 | C | T | 0.61 | Basophil percentage of white cells | 6.56E-48 | 0.05 | A |
| rs12975577 | chr19:33264458 | rs12975577 | chr19:33264458 | C | T | 1.00 | Crohns disease | 7.00E-06 | NA | D |
| rs12975577 | chr19:33264458 | rs17755899 | chr19:33264735 | T | C | 0.64 | High light scatter percentage of red cells | 2.32E-06 | 0.02 | A |
| rs12975577 | chr19:33264458 | rs12975577 | chr19:33264458 | C | T | 1.00 | Immature fraction of reticulocytes | 1.78E-07 | 0.02 | A |
| rs12975577 | chr19:33264458 | rs17755899 | chr19:33264735 | T | C | 0.64 | Inflammatory bowel disease | 5.32E-06 | 0.09 | E |
| rs12975577 | chr19:33264458 | rs12975577 | chr19:33264458 | C | T | 1.00 | Mean corpuscular hemoglobin | 3.16E-30 | 0.04 | A |
| rs12975577 | chr19:33264458 | rs12975577 | chr19:33264458 | C | T | 1.00 | Mean corpuscular volume | 3.71E-32 | 0.04 | A |
| rs12975577 | chr19:33264458 | rs17755899 | chr19:33264735 | T | C | 0.64 | Monocyte percentage of white cells | 3.30E-07 | 0.02 | A |
| rs12975577 | chr19:33264458 | rs12975577 | chr19:33264458 | C | T | 1.00 | Red blood cell count | 2.82E-19 | -0.03 | A |
| rs12975577 | chr19:33264458 | rs10401672 | chr19:33269055 | C | T | 0.61 | White blood cell count basophil | 2.00E-42 | -0.05 | C |
| rs1991866 | chr8:129611859 | rs4291235 | chr8:129603303 | T | C | 0.95 | Allergic disease | 8.99E-06 | -0.03 | F |
| rs1991866 | chr8:129611859 | rs201913843 | chr8:129605779 | C | T | 0.95 | Basophil percentage of granulocytes | 3.26E-09 | -0.02 | A |
| rs1991866 | chr8:129611859 | rs201913843 | chr8:129605779 | C | T | 0.95 | Basophil percentage of white cells | 4.91E-08 | -0.02 | A |
| rs1991866 | chr8:129611859 | rs4291235 | chr8:129603303 | T | C | 0.95 | Eosinophil count | 4.63E-07 | -0.02 | A |
| rs1991866 | chr8:129611859 | rs4291235 | chr8:129603303 | T | C | 0.95 | Eosinophil percentage of granulocytes | 1.60E-19 | -0.03 | A |
| rs1991866 | chr8:129611859 | rs4291235 | chr8:129603303 | T | C | 0.95 | Eosinophil percentage of white cells | 3.14E-18 | -0.03 | A |
| rs1991866 | chr8:129611859 | rs1991866 | chr8:129611859 | G | C | 1.00 | Granulocyte count | 3.02E-24 | 0.04 | A |
| rs1991866 | chr8:129611859 | rs201913843 | chr8:129605779 | C | T | 0.95 | Granulocyte percentage of myeloid white cells | 1.05E-62 | -0.06 | A |
| rs1991866 | chr8:129611859 | rs4291235 | chr8:129603303 | T | C | 0.95 | Immature fraction of reticulocytes | 2.12E-07 | 0.02 | A |
| rs1991866 | chr8:129611859 | rs1991866 | chr8:129611859 | G | C | 1.00 | Inflammatory bowel disease | 1.65E-09 | NA | G |
| rs1991866 | chr8:129611859 | rs2395902 | chr8:129599841 | T | G | 0.95 | Lymphocyte percentage of white cells | 5.98E-30 | -0.04 | A |
| rs1991866 | chr8:129611859 | rs1991866 | chr8:129611859 | G | C | 1.00 | Mean corpuscular hemoglobin | 2.72E-12 | 0.03 | A |

| | | | | | | | | | | |
|---|---|---|---|---|---|---|---|---|---|---|
| rs1991866 | chr8:129611859 | rs1991866 | chr8:129611859 | G | C | 1.00 | Mean corpuscular volume | 6.72E-17 | 0.03 | A |
| rs1991866 | chr8:129611859 | rs201913843 | chr8:129605779 | C | T | 0.95 | Monocyte count | 7.53E-177 | 0.10 | A |
| rs1991866 | chr8:129611859 | rs1991866 | chr8:129611859 | G | C | 1.00 | Monocyte lymphocyte ratio | 2.00E-06 | NA | C |
| rs1991866 | chr8:129611859 | rs201913843 | chr8:129605779 | C | T | 0.95 | Monocyte percentage of white cells | 3.31E-117 | 0.08 | A |
| rs1991866 | chr8:129611859 | rs1991866 | chr8:129611859 | G | C | 1.00 | Myeloid white cell count | 7.82E-38 | 0.05 | A |
| rs1991866 | chr8:129611859 | rs1991866 | chr8:129611859 | G | C | 1.00 | Neutrophil count | 7.32E-27 | 0.04 | A |
| rs1991866 | chr8:129611859 | rs4291235 | chr8:129603303 | T | C | 0.95 | Neutrophil percentage of granulocytes | 9.69E-22 | 0.03 | A |
| rs1991866 | chr8:129611859 | rs4291235 | chr8:129603303 | T | C | 0.95 | Neutrophil percentage of white cells | 3.47E-10 | 0.02 | A |
| rs1991866 | chr8:129611859 | rs4291235 | chr8:129603303 | T | C | 0.95 | Red blood cell count | 6.34E-09 | -0.02 | A |
| rs1991866 | chr8:129611859 | rs1991866 | chr8:129611859 | G | C | 1.00 | Sum basophil neutrophil counts | 1.94E-26 | 0.04 | A |
| rs1991866 | chr8:129611859 | rs1991866 | chr8:129611859 | G | C | 1.00 | Sum neutrophil eosinophil counts | 1.46E-24 | 0.04 | A |
| rs1991866 | chr8:129611859 | rs1991866 | chr8:129611859 | G | C | 1.00 | Ulcerative colitis | 2.00E-06 | NA | C |
| rs1991866 | chr8:129611859 | rs1991866 | chr8:129611859 | G | C | 1.00 | White blood cell count | 6.78E-26 | 0.04 | A |
| rs1991866 | chr8:129611859 | rs1991866 | chr8:129611859 | G | C | 1.00 | White blood cell count monocyte | 8.00E-10 | -0.03 | C |
| rs2047094 | chr1:8857756 | rs79565040 | chr1:8857339 | TTA | T | 1.00 | Basophil count | 1.20E-09 | 0.02 | A |
| rs2047094 | chr1:8857756 | rs11121246 | chr1:8857043 | G | T | 1.00 | Granulocyte count | 4.05E-11 | 0.02 | A |
| rs2047094 | chr1:8857756 | rs2047094 | chr1:8857756 | A | C | 1.00 | Mean platelet volume | 1.97E-06 | 0.02 | A |
| rs2047094 | chr1:8857756 | rs11121246 | chr1:8857043 | G | T | 1.00 | Myeloid white cell count | 2.29E-10 | 0.02 | A |
| rs2047094 | chr1:8857756 | rs11121246 | chr1:8857043 | G | T | 1.00 | Neutrophil count | 1.14E-10 | 0.02 | A |
| rs2047094 | chr1:8857756 | rs11121246 | chr1:8857043 | G | T | 1.00 | Neutrophil percentage of white cells | 8.49E-06 | 0.02 | A |
| rs2047094 | chr1:8857756 | rs11121246 | chr1:8857043 | G | T | 1.00 | Platelet count | 1.85E-06 | 0.02 | A |
| rs2047094 | chr1:8857756 | rs11121246 | chr1:8857043 | G | T | 1.00 | Plateletcrit | 3.00E-16 | 0.03 | C |
| rs2047094 | chr1:8857756 | rs11121246 | chr1:8857043 | G | T | 1.00 | Sum basophil neutrophil counts | 6.28E-11 | 0.02 | A |
| rs2047094 | chr1:8857756 | rs11121246 | chr1:8857043 | G | T | 1.00 | Sum neutrophil eosinophil counts | 7.35E-11 | 0.02 | A |
| rs2047094 | chr1:8857756 | rs11121246 | chr1:8857043 | G | T | 1.00 | White blood cell count | 2.47E-09 | 0.02 | A |
| rs2047094 | chr1:8857756 | rs79565040 | chr1:8857339 | TTA | T | 1.00 | White blood cell count basophil | 1.00E-09 | NA | C |
| rs309137 | chr2:136008381 | rs309137 | chr2:136008381 | C | T | 1.00 | Arthritis including non Rheumatoid | 3.22E-06 | NA | G |
| rs309137 | chr2:136008381 | rs309137 | chr2:136008381 | C | T | 1.00 | Granulocyte percentage of myeloid white cells | 1.76E-06 | 0.02 | A |
| rs309137 | chr2:136008381 | rs309137 | chr2:136008381 | C | T | 1.00 | Monocyte percentage of white cells | 5.56E-07 | -0.02 | A |
| rs309137 | chr2:136008381 | rs309137 | chr2:136008381 | C | T | 1.00 | Neuroblastoma | 3.48E-07 | NA | H |
| rs309137 | chr2:136008381 | rs309168 | chr2:135912728 | T | C | 0.91 | Parkinsons disease | 2.36E-07 | -0.09 | I |
| rs309137 | chr2:136008381 | rs309137 | chr2:136008381 | C | T | 1.00 | Rheumatoid arthritis | 3.22E-06 | NA | G |
| rs36084354 | chr19:1079960 | rs36084354 | chr19:1079960 | G | A | 1.00 | Eosinophil count | 4.03E-12 | 0.04 | A |
| rs36084354 | chr19:1079960 | rs36084354 | chr19:1079960 | G | A | 1.00 | Eosinophil counts | 4.00E-12 | -0.04 | C |
| rs36084354 | chr19:1079960 | rs36084354 | chr19:1079960 | G | A | 1.00 | Eosinophil percentage of granulocytes | 1.00E-11 | -0.04 | C |

| | | | | | | | | | | |
|---|---|---|---|---|---|---|---|---|---|---|
| rs36084354 | chr19:1079960 | rs36084354 | chr19:1079960 | G | A | 1.00 | Eosinophil percentage of white cells | 9.05E-09 | 0.04 | A |
| rs36084354 | chr19:1079960 | rs36084354 | chr19:1079960 | G | A | 1.00 | Lymphocyte count | 2.66E-21 | 0.06 | A |
| rs36084354 | chr19:1079960 | rs36084354 | chr19:1079960 | G | A | 1.00 | Lymphocyte counts | 3.00E-21 | -0.06 | C |
| rs36084354 | chr19:1079960 | rs35532684 | chr19:1078304 | T | C | 0.84 | Lymphocyte percentage of white cells | 2.00E-11 | -0.04 | C |
| rs36084354 | chr19:1079960 | rs36084354 | chr19:1079960 | G | A | 1.00 | Mean corpuscular volume | 2.75E-08 | -0.03 | A |
| rs36084354 | chr19:1079960 | rs12974537 | chr19:1074316 | C | T | 0.95 | Mean platelet volume | 1.00E-12 | -0.04 | C |
| rs36084354 | chr19:1079960 | rs12974537 | chr19:1074316 | C | T | 0.95 | Monocyte count | 9.66E-06 | 0.03 | A |
| rs36084354 | chr19:1079960 | rs36084354 | chr19:1079960 | G | A | 1.00 | Neutrophil percentage of granulocytes | 7.90E-10 | -0.04 | A |
| rs36084354 | chr19:1079960 | rs36084354 | chr19:1079960 | G | A | 1.00 | Neutrophil percentage of white cells | 2.00E-12 | -0.04 | C |
| rs36084354 | chr19:1079960 | rs12974537 | chr19:1074316 | C | T | 0.95 | Red cell distribution width | 7.20E-06 | -0.03 | A |
| rs36084354 | chr19:1079960 | rs36084354 | chr19:1079960 | G | A | 1.00 | Sum eosinophil basophil counts | 7.00E-11 | -0.04 | C |
| rs36084354 | chr19:1079960 | rs62131205 | chr19:1084027 | G | C | 0.74 | White blood cell count | 3.27E-06 | 0.03 | A |
| rs699585 | chr12:62912352 | rs772567 | chr12:62913546 | T | C | 1.00 | Mean corpuscular volume | 4.55E-06 | -0.02 | A |
| rs7705526 | chr5:1285859 | rs7726159 | chr5:1282204 | A | C | 0.79 | Breast cancer | 3.00E-08 | NA | C |
| rs7705526 | chr5:1285859 | rs7726159 | chr5:1282204 | A | C | 0.79 | Breast cancer estrogen receptor negative | 2.00E-06 | NA | C |
| rs7705526 | chr5:1285859 | rs7705526 | chr5:1285859 | A | C | 1.00 | Chronic lymphocytic leukemia | 6.00E-10 | 0.17 | C |
| rs7705526 | chr5:1285859 | rs7705526 | chr5:1285859 | A | C | 1.00 | Eosinophil percentage of granulocytes | 4.03E-07 | -0.02 | A |
| rs7705526 | chr5:1285859 | rs7705526 | chr5:1285859 | A | C | 1.00 | Eosinophil percentage of white cells | 9.40E-06 | -0.02 | A |
| rs7705526 | chr5:1285859 | rs7705526 | chr5:1285859 | A | C | 1.00 | Epithelial ovarian cancer | 8.00E-08 | 0.07 | C |
| rs7705526 | chr5:1285859 | rs7705526 | chr5:1285859 | A | C | 1.00 | Granulocyte count | 9.80E-19 | 0.03 | A |
| rs7705526 | chr5:1285859 | rs7734992 | chr5:1280013 | C | T | 0.55 | Granulocyte percentage of myeloid white cells | 2.74E-07 | 0.02 | A |
| rs7705526 | chr5:1285859 | rs4449583 | chr5:1284020 | T | C | 0.78 | High grade serous ovarian cancer | 2.65E-12 | 0.12 | J |
| rs7705526 | chr5:1285859 | rs7705526 | chr5:1285859 | A | C | 1.00 | Invasive epithelial ovarian cancer | 7.00E-10 | 0.09 | C |
| rs7705526 | chr5:1285859 | rs4449583 | chr5:1284020 | T | C | 0.78 | Invasive ovarian cancer | 7.76E-12 | 0.10 | K |
| rs7705526 | chr5:1285859 | rs7705526 | chr5:1285859 | A | C | 1.00 | Low grade and borderline serous ovarian cancer | 1.58E-20 | 0.28 | K |
| rs7705526 | chr5:1285859 | rs7705526 | chr5:1285859 | A | C | 1.00 | Low grade serous and borderline ovarian cancer | 2.00E-20 | NA | C |
| rs7705526 | chr5:1285859 | rs7705526 | chr5:1285859 | A | C | 1.00 | Lung adenocarcinoma | 4.00E-35 | 0.22 | C |
| rs7705526 | chr5:1285859 | rs7705526 | chr5:1285859 | A | C | 1.00 | Lung cancer in never smokers | 7.00E-08 | 0.21 | C |
| rs7705526 | chr5:1285859 | rs7734992 | chr5:1280013 | C | T | 0.55 | Lymphocyte percentage of white cells | 2.84E-08 | -0.02 | A |
| rs7705526 | chr5:1285859 | rs7734992 | chr5:1280013 | C | T | 0.55 | Malignant neoplasm of brain | 2.92E-06 | 0.00 | L |
| rs7705526 | chr5:1285859 | rs4449583 | chr5:1284020 | T | C | 0.78 | Mean corpuscular hemoglobin | 5.65E-33 | -0.05 | A |
| rs7705526 | chr5:1285859 | rs4449583 | chr5:1284020 | T | C | 0.78 | Mean corpuscular volume | 1.00E-35 | 0.05 | C |
| rs7705526 | chr5:1285859 | rs7705526 | chr5:1285859 | A | C | 1.00 | Mean platelet volume | 3.81E-08 | 0.02 | A |
| rs7705526 | chr5:1285859 | rs7705526 | chr5:1285859 | A | C | 1.00 | Myeloid white cell count | 6.00E-19 | 0.03 | C |
| rs7705526 | chr5:1285859 | rs7705526 | chr5:1285859 | A | C | 1.00 | Neutrophil count | 7.91E-20 | 0.04 | A |

| | | | | | | | | | | |
|---|---|---|---|---|---|---|---|---|---|---|
| rs7705526 | chr5:1285859 | rs7705526 | chr5:1285859 | A | C | 1.00 | Neutrophil percentage of granulocytes | 1.33E-07 | 0.02 | A |
| rs7705526 | chr5:1285859 | rs7734992 | chr5:1280013 | C | T | 0.55 | Neutrophil percentage of white cells | 3.00E-11 | 0.02 | C |
| rs7705526 | chr5:1285859 | rs7705526 | chr5:1285859 | A | C | 1.00 | Platelet count | 2.18E-20 | 0.04 | A |
| rs7705526 | chr5:1285859 | rs7734992 | chr5:1280013 | C | T | 0.55 | Platelet distribution width | 1.75E-08 | 0.02 | A |
| rs7705526 | chr5:1285859 | rs7705526 | chr5:1285859 | A | C | 1.00 | Plateletcrit | 1.52E-41 | 0.05 | A |
| rs7705526 | chr5:1285859 | rs7725218 | chr5:1282299 | A | G | 0.71 | Prostate cancer | 3.00E-11 | NA | C |
| rs7705526 | chr5:1285859 | rs4449583 | chr5:1284020 | T | C | 0.78 | Red blood cell count | 1.80E-25 | 0.04 | A |
| rs7705526 | chr5:1285859 | rs4449583 | chr5:1284020 | T | C | 0.78 | Seborrhoeic keratosis | 9.80E-06 | 0.00 | L |
| rs7705526 | chr5:1285859 | rs7705526 | chr5:1285859 | A | C | 1.00 | Polycythaemia vera | 6.80E-06 | 0.00 | L |
| rs7705526 | chr5:1285859 | rs7705526 | chr5:1285859 | A | C | 1.00 | Serous borderline ovarian cancer | 5.50E-19 | 0.32 | K |
| rs7705526 | chr5:1285859 | rs4449583 | chr5:1284020 | T | C | 0.78 | Serous invasive ovarian cancer | 7.99E-14 | 0.12 | K |
| rs7705526 | chr5:1285859 | rs7705526 | chr5:1285859 | A | C | 1.00 | Sum basophil neutrophil counts | 1.00E-19 | 0.04 | C |
| rs7705526 | chr5:1285859 | rs7705526 | chr5:1285859 | A | C | 1.00 | Sum neutrophil eosinophil counts | 5.62E-19 | 0.03 | A |
| rs7705526 | chr5:1285859 | rs7705526 | chr5:1285859 | A | C | 1.00 | Telomere length | 6.00E-06 | 0.09 | C |
| rs7705526 | chr5:1285859 | rs7705526 | chr5:1285859 | A | C | 1.00 | White blood cell count | 7.74E-18 | 0.03 | A |

**PhenomeScanner data sets: (A)** Astle-W_Blood-Cell-Traits_EUR_2016; **(B)** van-der-Harst-P_CAD_Mixed_2018; **(C)** NHGRI-EBI_GWAS_Catalog; **(D)** IBDGC_Chrons-Disease_EUR_2012; **(E)** IBDGC_IBD_EUR_2015; **(F)** Ferreira-M_Allergic-Disease_EUR_2017; **(G)** GRASP; **(H)** dbGap; **(I)** Nalls-M_Parkinsons-Disease_EUR_2014; **(J)** Phelan-M_HGSOC_EUR_2017; **(K)** Phelan-M_Invasive-ovarian-cancer_EUR_2017; **(L)** Neale-B_UKBB_EUR_2017.

### Supplementary Table 7

sgRNA and primer sequences used in CRISPR/Cas9 experiments for the *CXCR4* and *PPM1H* loci.

| sgRNA(s) (+ strand) | | PAM | | Stran | | Primers for deletion confirmation | Deletion |
|---|---|---|---|---|---|---|---|
| 5' | 3' | 5' | 3' | 5' | 3' | Forward/reverse | size (bp) |
| sgRNA pair for deleting *CXCR4* rs309107 region (chr2:136,008,381; "V1") | | | | | | | |
| CACCG**ACAGTGGGGTGTGACAGTCA** | CACCG**CCAAGGAAAACTATGGGTGG** | CGG | GGG | + | + | ACCAGCAACCCTAATCTGCC | 587 |
| chr2:136,008,297-136,008,316 | chr2:136,008,884-136,008,903 | | | | | TGCCCTCCAGATGACTTTGA | |
| sgRNA pair for deleting *CXCR4* rs59222832 region (chr2:136,122,815; "V2") | | | | | | | |
| CACC**GGCGGGGTGTGGTTAGGCAA** | CACCG**TTCCCTCAAAGGGCAATGGA** | GGG | GGG | - | - | AGACTATTTCTCAAGGCGCAAG | 1170 |
| chr2:136,122,245-136,122,264 | chr2:136,123,415-136,123,434 | | | | | TCAGTGTAAACTGGAAGACTGGA | |
| sgRNA pair for deleting *CXCR4* rs770321415 region (chr2:136,119,459; "V3") | | | | | | | |
| CACCG**CTGTGATGGTAATACCCACA** | CACCG**AGAAACTCCAGGTTCTTGGA** | CGG | GGG | - | + | TCACTAGGGTCAGGTGCAGA | 878 |
| chr2:136,118,943-136,118,962 | chr2:136,119,807-136,119,826 | | | | | AGACATCGTGCAGGGAGGAG | |
| sgRNA pair for deleting *CXCR4* rs10193623 region (chr2:136,143,990; "V4") | | | | | | | |
| CACC**GAGCTTATCACATCAAATGG** | CACC**GAAGCACAGAGAGCTCAATG** | GGG | TGG | + | + | GGGAAACCTCAATAGAGACTAGAGG | 1025 |
| chr2:136,143,767-136,143,786 | chr2:136,144,792-136,144,811 | | | | | TTGTCCAGGTTTGTGCCATC | |
| sgRNA pair for deleting random control region (intronic region in *WAC*) | | AGG | AGG | - | - | GGAACTTTGACCTTTTAAAGATTTCAGTTTGAGATC | 139 |
| CACCG**CTCCTACCACAAAGAGCTTG** | CACCG**CAACATGTAGGAGAGAATAT** | | | | | GAACTACTTTCTTCTCATCCAAATTATTCAATCCCAG | |
| chr10:28,536,680-28,536,702 | chr10:28,536,819-28,536,841 | | | | | | |
| sgRNA *PPM1H* for disrupting rs772557 (chr12:62,910,704) | | | | | | | |
| CACC**GAAGAATCAGACCAAGCC[A/G]T** | | TGG | | + | | | n/a |
| chr12:62,910,786-62,910,805 | | | | | | | |

### Supplementary Table 8

Motif analysis for rs772557 using PERFECTOS-APE with the following motif databases: **(A)** HOCOMOCO-11, **(B)** JASPAR, **(C)** HT-SELEX, **(D)** SwissRegulon, and **(E)** HOMER. The analysis predicts changes in MYB binding with several motif models.

| rsID | DB | Motif | Alleles | Reference allele | | | | Alternative allele | | | | Foldchange |
|---|---|---|---|---|---|---|---|---|---|---|---|---|
| | | | | Pos | Strand | Word | P | Pos | Strand | Word | P | |
| rs772557 | A | CXXC1_HUMAN.H11MO.0.D | A/G | -1 | pos | cAttgga | 0.029002 | -1 | pos | cGttgga | 1.07E-04 | 2.72E+02 |
| rs772557 | C | MYBL2_3 | A/G | -6 | pos | caagccAttgg | 0.033675 | -6 | pos | caagccGttgg | 3.78E-04 | 8.90E+01 |
| rs772557 | C | MYBL1_3 | A/G | -6 | pos | caagccAttgg | 0.033675 | -6 | pos | caagccGttgg | 4.60E-04 | 7.32E+01 |
| rs772557 | A | MYBA_HUMAN.H11MO.0.D | A/G | -7 | pos | ccaagccAttggagc | 0.003097 | -7 | pos | ccaagccGttggagc | 5.34E-05 | 5.80E+01 |
| rs772557 | A | MYB_HUMAN.H11MO.0.A | A/G | -7 | pos | ccaagccAttgg | 0.013239 | -7 | pos | ccaagccGttgg | 3.50E-04 | 3.78E+01 |
| rs772557 | D | MYB.p2 | A/G | -4 | pos | agccAttg | 0.003081 | -4 | pos | agccGttg | 1.29E-04 | 2.38E+01 |
| rs772557 | E | MYB_ERMYB-Myb-ChIPSeq | A/G | -4 | pos | agccAttg | 0.006269 | -4 | pos | agccGttg | 3.59E-04 | 1.75E+01 |
| rs772557 | A | ZN134_HUMAN.H11MO.1.C | A/G | -9 | neg | gctccaaTggcttggtc | 2.89E-03 | -9 | neg | gctccaaCggcttggtc | 2.24E-04 | 1.29E+01 |
| rs772557 | A | MLXPL_HUMAN.H11MO.0.D | A/G | -15 | pos | gaatcagaccaagccAttg | 4.45E-04 | -15 | pos | gaatcagaccaagccGttg | 7.72E-05 | 5.76E+00 |
| rs772557 | A | MCR_HUMAN.H11MO.0.D | A/G | -11 | pos | cagaccaagccAttgga | 5.37E-04 | -11 | pos | cagaccaagccGttgga | 9.61E-05 | 5.592486371 |
| rs772557 | C | SOX1_1 | A/G | -9 | pos | gaccaagccAttgga | 4.8E-04 | -9 | pos | gaccaagccGttgga | 0.005023 | 9.6E-02 |

#### Supplementary Table 9

Genome coordinates of human Blood ENhancer Cluster (BENC) modules. Data from Bahr *et al.* , *Nature* 2018 Jan 25;553(7689):515-520. Converted from hg19 to hg38 coordinates using the UCSC LiftOver tool. The credible set belonging to the 8q24 association with blood CD34$^{+}$ levels spans hg38 chr8:129590035-129612415, corresponding to module D (red).

| | **Chr 8 position (hg19)** | | **Chr 8 position (hg38)** | |
|---|---|---|---|---|
| **Bahr *et al.* module** | **Start** | **End** | **Start** | **End** |
| A | 130558974 | 130559436 | 129546728 | 129547190 |
| B | 130564564 | 130566224 | 129552318 | 129553978 |
| C | 130594181 | 130594694 | 129581935 | 129582448 |
| D | 130604090 | 130605985 | 129591844 | 129593739 |
| E | 130627170 | 130627469 | 129614924 | 129615223 |
| F | 130648267 | 130648647 | 129636021 | 129636401 |
| G | 130678561 | 130679020 | 129666315 | 129666774 |
| H | 130705012 | 137705415 | 129692766 | 136693172 |

### Supplementary Table 10

Association statistics with blood $CD34^+$ levels for significant variants in blood donors *vs* primary care patients (Swedes and non-Swedes combined). All variants showed effects in the same direction in the two subgroups. Only the rare variant rs555647251 showed Bonferroni-significant heterogeneity (9 tests) between the two subgroups. The other 8 variants did not display significant heterogeneity.

| | | | | | | Blood donors | | Primary care patients | | |
|---|---|---|---|---|---|---|---|---|---|---|
| **Position** | **rsID** | **Gene** | **OA** | **EA** | **EAF (%)** | **β** | ***P*** | **β** | ***P*** | $P_{het}$ |
| chr1:8857756 | rs2047094 | *ENO1, RERE* | A | C | 49.2 | -0.118 | 1.10E-09 | -0.085 | 6.20E-07 | 0.2 |
| chr2:136008381 | rs309137 | *CXCR4* | T | C | 25.5 | -0.181 | 2.40E-13 | -0.101 | 1.40E-06 | 0.014 |
| chr2:136125862 | rs11688530 | *CXCR4* | G | A | 7.1 | 0.253 | 5.20E-11 | 0.218 | 6.00E-12 | 0.48 |
| chr2:136136760 | rs555647251 | *CXCR4* | C | T | 0.083 | 2.692 | 1.40E-09 | 1.073 | 2.20E-03 | 0.0042 |
| chr2:136143990 | rs10193623 | *CXCR4* | TA | T,TT | 5.9 | -0.207 | 3.00E-07 | -0.191 | 1.60E-08 | 0.76 |
| chr3:37477922 | rs201494641 | *ITGA9* | TTT | T,TT,TTTT | 13.0 | -0.139 | 9.70E-07 | -0.108 | 1.60E-05 | 0.41 |
| chr12:62912352 | rs699585 | *PPM1H* | T | G | 46.1 | -0.123 | 2.80E-10 | -0.127 | 4.80E-14 | 0.88 |
| chr19:1079960 | rs36084354 | *ARHGAP45* | G | A | 8.8 | 0.100 | 1.00E-02 | 0.203 | 3.30E-08 | 0.054 |
| chr19:33264458 | rs12975577 | *CEBPA* | T | C | 49.0 | -0.117 | 8.50E-09 | -0.079 | 1.30E-05 | 0.16 |

**Abbreviations:** Other allele (OA); Effect Allele (EA); Effect allele frequency (EAF).

### Supplementary Table 11

List of antibodies used for phenotyping.

| | Antigen | Fluorochrome | Clone | Manufacturer | Catalog no. | Comment |
|---|---|---|---|---|---|---|
| **Adult blood** | CD34 | PerCP-Cy5.5 | 8G12 | BD | 347222 | Phase I and II |
| | CD34 | PE-CF594 | 563 | BD | 562449 | Phase III |
| | CD45 | APC-H7 | 2D1 | BD | 560178 | Phase I, II and III |
| **Cord Blood** | CD45 | Alexa Fluor 700 | HI30 | BioLegend | 304024 | HSPC markers |
| | CD34 | PE-Cy7 | 581 | BioLegend | 343516 | |
| | CD38 | BV421 | HIT2 | BD Horizon | 562444 | |
| | CD90 | PE | 5E10 | BioLegend | 328110 | |
| | CD45RA | FITC | MEM-56 | Life Technologies | MHCD45RA01 | |
| | CD10 | APC | HI10a | BD Bioscience | 332777 | |
| | CD135 | BV711 | 4G8 | BD Horizon | 563908 | |
| | CD3 | APC-H7 | SK7 | BD Pharmingen | 560176 | Lineage markers |
| | CD4 | BV510 | SK3 | BD Horizon | 562970 | |
| | CD8 | PerCP-Cy5.5 | SK1 | BD Pharmingen | 565310 | |
| | CD14 | BV605 | M5E2 | BD Horizon | 564055 | |
| | CD16 | BV786 | 3G8 | BD Horizon | 563689 | |
| | CD56 | BV650 | NCAM16.2 | BD Horizon | 564058 | |
| | CD19 | PE-CF594 | HIB19 | BD Horizon | 562321 | |
